# Analysis of SARS-CoV-2 synonymous codon usage evolution throughout the COVID-19 pandemic

**DOI:** 10.1101/2021.12.17.472912

**Authors:** Ezequiel G. Mogro, Daniela Bottero, Mauricio J. Lozano

## Abstract

SARS-CoV-2, the seventh coronavirus known to infect humans, can cause severe life-threatening respiratory pathologies. To better understand SARS-CoV-2 evolution, genome-wide analyses have been made, including the general characterization of its codons usage profile. Here we present a bioinformatic analysis of the evolution of SARS-CoV-2 codon usage over time using complete genomes collected since December 2019. Our results show that SARS-CoV-2 codon usage pattern is antagonistic to, and it is getting farther away from that of the human host. Further, a selection of deoptimized codons over time, which was accompanied by a decrease in both the codon adaptation index and the effective number of codons, was observed. All together, these findings suggest that SARS-CoV-2 could be evolving, at least from the perspective of the synonymous codon usage, to become less pathogenic.

**Graphical Abstract:** 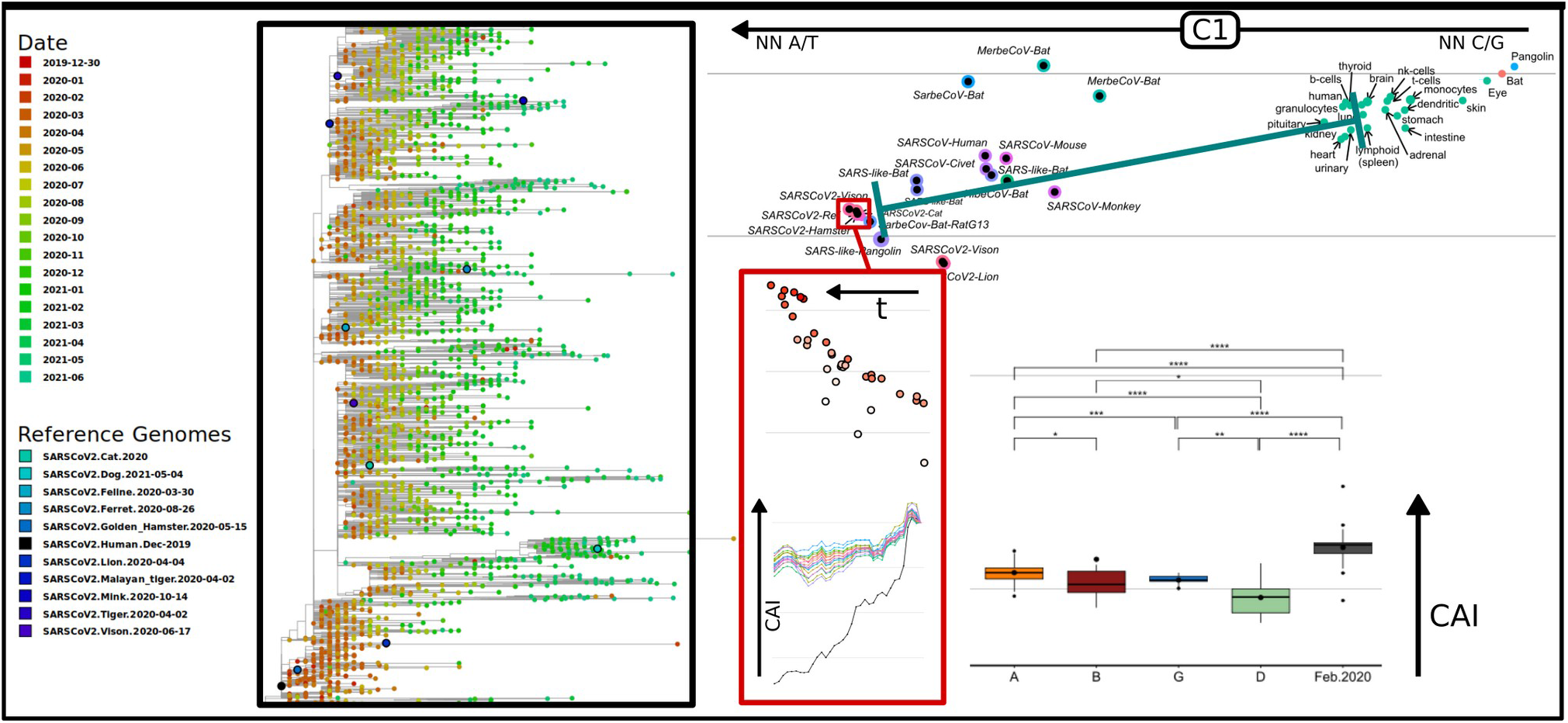

## Introduction

Coronaviruses (CoVs) are members of the *Coronaviridae*, a highly diverse family of enveloped positive-sense single-stranded RNA viruses, further divided in the *Orthocoronavirinae* subfamily, which consists of four genera: *alphacoronavirus*, *betacoronavirus*, *gammacoronavirus* and *deltacoronavirus*. Of these, *alphacoronavirus* and *betacoronavirus* only infect mammalian species, producing respiratory and enteric diseases. Severe acute respiratory syndrome coronavirus 2 (SARS-CoV-2) is the seventh coronavirus known to infect humans; HKU1, NL63, OC43 and 229E viruses cause seasonal respiratory tract infections with usually mild clinical symptoms (common cold), while <u>S</u>evere <u>A</u>cute <u>R</u>espiratory <u>S</u>yndrome coronavirus (SARS-CoV), <u>M</u>iddle <u>E</u>ast <u>R</u>espiratory Syndrome coronavirus (MERS-CoV) and SARS-CoV-2 can cause severe lifethreatening respiratory pathologies and lung injuries [1–3]. Further, SARS-CoV-2 can present several extrapulmonary manifestations that may affect the urinary, cardiovascular, gastrointestinal, hematological, hematopoietic, neurological, or reproductive systems [4–9].

All seven CoVs have similar genomes consisting of a single-stranded RNA molecule of around 27-32 Kb, encoding for a polyprotein, pp1ab (ORF1ab), which is further cleaved into 16 non-structural proteins that are involved in genome transcription and replication; four structural proteins, including spike (S), nucleocapsid (N), envelope (E), and membrane (M) proteins; and a variable number of species-specific accessory proteins. In particular, SARS-CoV-2 reference genome (NCBI Accession NC_045512.2, WHCV) was annotated to possess at least 14 ORFs predicted on the basis of those of known coronaviruses [10, 11], including OR-F1ab, spike (S), envelope (E), membrane (M), nucleocapsid (N) and several accessory proteins (3a, 6, 7a, 7b, 8, and 10). S, the spike glycoprotein, is involved in the attachment to the cell membrane by interacting with the host receptor Angiotensin Converting Enzyme 2 (ACE2), and mediates the internalization of the virus into endosomes (UNIPROT, SPIKE_SARS2) [1]. N, the nucleocapsid phosphoprotein, physically links the +RNA genome to the envelope, interacts with the membrane protein M, and is involved in the RNA packaging and encapsidation [12]. M, is the central organizer of coronavirus assembly, interacting with all other major coronaviral structural proteins, including N, S and E (InterPro, IPR002574). E is an integral membrane protein which forms a Ca^2+^ permeable channel in the endoplasmic reticulum / Golgi apparatus, and is involved in assembly, budding, envelope formation, and pathogenesis. ORF3a, is a pro-apoptosis-inducing protein that localizes to the endoplasmic reticulum (ER)-Golgi compartment and forms homotetrameric potassium, sodium or calcium sensitive ion channels (viroporin) that causes ER-stress in host cells and may modulate virus release (PFAM, PF11289). ORF6 is located in the endoplasmic reticulum, and it has been reported to increase the cellular gene synthesis, induce apoptosis, and to modulate host antiviral responses (InterPro, IPR022736). ORF7a is a viral structural protein involved in the induction of apoptosis that may participate in virus-host interactions (InterPro, IPR014888). ORF7b is a membrane protein necessary for the localization into the Golgi complex (InterPro, IPR021532). ORF8 is a potential pathogenicity factor which evolves rapidly to evade the immune response and facilitate the transmission between hosts (InterPro, IPR022722). Orf10 appears to have no homologous proteins in SARS-CoV and other coronaviruses, and it has been suggested that it may not have a protein coding function (InterPro, IPR044342). Further, a high-confidence protein-coding gene set was verified by ribosome-profiling experiments [13] and comparative genomics [14].

Biological beings share a set of 20 amino acids, eighteen of which can be encoded by more than one synonymous codon. The codon usage frequency is usually not random, and has been related to translation efficiency and accuracy, mutational drift, and other selection pressures [15–22]. In general, viruses only show a slight codon usage bias (CUB), mainly explained by uneven base composition and, hence, by mutation pressure [23]. It has been proposed that a low and non-optimal codon usage allows viruses to adapt to a wider range of hosts with various codon usage preferences [24–26]. In addition, a deficiency in CpG and UpA dinucleotides was observed in most single-stranded RNA and small DNA viruses, probably related to the immunostimulatory properties of unmethylated CpGs [23, 27–30], and to a marked cytosine deamination [31]. In RNA viruses such as SARS-CoV and Ebola Zaire (ZEBOV), mutational pressure was proposed as the most important cause of patterns of codon usage [24, 32–34]. However, it is not completely clear whether this could be generalized, since for Zika virus [25] and MERS-CoV, only a small fraction of the CUB (< 16%) could be explained by mutational pressure [35]. Further, for SARS-CoV-2, different results were reported, indicating from a main role of mutational pressure, to a strict selection pressure [33, 36–38]. In the case of human coronaviruses, including SARS-CoV, MERS-CoV and SARS-CoV-2, several CUB analyses were carried out [26, 33, 43–49, 35–42]. As a result, some general conclusions could be made: first, all of them possessed high AU content and low GC content, with the CpG dinucleotide markedly under-repre - sented, and in the case of SARS-CoV-2, a preferred use of U-ending codons; codon usage bias and codon pair usage were found to be quite different from that of the human host, even when particular tissues such as lung and kidneys were analyzed [50]; high <u>E</u>ffective <u>N</u>umber of <u>C</u>odons (ENC) [51] values were found (although lower than those of other coronaviruses), suggesting a slight codon usage bias; in comparison to other coronaviruses, SARS-CoV, MERS-CoV, and SARS-CoV-2 presented the highest values of the <u>C</u>odon <u>A</u>daptation <u>I</u>ndex (CAI) [52] calculated using human proteins as the reference set, suggesting that these viruses are more adapted to the human host than other coronaviruses that present milder clinical symptoms; a relatively high average CAI value was found for SARS-CoV-2 (approximately 0.7), however, its value was smaller than the average for human genes (approximately 0.8) [44]. Moreover, MERS-CoV and SARS-CoV clustered closer to human genes in correspondence analyses of <u>R</u>elative <u>S</u>ynonymous <u>C</u>odon <u>U</u>sage (RSCU), and presented higher CAI values than SARS-CoV-2, indicating a relatively lower adaptation of SARS-CoV-2 to human cellular systems [37].

The <u>C</u>odon <u>U</u>sage <u>B</u>ias (CUB) of individual proteins from SARS-CoV-2 was also analyzed, and compared with that from other coronaviruses. In the case of Spike (S) protein, the CUB was found to be similar to that of other coronaviruses, with preferential use of A/U ending codons, and partly governed through the mutational pressure (27.35%) and majorly through natural selection and other factors (72.65%). In addition, CAI values for the S-gene indicated a relatively better adaptability in humans, when compared to other mammals [53]. The same bias was observed for ORF1ab, and it was reported that it was more pronounced in SARS-CoV-2 than in SARS-CoV [54]. The mutational status of genes N, S, M, RdRP and S revealed that N, RdRP and S evolve faster than N and M, accumulating amino acid substitution more rapidly and presenting lower ENC values [39, 41].

Further, it has been reported that introducing rare codons within highly expressed genes can affect the translation of other genes, even in a proteome-wide manner, by reducing the availability of the corresponding t-RNAs [55]. The same reasoning is valid in the case of the introduction of highly expressed foreign genes, which can deplete the host cell’s t-RNA pools affecting translation and producing deleterious collateral effects, whether rare or optimized codons are used. Such is the case of viruses, for which a higher similarity of codon usage frequencies was observed for symptomatic compared to asymptomatic hosts [35, 56]. Besides, it has been reported that during SARS-CoV-2 infection, the translation of highly expressed human genes sharing the codon usage of the virus ORFeome appears to be down-regulated [57]. A similar approach was used to identify human genes that could be potentially deregulated due to the codon usage similarities between the host and the viral genes [58].

Although the codon usage pattern of SARS-CoV-2 has been thoroughly described, there are only a few works assessing the adaptation of SARS-CoV-2 codon usage since its transfer to human hosts [40, 43]. It was reported that during the first six months of the COVID-19 pandemic, SARS-CoV-2 average ENC decreased, principally due to C to U mutations (i.e. 47% of all the mutations) that occurred on the 2nd and 3rd codon positions, resulting in a more biased codon usage. Furthermore, the codon usage profile of SARS-CoV-2 seems to have moved away from the human optimal. Interestingly, the CpG and UpA dinucleotides, which are markedly suppressed in many RNA and small DNA viruses, appear to be increasing in the SARS-CoV-2 genome over time, and could result in virus attenuation and decreased pathogenicity [40]. CAI values for most of SARS-CoV-2 coding sequences have also decreased, further suggesting that pathogenicity in humans could be decreasing [43]. In this work we provide an updated analysis of the evolution of the pattern of codon usage of SARS-CoV-2, using a time-series of complete genome sequences collected since December-2019, including the more relevant variants of concern.

## Material and Methods

### Retrieval of genomic sequences

The betacoronavirus (taxid=694002) genome and coding sequences used on this work were downloaded from NCBI Virus Variation Resource [59] (https://www.ncbi.nlm.nih.gov/labs/virus/vssi/#/), filtering for complete genomes (nucleotide completeness = complete) with no ambiguous characters. In the case of reference genomes, a Refseq genome completeness filter was added. SARS-CoV-2 sequences were downloaded using taxid 2697049, and in addition to the previously used filters, sequences were manually selected to be representative of different geographic locations, PANGOLIN [60] lineage classification, and different collection dates (with at least month and year information) from December 2019 to July 2021. Both complete genomes and coding sequences (CDS) were downloaded and grouped every two weeks. In the case of complete SARS-CoV-2 genomes, 2834 sequences were downloaded. CD-HIT server [61] (http://weizhonglab.ucsd.edu/cdhit-web-server/) was used to remove identical sequences, a total of 2725 sequences remained. All the Coding Sequences (CDS) from a total of 209,436 SARS-CoV-2 genomes were downloaded for the time-series correspondence analysis of codon usage frequencies. For the analysis of individual proteins, a homemade script was used to separate the sequences by ORF, using the annotation in the Fasta file headers.

For the analysis of variants of concern, a set of sequences was downloaded from NCBI Virus Variation Resource using taxid 2697049, and in addition to the previous filters, the PANGOLIN lineage prediction filter was used to select for variants: Alpha – B.1.1.7, Beta – B.1.351, Gamma – P.1 and Delta – B.1.617.2 (https://www.cdc.gov/coronavirus/2019-ncov/variants/variant.html).

The transcripts for all the human genes were downloaded from Gencode release 38 (https://www.gencode-genes.org/human/). Highly expressed proteins sets from different human tissues were obtained from the Human protein atlas (https://www.proteinatlas.org/; [62]) and their coding sequences were extracted from the human transcript files using homemade scripts. The horseshoe bat (*Rhinolophus ferrumequinum)* and chinese pangolin (*Manis pentadactyla*) genome coding sequences were downloaded from NCBI genomes (Assembly accessions GCF_004115265.1 and GCF_014570555.1 respectively). The complete list of SARS-CoV-2 sequences used is on Table S1.

### Phylogenetic analysis

To construct a phylogenetic tree, a Multiple Sequence Alignment (MSA) of the complete genomes of reference beta-coronavirus (Beta-CoVs) and SARS-CoV-2 was obtained using MAFFT [63] (https://mafft.cbr-c.jp/alignment/software/closelyrelatedviralgenomes.html; MAFFT v7). First a MSA of all the reference sequences was obtained using the Iterative refinement method E-INS-i and default parameters. In a second step all the SARS-CoV-2 genome sequences were added to the previous MSA following the recommended instructions for full-length MSA of closely-related viral genomes (https://mafft.cbrc.jp/alignment/server/add_fragments.html?frommanualnov6; Default parameters were used).

A maximum likelihood phylogenetic tree was constructed for the first MSA using IQtree [64] (Galaxy Australia, Version 2.1.2). The Best-fit model according to BIC (GTR+F+I+G4) and ultrafast bootstrap were used as evolutionary model and branch support (1000 bootstrap), respectively. For the second alignment, containing all 2725 SARS-CoV-2 sequences, a maximum likelihood phylogenetic tree was constructed with Fasttree [65] (v2.1.10, at Galaxy Australia) using the GTR model with four gamma rate categories.

### Synonymous codon usage analysis

The codon usage analysis was done using homemade Bash and R scripts. First, codon counts, synonymous codon usage frequencies, and average codon usage frequencies for each set of genes were calculated using either Gary Olsen codon usage scripts (https://www.life.illinois.edu/gary/programs/codon_usage.html), or the coRdon R package [66]. Relative synonymous codon usage frequencies calculated with G. Olsen codon usage scripts are similar to the commonly used RSCU, but normalized to 1 (i.e. the maximum value for each codon is 1). Averaged codon usage frequencies were calculated from the summed codon counts of all the genes in a multifasta sequence file. Codon counts for ENC, and CAI determination were calculated with coRdon [66], using the concatenated ORFs (or equivalently, the codon count for each ORF were summed) for each SARS-CoV-2 genome. CAI was calculated using a homemade R script following Sharp’s CAI equations [52]. Highly expressed proteins in different human tissues were used as reference sets (https://www.proteinatlas.org/; [62]). For *W* determination, all the CDSs in the reference set were concatenated (or equivalently, the codon counts for each CDS were summed) and pseudo-sums of 0,01 were added to 0 frequency codons (*Wconcat*). Alternativelly, a *Wi* was calculated for each gene, and the average *W* (*Wavg*) was used for CAI calculation. In the case of SARS-CoV-2 genomes, all the CDS were concatenated except OR-F1a, which overlaps with ORF1ab.

## Results

### Phylogeny of SARS-CoV-2

In order to analyze the divergence of SARS-CoV-2 since its onset in 2019, a maximum likelihood phylogenetic analysis of complete genomes was made. In a first instance, IQTree was used to construct a phylogenetic tree including reference Beta-CoVs from the *Merbecovirus*, *Nobecovirus*, *Embecovirus*, *and Sarbe-covirus* subgenuses. As previously reported, SARS-CoV-2 clustered together with SARS-CoV and other SARS-related coronaviruses found mainly in bats, within the *Sarbecovirus* subgenus (Fig. 1) [11, 29, 43, 67, 68]. Then, a maximum likelihood phylogenetic tree including a subset of NCBI public SARS-CoV-2 sequences representing the most relevant SARS-CoV-2 variants sampled by date and geographic region, was constructed using FastTree (Fig. 2). It can be seen that most SARS-CoV-2 sequences clustered by variant rather than by geographic location, although some variants also clustered by region. In addition, in accordance to previous reports [69, 70], newly sequenced isolates presented more divergence with respect to the Wuhan-2019 reference sequence (i.e. 1 × 10^-3^ substitutions per site, or about 30 substitutions per genome. Fig. 2), suggesting that they could be used to monitor the evolution of SARS-CoV-2 codon usage pattern in humans over time. However, since most of the registered single nucleotide polymorphisms (SNPs) produced non-synonymous mutations (i.e. a ratio of non-synonymous to synonymous substitutions of 1.88, and an 80% of the recurrent mutations) [69], we expected to observe only a small variation in the codon usage profile. Similar results were observed using the Nextstrain web portal, with selected data from GISAID and coloring by emerging lineage and date respectively (Fig. S1, https://nextstrain.org/ncov/gisaid/global, accessed 2021-09-22) [71].

**Figure 1.**
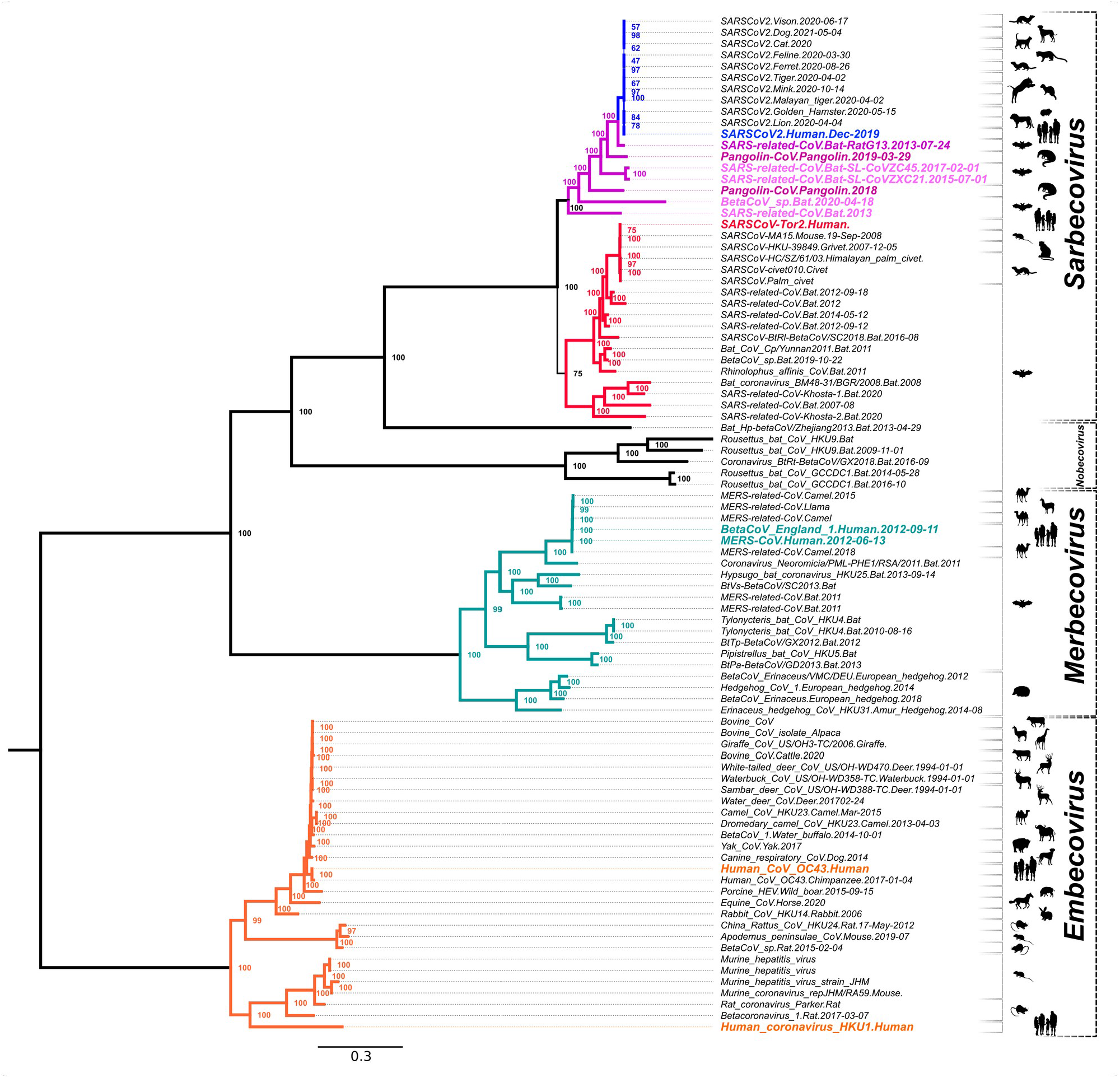
Maximum Likelihood Phylogenetic tree of Betacoronavirus. Maximum Likelihood phylogenetic tree constructed using full genomes of betacoronaviruses belonging to the subgenuses *Sarbecovirus*, *Nobecovirus*, *Merbecovirus* and *Embecovirus*. Genomic sequences were downloaded from NCBI Virus database, aligned with MAFFT and a ML phylogenetic tree was constructed with IQTree. The most relevant Beta-CoV isolates are highlighted with different colors. Blue: human SARS-CoV-2 Wuhan 2019 isolate. Purple: Pangolin-CoVs and Bat RaTG13, SL-CoVZC45 and SL-CoVZXC21 isolates. Red: human SARS-CoV Tor2 isolate. Teal: human MERS-CoV isolates. Orange: Human CoVs from the *Embecovirus* subgenus. Numbers represent the bootstrap support for each node.

**Figure 2.**
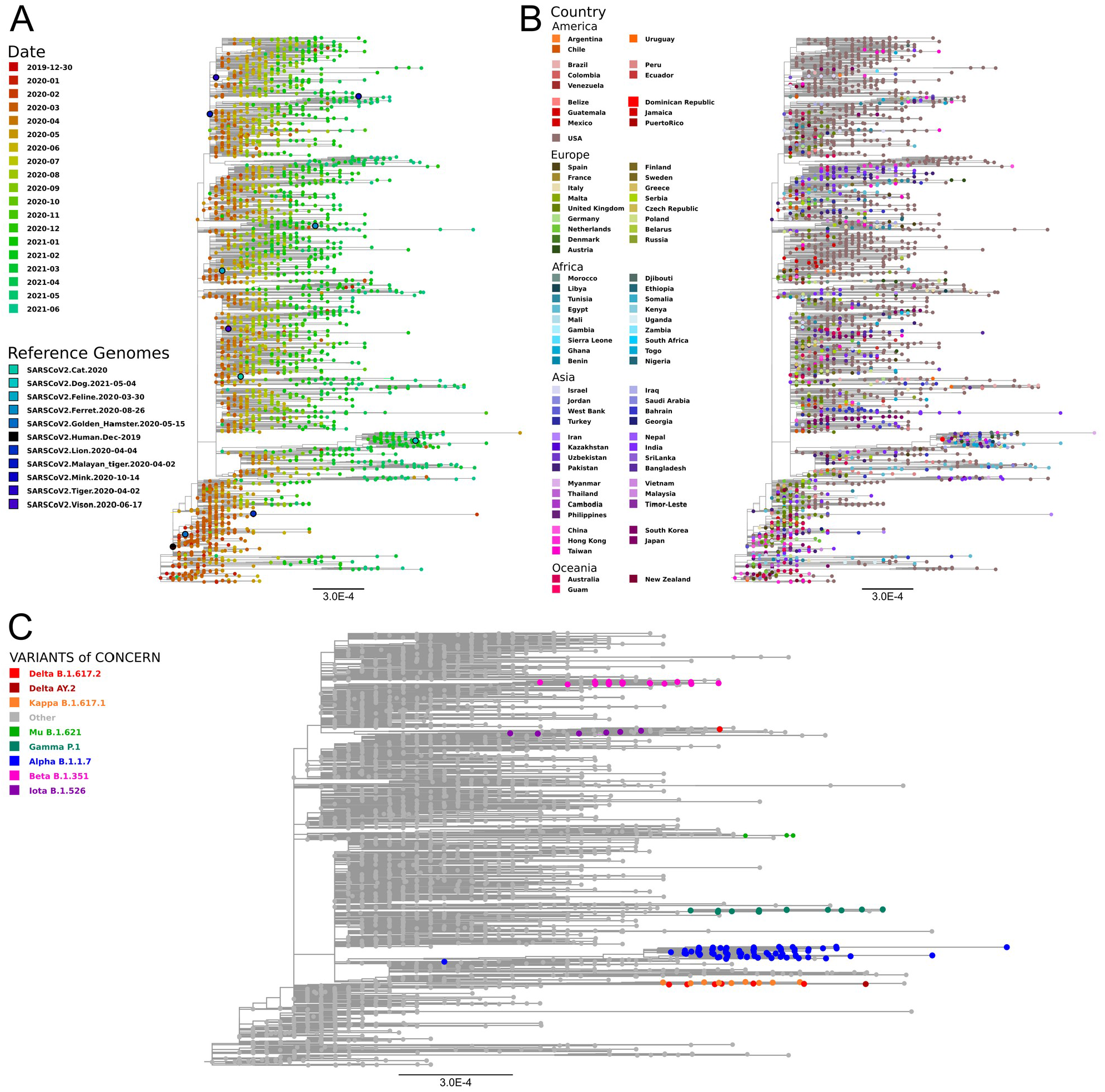
SARS-CoV-2 Maximum Likelihood phylogenetic tree constructed with Fasttree using full genomes of isolates with different collection dates and geographic origins. Genomic sequences were downloaded from NCBI virus database, selecting isolates from different geographic regions, SARS-CoV-2 variants and collection dates. Nucleotide sequences were aligned with MAFFT and a ML phylogenetic tree was constructed with Fasttree. A) Leaves colored by time. Different colors from red (Dic-2020) to green (Jun-2021). Reference sequences correspond to SARS-CoV-2 isolated from human (Wuhan isolate 2019) and different animals. B). Leaves colored by geographic region. C) Leaves colored by SARS-CoV-2 variant. Only Alpha, Beta Gamma, Delta, Mu, Iota and Kappa are shown.

### Codon usage analysis

In order to evaluate the relative synonymous codon usage frequencies either belonging to beta-CoVs or humans we performed a <u>C</u>orrespondence <u>A</u>nalysis (CA), using as input the <u>A</u>verage <u>C</u>odon <u>U</u>sage <u>F</u>requencies (ACUF) for different gene sets. We included human tissues that can be infected by SARS-CoV-2 including lungs, heart, kidneys, and brain among others. Human transcripts were extracted from Gencode R38, and highly expressed gene sets were created for each organ using tissue specific proteomes from the human protein atlas [62]. As it can be seen on the CA shown in Figure 3, SARS-CoV-2 isolates clustered together with SARS-related coronaviruses including Bat and Pangolin CoVs, and are quite far from the human tissues, and also from the genomes of Bat and Pangolin. The phylogenetic and codon usage analysis results support that SARS-CoV-Bat-RatG13 is the most closely related to SARS-CoV-2 reference strain. Besides, SARS-CoV-2 is more distant from human tissues that SARS-CoVs or human MERS-CoVs. It is also clear that the main axis (C1) of the CA, accumulating 73% of the variation, corresponds to the difference in the frequency of AT or C-G terminated codons, while the secondary axis (C2, 15% of the variation) is mainly defined by the differences in the Tyr (TAT vs TAC), Leu (TTG vs TTA), Arg (CGT, CGA, CGC, and CGG vs AGA), Asp (GAT vs GAC), Ser (TCG, AGC and AGT vs TCA), Glu (GAG vs GAA), Phe (TTT vs TTC), Lys (AAG vs AAA), Gln (CAG vs CAA), Asn (AAT vs AAC), Ile (ATT, ATA vs ATC), Thr (ACG, ACC and ACT vs ACA) and Pro (CCG, CCC vs CCA, CCT) codon choice (Fig. 3. Inner light gray shaded plot). Next, in order to analyze the time variation of ACUF in SARS-CoV-2, sequences were clustered by date in bins of 15 days, and a correspondence analysis was done (Fig. 4). The first conclusion that can be drawn from these analyses is that in the time that SARS-CoV-2 has been infecting human hosts, only a slight variation in the average codon usage has taken place. It also appears, that opposite to what it would be required for a higher expression of SARS-CoV-2 proteins, the codon usage tends to be slightly more distant (C1) from that of the human host in the most recently collected samples. Since new SARS-CoV-2 variants appeared over time and gained importance, correspondence analysis of sequence sets clustered by variant (i.e. as determined by PANGOLIN on NCBI Virus Variation Resource, [59]), or by variant and date, were made, including the new sequence sets to the previous analysis (i.e. all the previous sequences plus the new ones were used in the CA. Fig. 5). Remarkably, Delta (D) and Gamma (G) variants could be clearly separated based on the CA of ACUF, while Beta (B) and Alpha (A) were overlapped with the sequence sets corresponding to the time-series. However, no clear time dependence was observed (Fig. S2), possibly as an effect of the brief time since the appearance of these variants. To try to elucidate which codons presented more variation over time for the selected variants, the difference of ACUF with that of the month of January 2020 was calculated for each codon and time (Figures 6, S3 and S4). As it was previously reported [37], an overall antagonistic codon usage pattern to human t-RNA isoacceptors was found (Table S2), with a total of 11 amino acids encoded by antagonistic codons, including all the amino acids encoded by two codons. During the COVID-19 pandemic, most of the U ending codons increased their frequency (AAT, ATT, ACT, CAT, CCT, CGT, CTT, GGT, GTT, TTT, TCT, TGT) including some of the antagonistic codons preferred by SARS-CoV-2 (AAT, GGT, CAT, GTT, TTT, TGT). Only a few of the SARS-CoV-2 antagonistic codons (CAA, TAT) got closer to human frequencies. Also, the results show that several codons presented a higher variation, being AAT, AAC (Asn); ACT, ACC (Thr); AGA, AGG, CGA, CGT (Arg); ATA, ATC, ATT (Ile); CAA, CAG (Gln); CAT, CAC (His); CCT, CCA (Pro); CTT, TTA, TTG (Leu); GAC, GAT (Glu); GAT, GAC (Asp); GCT (Ala); GGA, GGC, GGT (Gly); GTT (Val); TAT, TAC (Tyr); TCC, TCT (Ser); TGC, TGT (Cys); and TTT, TTC (Phe) the most representative. In addition, some codons presented more variation between the new variants (i.e. AAA/AAG-Lys, AAC/AAT-Asn, AGG/CGA/CGT-Arg, ATA/ATC/ATT-Ile, CAT/CAT-His, CCA/CCC/CCG-Pro, GAC/GAC-Asp, GCT-Ala, TAC/TAT-Tyr, TGC/TGT-Cys, TTT/TTC-Phe), while others presented a steady increase (AAG, CCT, CGA, GCG, GCC, GTT, GAA, TAC, TTG) or decrease (AAA, AGA, GCT, GTC, TAT, TTC) on their ACUF (Fig. S4). The main codons contributing to differences in SARS-CoV-2 variants were AAA-Lys (A<D/G<B), AAG-Lys (B<G/D<A), ACA-Thr (D<A/B/<G), ACC (B/G < A/D), ACG (A/B < D/G), AGG (G<D<A<B), CCA (G<D<B<A), CCC (A<B<D<G), CCG (A<B/D<G), CGA (D/B<A<G), CGG (B/ G/A<D), CTG (D<G/B/A), GAA (ABGD < average), GAG (ABDG > average), GCA (BDG < A), GTG (ABD < G), TAC (G<D/B<A), TAT (A<B/D<G), TCC (A<G<B/D), TGC (A/D < G/B), TGT (G/D<A/B), and TTG (ABGD < average). Interestingly, some of the codons which presented an increase in the ACUF (although slight) contained CG or TA dinucleotides (ATA, ATT, CAT, GCG, and CGA), and it was speculated that an increase of CpG and UpA dinucleotides could reduce SARS-CoV-2 pathogenicity [40].

**Figure 3.**
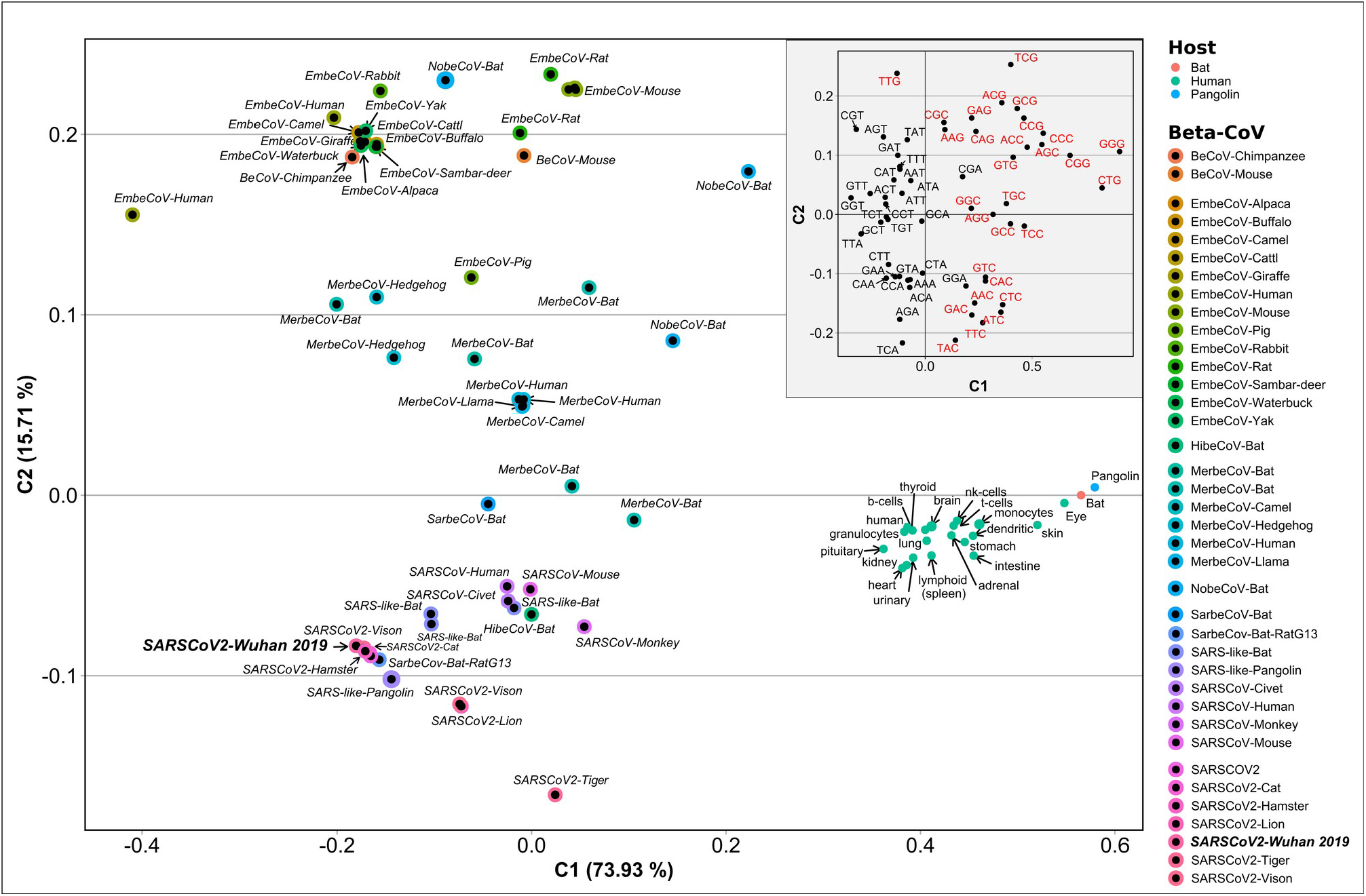
Correspondence Analysis of Average Codon Usage Frequencies (ACUF) for Betacoronavirus and their hosts: Human, Bat, and Pangolin. Coding sequences for SARS-CoV-2, Human, Bat, and Pangolin genes were downloaded from NCBI and ACUFs were calculated and used in a Correspondence Analysis as described in the Material and Methods section. The first two components representing 89 % of the total inertia are shown. The inner plot (light gray shading) corresponds to the column variables (codons). Red: Codons with C or G in the third position. Black: Codons with A or T in the third position.

**Figure 4.**
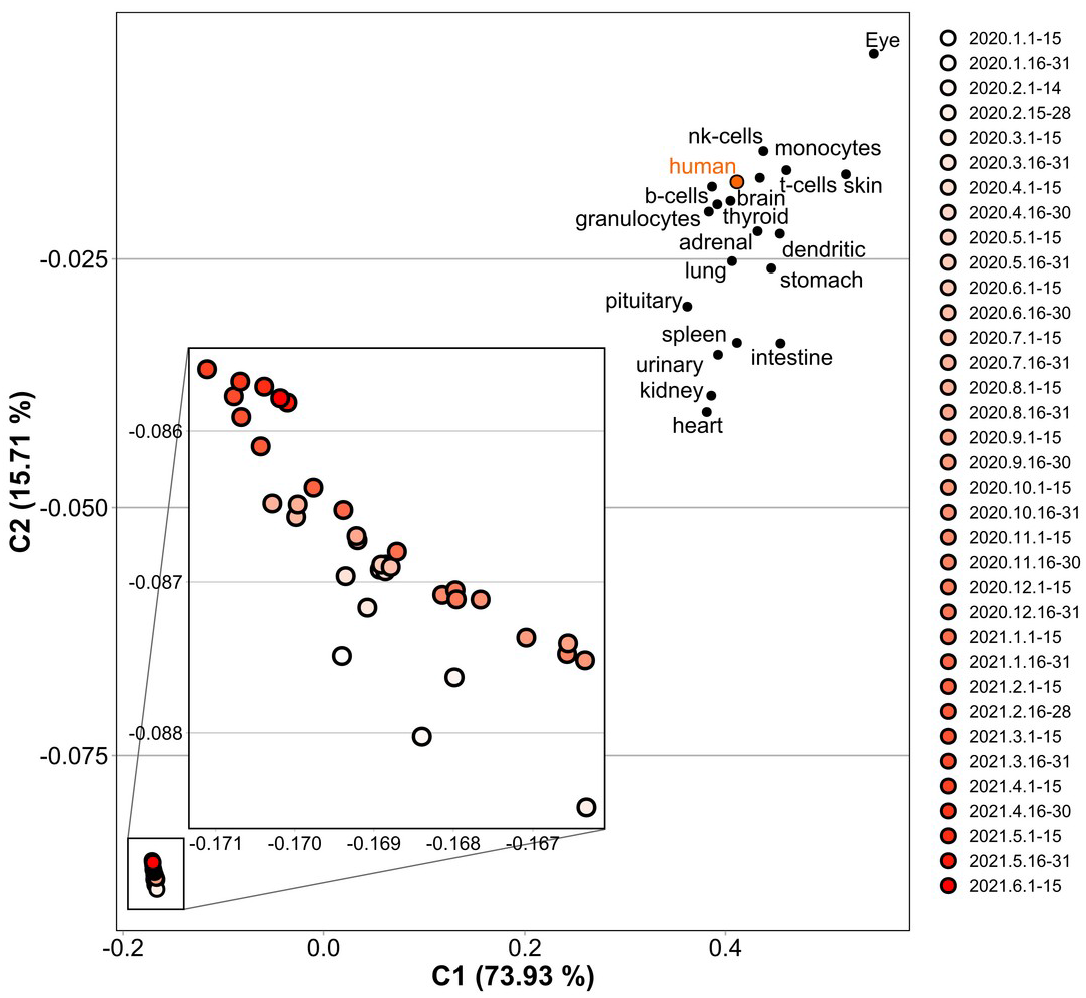
Correspondence Analysis of Average Codon Usage Frequencies (ACUF) for SARS-CoV-2 time-series and human tissues. Coding sequences for SARS-CoV-2 and human genes were downloaded from NCBI and ACUFs were calculated and used in a Correspondence Analysis as described in the Material and Methods section. On this figure, only the points corresponding to the concatenated SARS-CoV-2 coding sequences binned by date, and human genes with elevated expression in different tissues are shown. The Inner plot shows an amplification of the SARS-CoV-2 region of the graph. Colors represent the collection date, from Jan-2020 to Jun-2021.

**Figure 5.**
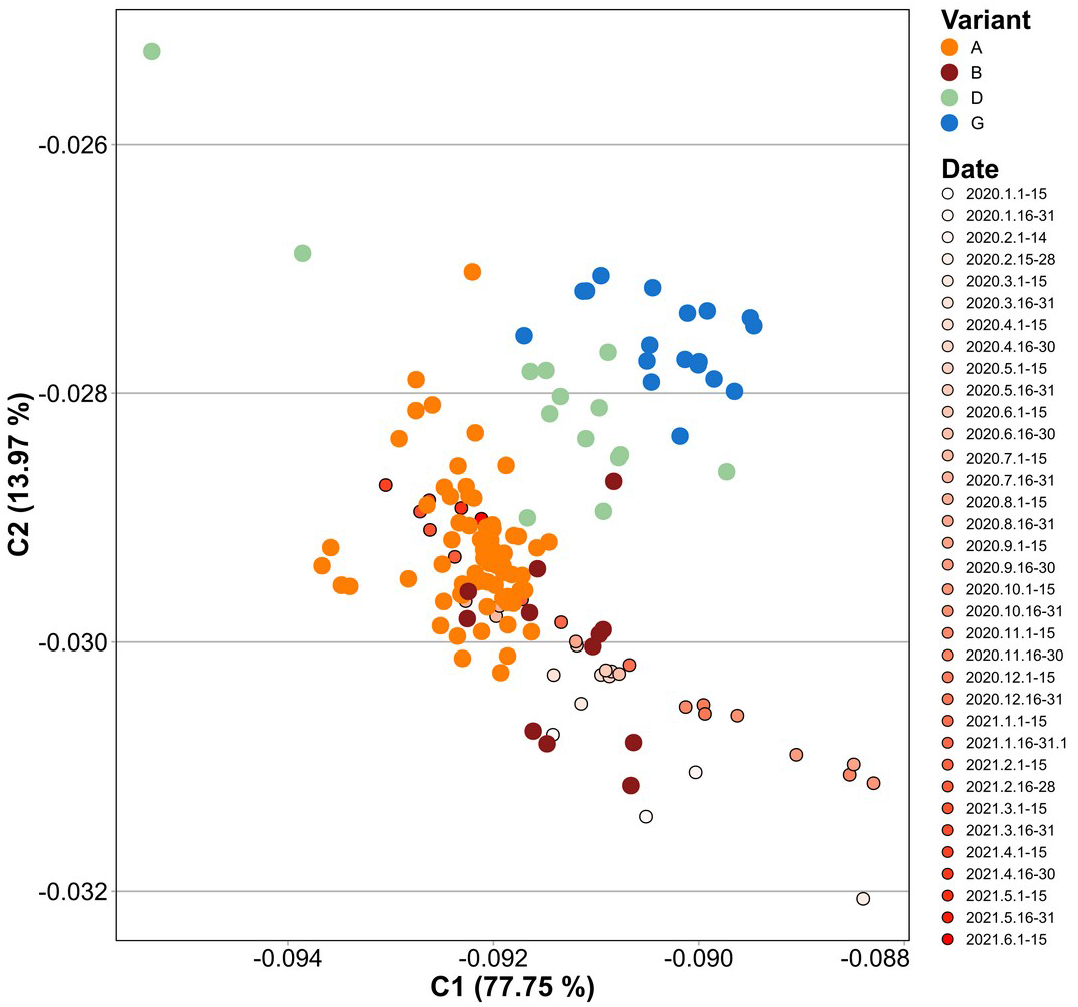
Correspondence Analysis of Average Codon Usage Frequencies (ACUF) for SARS-CoV-2 time-series and selected variants of interest. Coding sequences for SARS-CoV-2 time-series, and for a manual selection of genomes representing Alpha (A), Beta (B), Gamma (G), and Delta (D) variants were downloaded, their ACUFs calculated, and a correspondence analysis was performed as described in the Material and Methods section. The first two components representing 91 % of the total inertia are shown.

**Figure 6.**
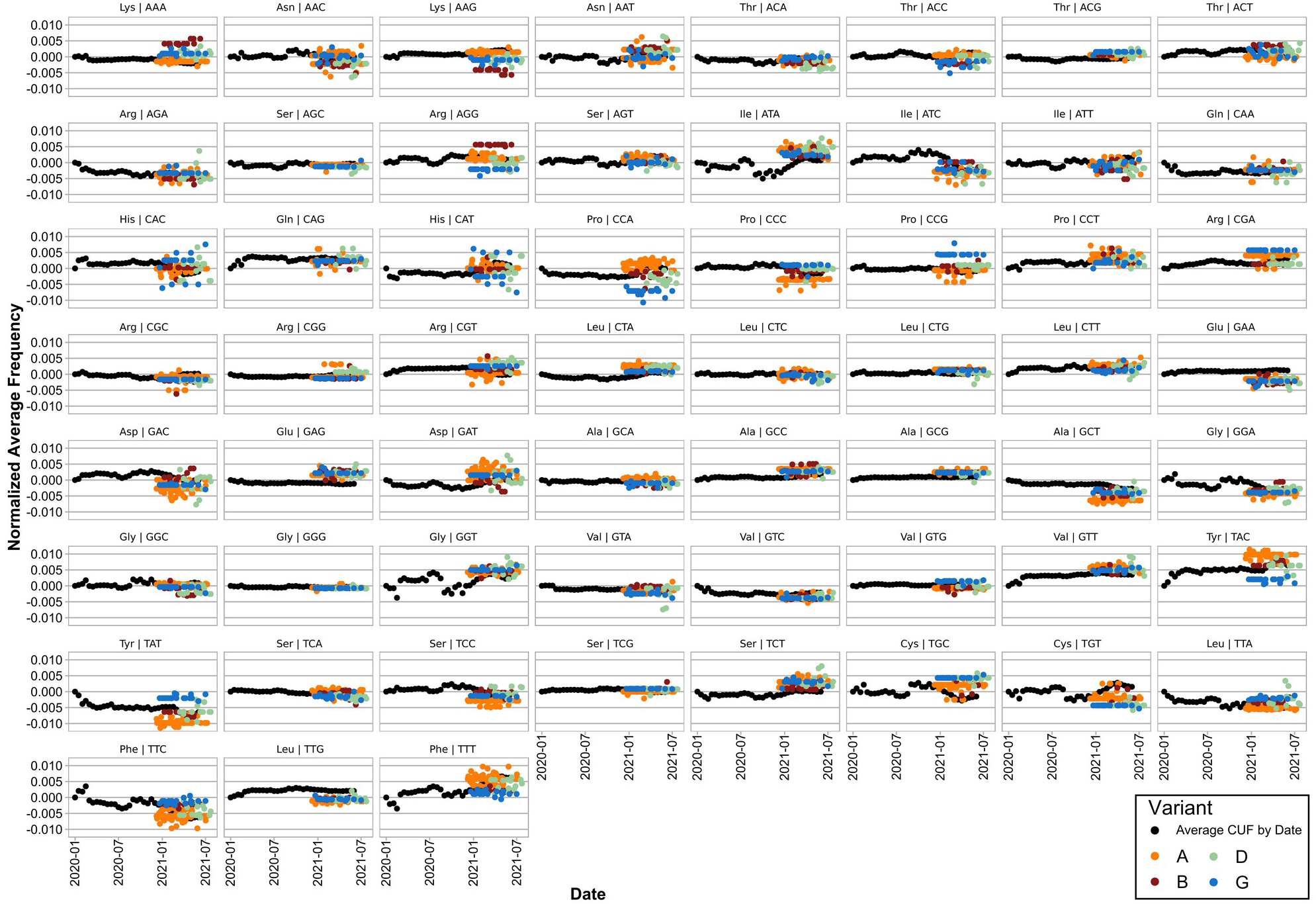
Evolution of the Average Codon Usage Frequency (ACUF) for each codon over time. ACUFs were calculated for concatenated SARS-CoV-2 genes grouped by fortnight, and normalized by subtracting the values registered for the first half of January 2020. Black points represent the ACUF for SARS-CoV-2 isolates from the time series dataset. Color points correspond to selected SARS-CoV-2 variants of interest: A (Alpha) orange, B (Beta) dark red, D (Delta) light green, and G (Gamma) blue.

Next, SARS-CoV-2 coding sequences were clustered by open reading frame (ORF), averaged by date, and a CA of their ACUF was performed, which showed a clear separation by ORF (Fig. 7). Remarkably, S, N, M, ORF3a, ORF7a and ORF8 ORFs were located nearest to the human genes in the CA, suggesting that they could be more adapted for high expression.

**Figure 7.**
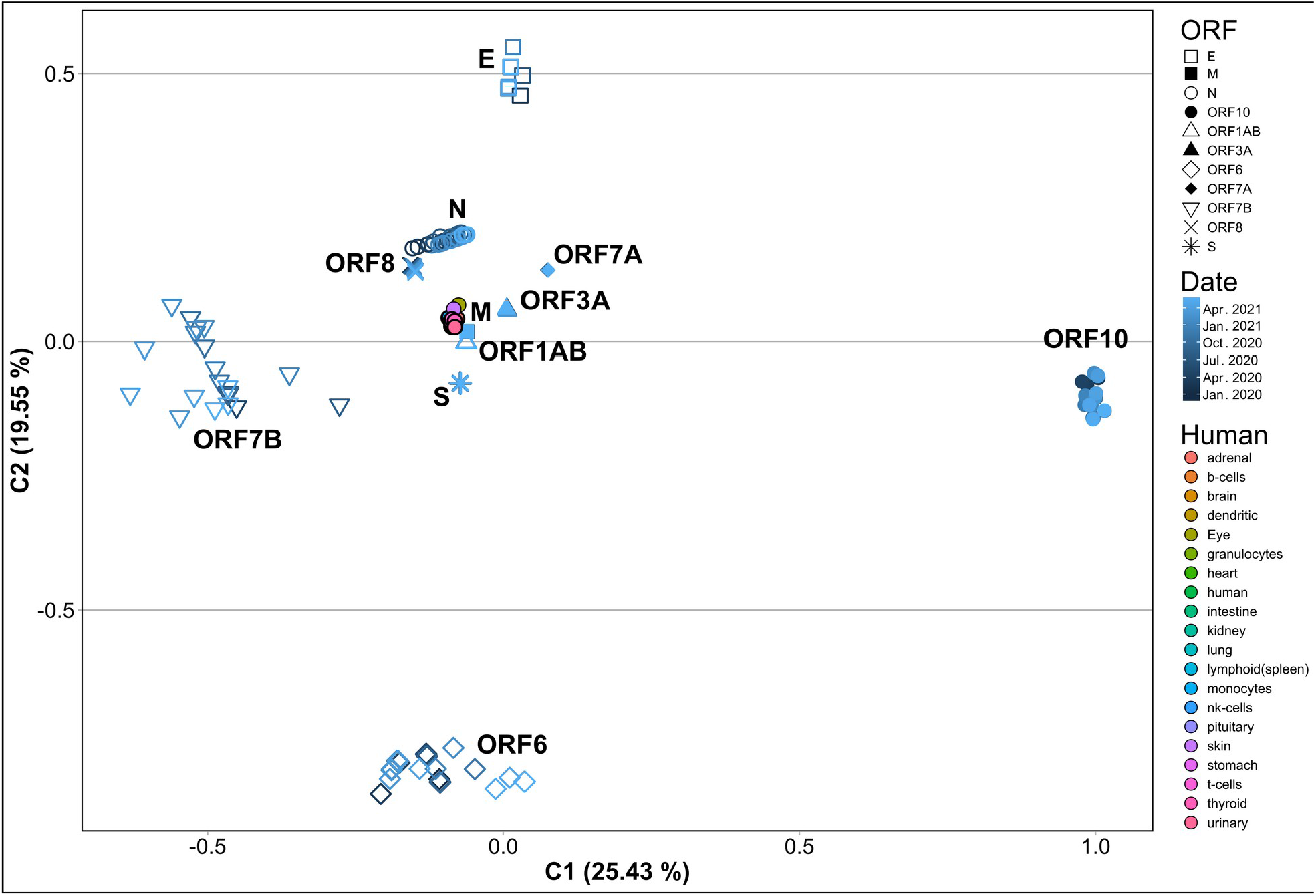
Correspondence Analysis of ACUF for each SARS-CoV-2 ORF from the time-series dataset averaged by month. The coding sequences corresponding to ORF1ab, S, M, N, E, ORF3a, ORF6, ORF7a, ORF7b, ORF8, and ORF10 were extracted from the SARS-CoV-2 time-series dataset, their ACUF were calculated, averaged by month and a CA was performed. Shapes indicate the different ORFs. Dark to light blue colors indicate dates from Jan-2020 to Jun-2021. Human: colors indicate the ACUF for genes with elevated expression in different tissues.

Further, the ACUF for some proteins presented slight or no time dependence at all (i.e. S, M, ORF1ab, ORF3a, ORF7a and ORF8), while other presented considerable variation (i.e. ORF10, ORF6, E and N) with ORF7b presenting the higher variation (Fig. 7 and Fig. S5). In particular, N, ORF3a and ORF8 appear to be getting closer to the human genes, while ORF1ab is getting apart (Fig. S5). Finally, a CA of Codon <u>U</u>sage <u>F</u>requencies (CUF) calculated for sequence sets clustered by ORF, and chosen to represent different dates, geographic regions and variants (Fig. 8 and Fig. S6), was made. In this case, only a slight variation in CUF was observed, with most ORFs clustering together, and only showing an apparent separation by SARS-CoV-2 variant in the case of ORF 8, N, ORF1ab and S (Fig. 8 and Fig. S6). Also for some isolates of the Alpha and Delta variants a greater variation in the CUF for ORFs ORF6, ORF7b, ORF8 and ORF10 was observed.

**Figure 8.**
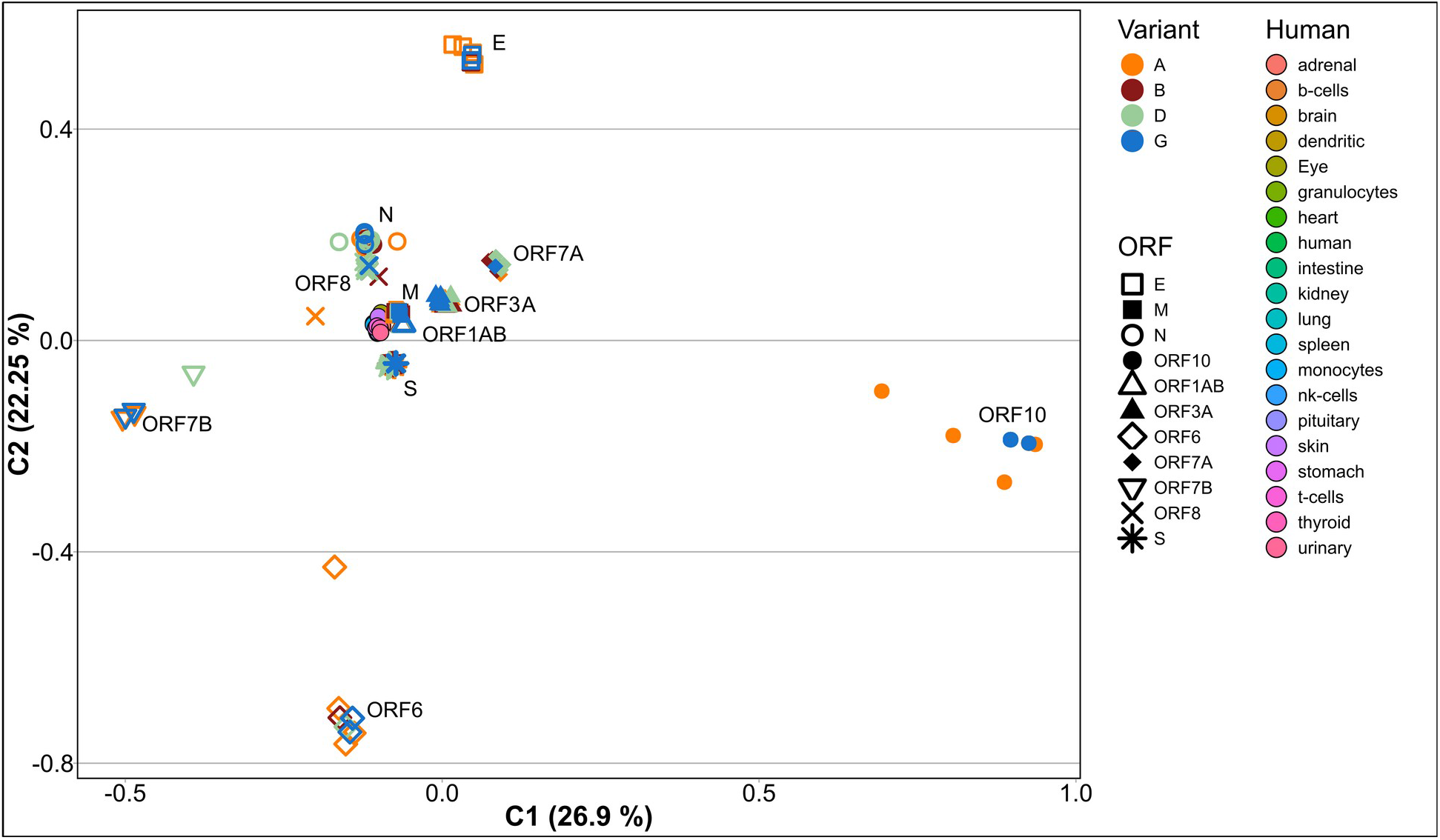
Correspondence Analysis of Codon Usage Frequencies for each SARS-CoV-2 ORF from Alpha, Beta, Gamma and Delta variants. The coding sequences corresponding to ORF1ab, S, M, N, E, ORF3a, ORF6, ORF7a, ORF7b, ORF8, and ORF10 were extracted from the genomes of SARS-CoV-2 Alpha (A), Beta (B), Gamma (G), and Delta (D) variants, their codon usage frequencies were calculated and a CA was performed. Shapes indicate the different ORFs. Human: colors indicate the ACUF for genes with elevated expression in different tissues.

### Effective number of codons and Codon adaptation index

The variation of the <u>E</u>ffective <u>N</u>umber of <u>C</u>odons (ENC) with time was analyzed. ENC is a measure of the degree of codon usage bias in a gene, its values are between 20 and 61, with values near 20 indicating an extreme bias, and higher values, approaching 61, indicating that the codons are randomly used. It has been previously reported that during the first four months of evolution of SARS-CoV-2 in humans, the ENC value was decreasing (i.e. more biased codon usage) with time [43], so an analysis of the variation of ENC with time for concatenated SARS-CoV-2 coding sequences collected from January 2020 to July 2021 was made. First, we evaluated the ENC values for different human tissues, all of which presented almost no codon bias (i.e. high ENC values, above 50, both for values calculated with the concatenated sequences, or for the average of all genes in a determined tissue, Fig. S7 A and B respectively), with Eye, Kidney and Skin presenting the lower values. In contrast, concatenated SARS-CoV-2 genes presented ENC values close to 45.45, indicating a slight codon usage bias. As shown in figure S8, during the 2020 and the first months of 2021, ENC only presented a slight variation (i.e. less than 0.05 units). In the first two months, an apparent increase of ENC was observed, however, these months presented the smaller number of analyzed genomes (approximately 180 each), while thereafter at least 2,000 genomes per month were included. A comparison of the calculated ENC values with those of February 2020 revealed significantly slower values, and a steady decrease until December 2020. However during 2021, ENC increased again, reaching values close to those at the beginning of the COVID-19 pandemic. This was not expected, since an adaptation to the host generally implies a decrease in ENC, and a more biased codon usage. The observed results could be related to the emergence and propagation of novel SARS-CoV-2 variants presenting higher average ENC values. To further test that hypothesis, the ENC values for the concatenated genes were calculated for Alpha, Beta, Delta and Gamma SARS-CoV-2 variants (Fig. S9). Alpha and Beta variants have shown slightly decreased ENC, while Gamma presented a significantly higher ENC when compared to those of February 2020 isolates. The observed variation of ENC over time (Fig. S8) could be related to the higher relative abundances of Alpha, Gamma and Delta variants (i.e. the proportion of those variants in the analyzed genomes, for each time sampled), which present the lowest and higher ENC values respectively (Fig. S9).

ENC values were also analyzed for every SARS-CoV-2 ORF (Fig. S10), and compared between variants. In that case, a statistically significant difference in the ENC values calculated for each ORF between the analyzed variants was observed. The Alpha variant, presented a significantly smaller ENC for ORF1ab and a higher one for S, N, and ORF8. Beta, Gamma and Delta presented a higher ENC for ORF3a. In the case of the Delta variant, a higher ENC was observed for S, while a smaller value was observed for N and M. For the Beta variant, a remarkably higher ENC value was observed for the E protein. Finally, in the case of the Gamma variant smaller ENC values for S and N proteins, and a higher ENC value for ORF8 were observed.

In relation to the <u>C</u>odon <u>A</u>daptation Index (CAI), Huang *et al*. [43] also reported a decrease over time during the first four months of the COVID-19 pandemic. Here we extended that analysis to 18 months, and used highly expressed human proteins in different tissues as reference sets for CAI calculation. The reference sets were extracted from transcripts annotated in Gencode R38 (https://www.gencodegenes.org/human/) using human protein atlas expression profiles (https://www.proteinatlas.org/) [62] to select for highly expressed proteins in lungs, spleen, stomach, kidney, skin, heart, brain, eye, intestine, urinary bladder, thyroid, adrenal and pituitary glands, and different immunity cells like B, T, natural killer, dendritic cells, monocytes, and granulocytes. The relative adaptiveness (*W*) for the reference sets was obtained in two different ways, either by calculating and averaging *Wi* for every gene in the reference set (*Wavg*), or directly by calculating *W* for the concatenated genes (*Wconcat*). In addition, *W* for human ribosomal proteins was extracted from Lei *et al*. [72] (*WLei*). Next, average CAI values were calculated for SARS-CoV-2 isolates collected on the first half of February 2020 (2020/02/01-14). As can be seen in figure S11, depending on the method used for *W* calculation the results varied (mainly in their magnitude) with CAIs calculated with *Wconcat* presenting higher values (Fig. S11.B1). In both cases, granulocytes, kidneys, heart and the pituitary and thyroid glands presented the higher CAI values, which might indicate a higher expression of SARS-CoV-2 proteins in those tissues. In contrast, skin, stomach, intestine, dendritic cells, monocytes and eyes presented the lowest CAI values. Also, SARS-CoV-2 average CAI values were in general smaller than the average CAI for human genes, presenting the smaller difference in the granulocytes, kidneys, heart, and the pituitary and thyroid glands (Fig. S11.A2. And B2.). Moreover, when CAI was calculated using *Wconcat*, SARS-CoV-2 presented the highest CAI values in the heart and pituitary gland. The biggest difference was found on skin, eye, intestine, dendritic cells, and monocytes, which could indicate a lower expression of SARS-CoV-2 proteins in those tissues.

Then, CAI was calculated for time-series of SARS-CoV-2 isolates and averaged by date. Using pairwise Wilcoxon rank-sum test to compare each time with the first half of February 2020, significant differences could be found that support a variation in the average CAI for SARS-CoV-2 over time (Fig. 9 and Fig. S12). In general the behavior of CAI was similar for all tissues, presenting an increase in February, a steady decrease until July-2020, and another small increase followed by a decrease presenting spikes of higher CAI. The major difference between tissues was observed between July-2020 and March-2021. In the case of CAI values calculated using *Wconcat* (Fig. 9.B.) the difference was bigger, with some tissues presenting higher (e.g., pituitary gland and urinary bladder) or smaller (e.g., eye and skin) CAI values. In the first half of 2021, CAI values for eye and skin presented a more pronounced increase than the rest of the analyzed tissues. In addition, CAI values calculated for SARS-CoV-2 with *WLei* presented a constant and more pronounced decrease over time.

**Figure 9.**
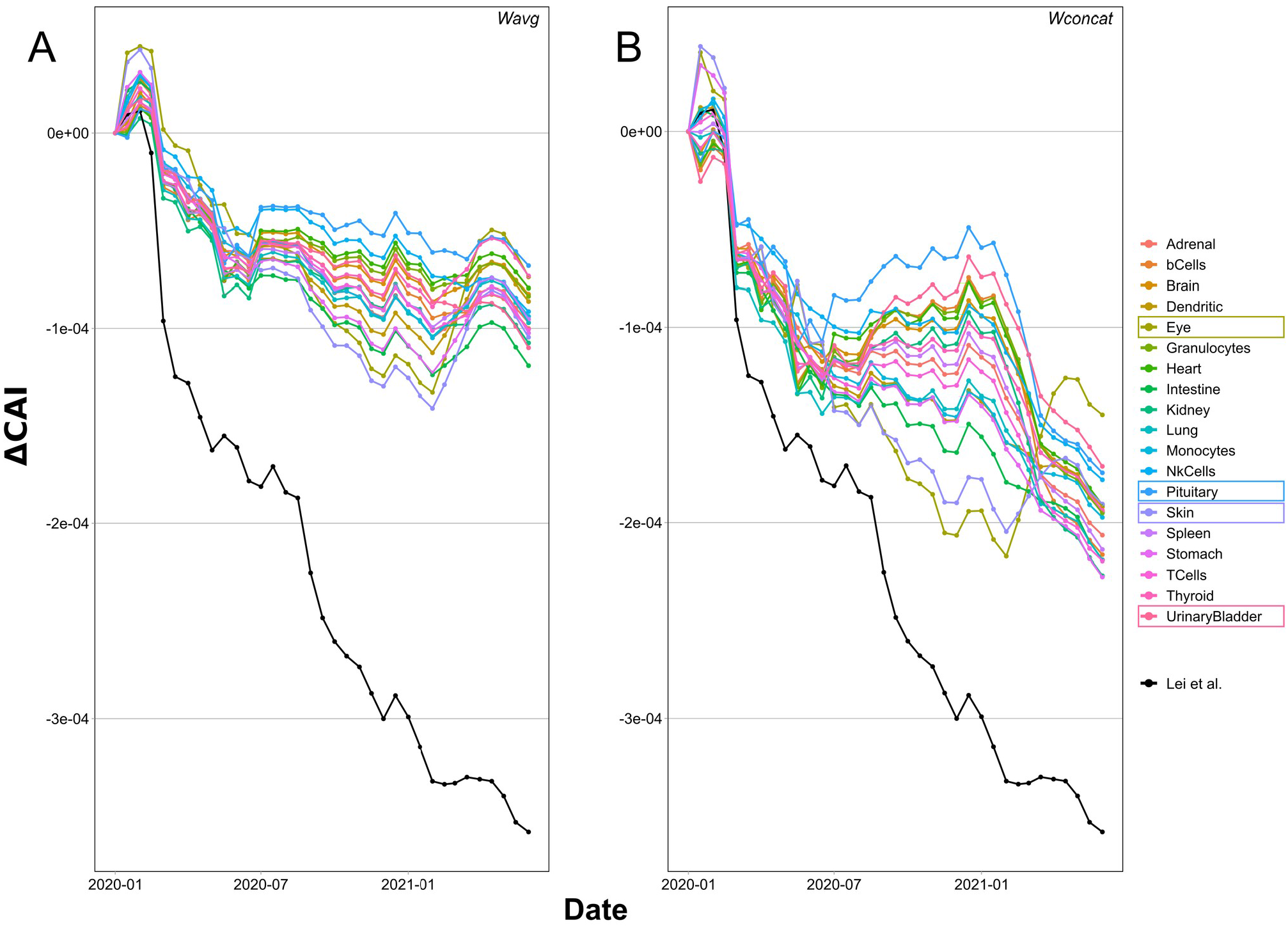
Evolution of the Codon Adaptation Index (CAI) over time for SARS-CoV-2 calculated in reference to different human tissues. CAI was calculated for the concatenated SARS-CoV-2 genes using elevated proteins on each human tissue as reference set, and averaged by month. The figure shows the difference of CAI for each month with the values calculated for Jan-2020 (ΔCAI). Human highly expressed proteins for each tissue were obtained from the human protein atlas. A) CAI values obtained using *Wavg*. B) CAI values obtained using *Wconcat*. The black line corresponds to CAI values calculated using *WLei*.

In order to determine if this behavior was related to the spread of new SARS-Cov-2 variants, CAI was determined for the Alpha (B.1.1.7), Beta (B.1.351), Gamma (P.1), and Delta (B.1.617.2) variants (Fig 10.A.). In this case, *WLei* was used. The results show that all the variants presented lower CAI values than the SARS-CoV-2 isolates collected in the first half of February 2020, with Delta having the lowest CAI. Finally, an analysis of CAI for every ORF and variant was made (Fig. 10.B.). In accordance with the results of the correspondence analysis, the proteins with the higher CAI values were N, S, ORF7a, ORF3a, ORF1ab, M, and ORF8. For most ORFs the variation in CAI values was very small (between 0,001 and 0,005) but statistically significant differences were found. In the case of ORF1ab, a clear decrease in CAI was observed in all variants, being more pronounced in Delta and Gamma. The Delta variant presented the most different CAI profile, with low CAI values for most of the ORFs, except for N and ORF7b for which an increase was observed. Beta was characterized by presenting the highest CAI for S, high CAI for N, and the lowest CAI values for E and ORF3a. Alpha presented the lowest CAI for N and ORF8, and high CAI values for ORF3a and ORF1a. In this case, for most of the available genomes ORF8 was not annotated, or only a truncated version was available (On NCBI Virus resource, of 193,493 only 55 genomes presented a protein annotated as ORF8. Accessed 17-11-2021), thus the observed difference might be artificial, representing only a few shorter sequences. Finally, Gamma presented the highest CAI for ORF3a, and low CAI values for N and ORF1a.

**Figure 10.**
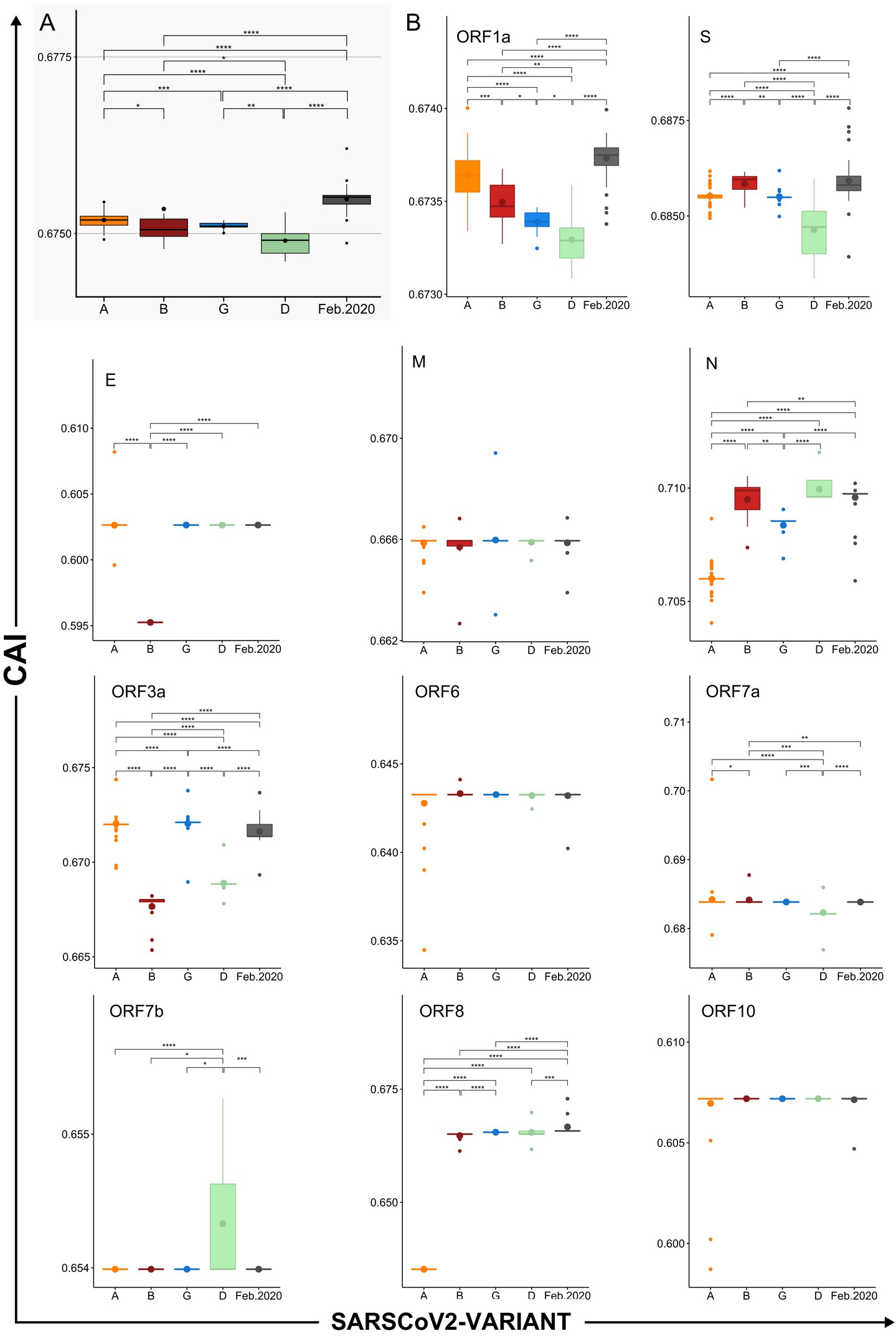
Differences in CAI between SARS-CoV-2 variants and their ORFs. CAI values calculated for concatenated SARS-CoV-2 genes using *WLei* for a selection of genomes representing different dates, variants and geographic locations. A) Box plot of CAI values grouped by variant. B) Box plots of CAI values calculated for the indicated ORFs and grouped by variant. Horizontal lines represent medians. Bigger dots represent mean values. Asterisks represent significant differences. Alpha (A), Beta (B), Gamma (G), and Delta (D) variants. P values were calculated using Wilcoxon rank sum test (* < 0.05, ** < 0.01, *** < 0.001, **** < 0.0001)

### Analysis of SARS-CoV-2 divergence in California, USA

Based on the previous results, the question of whether the variation of CAI and ENC values over time was determined only by the most abundant of the circulating variants, and not by a general trend of SARS-CoV-2 evolution, was raised. To try to answer this question, an analysis of the variation of ENC and CAI over time, considering the relative abundance of all the circulating SARS-CoV-2 variants in a determined geographic region, was made. First, we looked for the geographic region with more SARS-CoV-2 complete genomic sequences, which turned out to be California (CA) USA, with a total of 15,468 complete genomes over a total of 154,837 sequences (Date accessed, 10/05/2021). It has to be noticed that, since only complete genomic sequences were used, the results shown below do not accurately represent the real proportion of the circulating strains, which since September 2021 has been reported to be mainly Delta (https://covid.cdc.gov/covid-data-tracker/#variant-proportions; https://nextstrain.org/ncov/open/north-america. Accessed 15/10/2021). Overall, it can be seen that the ENC values for the circulating SARS-CoV-2 isolates in California show a similar profile to that of the global analysis, although with slightly higher values (Fig. S13). In the case of CAI (calculated using *WLei*) the obtained values mimic the global behavior, presenting the higher CAI on February and decreasing steadily thereon (Fig. 9, and Fig. 11.A. black curve).

**Figure 11.**
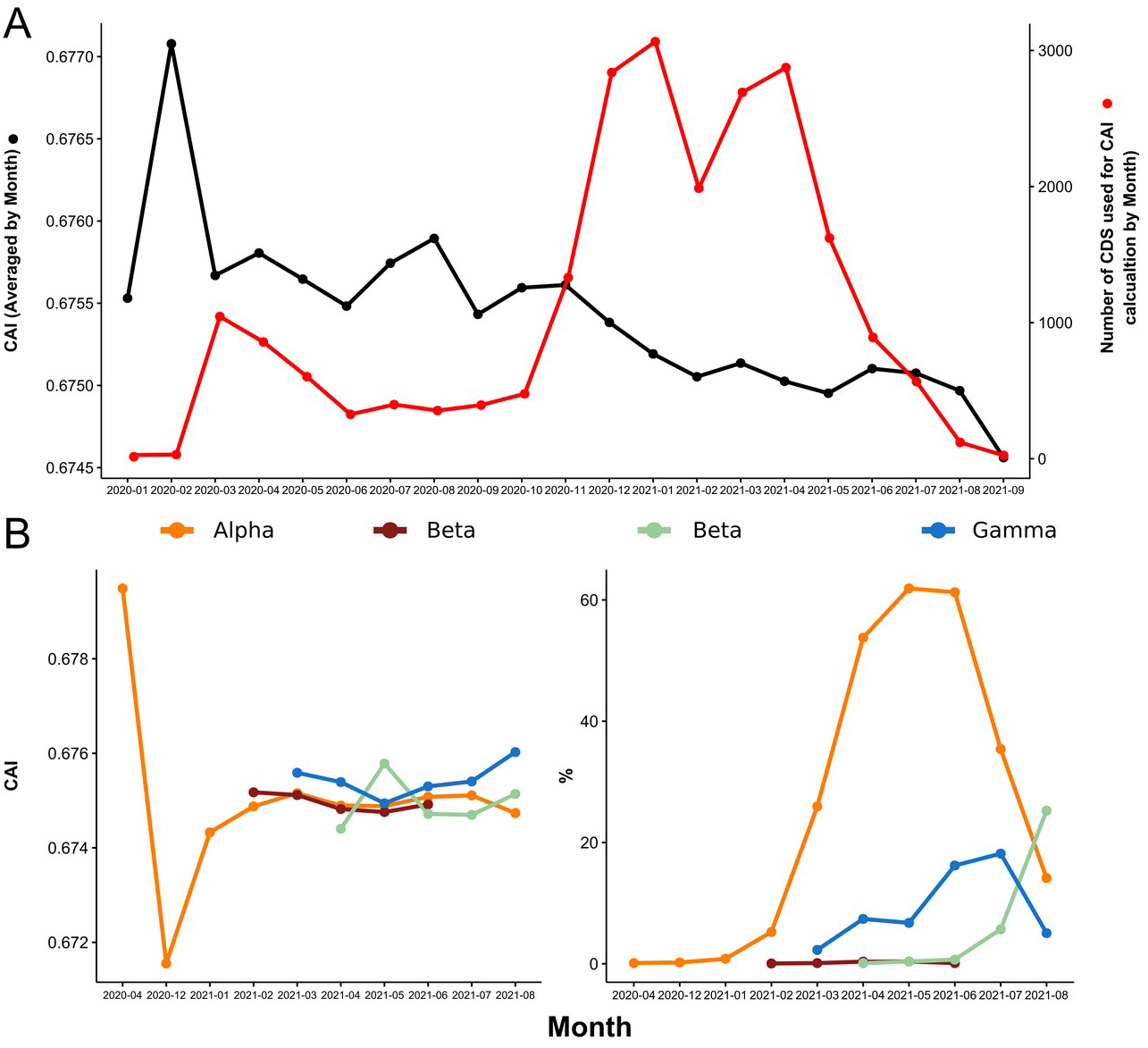
Evolution of CAI over time for the California dataset. CAI was calculated for concatenated SARS-CoV-2 genes using *WLei* and averaged by month. A) CAI calculated for the complete dataset. Black line, Evolution of average CAI values over time. Red line, total number of coding sequences (CDS) analyzed for each month. B) Left, CAI values for selected variants of interest. Right, percentage of coding sequences belonging to each variant.

The CAI spikes on February and April 2020 corresponded to the A. 1 variant (according to Pangolin lineage classification) which presents the highest CAI value (Fig. S14, Table 1S, CA-USA-Variants). Further, the decrease in CAI over time since December 2020 correlates with the most abundant variants having lower average CAI than the strains circulating on February 2020, with the Alpha (B.1.1.7), Gamma (P.1) and Delta (B.1.617.2 and AY.35) variants being the most abundant (Fig 11, Fig. S14, Table S1 – CA-USA-Variants). However, a general decrease in CAI values for each variant could not be observed, with most of them presenting nearly constant average values (Fig. 11.B, left, and Table S1 California-USA-Variants). In addition, figure S15 shows a monthly comparison between the most (Proportion > 5 %) and the less (Proportion < 1 %) abundant variants, and it can be observed that during 2021, although the most abundant variants presented lower CAI values, the difference with the less abundant variants is not significant.

Next, an analysis of CAI for all SARS-CoV-2 ORFs from the California dataset was made (Fig. 12, Fig. S16). Most of the variation in average CAI values was in the range of 0,001 units, with ORF8, N, ORF3a, ORF7a and S showing the biggest difference with time. In the case of ORF8, a particular behavior was observed because it was not annotated in the Alpha variant genomes, which were the most abundant between January and July 2021. In the case of N, most of the variation in average CAI values corresponded to the rise and fall of the A variant, which presented the lowest CAI value for N (Fig. 12 and Fig. S16). The Delta variant presented the lowest CAI values for ORF3a, ORF7a and S, and the general decrease observed for CAI since July 2021 (Fig. 11) corresponded to its onset. Remarkably, a steady increase in CAI for the S protein of the Delta variant was observed (Fig. S16), which could suggest that an adaptation of S for better expression is taking place.

**Figure 12.**
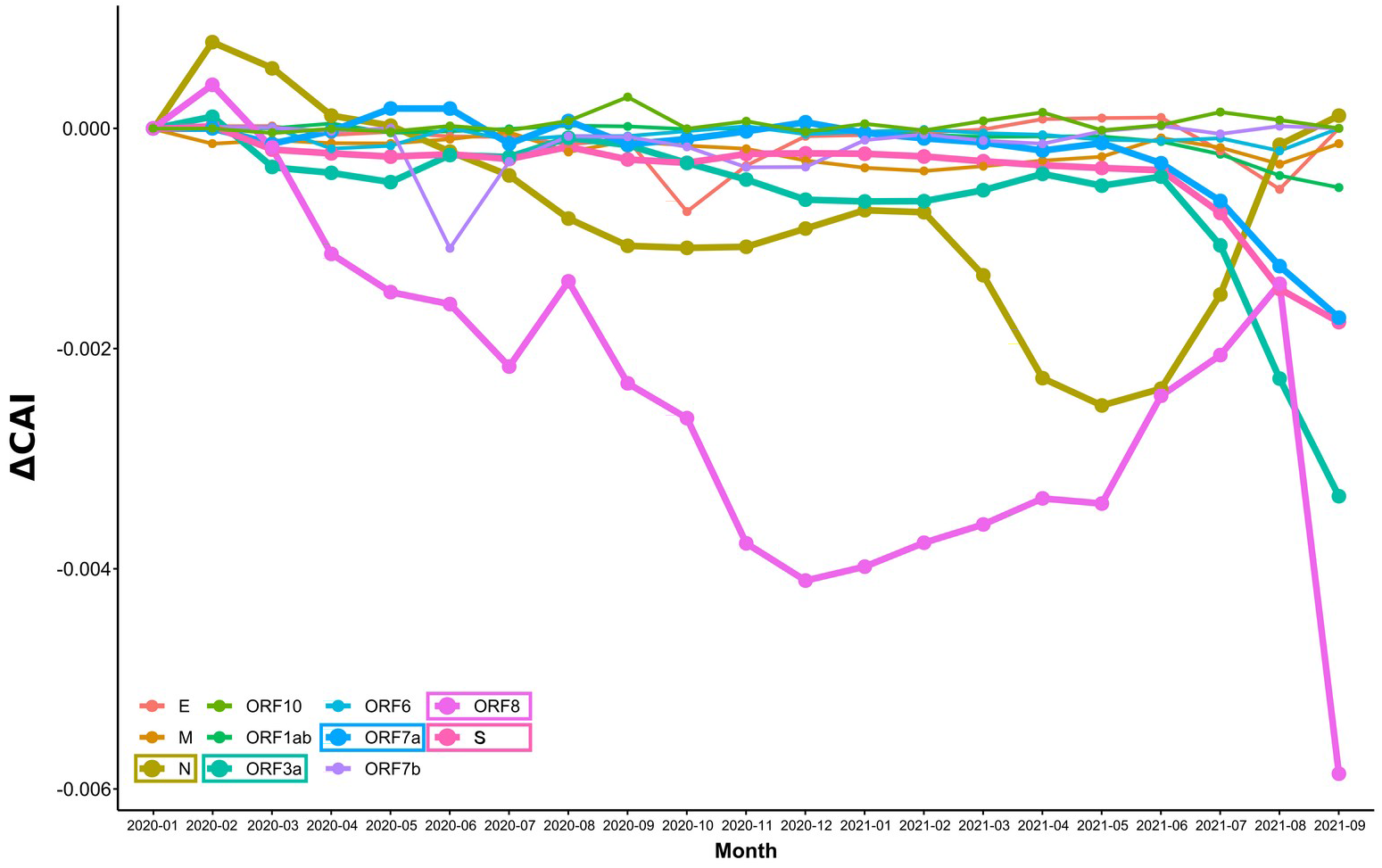
Evolution of CAI over time for each SARS-CoV-2 ORF of the California dataset. CAI values were calculated for each SARS-CoV-2 ORF using *WLei*, and averaged by month. In the figure, the difference of CAI with respect to Jan-2020 is represented (ΔCAI). Lines for the ORFs with greater variation over time are thicker (N, S, ORF3a, ORF7a, and ORF8).

### Omicron variant

On 26 November 2021, WHO designated the variant B.1.1.529 a variant of concern, named Omicron [73]. This variant presents a large number of mutations, some of which are concerning [74], and the apparent capacity to infect people who recovered from COVID-19 caused by Delta and other variants [75]. Whether Omicron causes milder or more severe disease is still unknown, although most of the reports, which had occurred in the younger population, presented mild symptoms. Here, we downloaded the complete genome of the Omicron variant from NCBI Virus (Accession number: OL672836) and calculated the ENC and CAI (calculated using *WLei*) values as a measure of codon adaptation. Our results (Table S2, Omicron ENC and CAI) show that Omicron presents an ENC value greater than the average for other SARS-CoV-2 isolates (ENC, 45.51), although not the highest registered in our study. It also has a higher CAI, compared to the other variants of concern (CAI, 0.675), but lower than the CAI values for SARS-CoV-2 isolates from the first half of February 2020. The ENCs for the individual ORFs were also calculated, showing that Omicron presents a different profile, with higher ENCs for ORF1ab and S, and lower ENCs for E and ORF3a. Particularly low ENC values were found for M and ORF7b proteins. In the case of CAI, S and M presented some of the higher values, while E, N and ORF7b some of the lower, when compared to other SARS-CoV-2 isolates.

## Discussion

The COVID-19 pandemic provided an unprecedented dataset of complete genomic sequences, with complete metadata including the sample collection date, which made possible the study of codon usage bias and its adaptation to the human host in short periods of time. First, using both maximum likelihood phylogeny and Correspondence Analysis (CA) of Average Codon Usage Frequencies (ACUF), we corroborated that SARS-CoV-2 is highly related to the SARS-related coronavirus RatG13 isolated from Bats, and to Pangolin CoV [45, 47], being more distantly related to SARS-CoV and MERS-CoV. In addition, the CA showed that SARS-CoV-2 was farther from the human tissues than SARS-CoV and MERS-CoV, suggesting that it is less adapted than the latter for protein expression in the human host. Previous reports indicated that coronaviruses show low substitution rates over time, normally in the range of 1 × 10^-4^ to 1 × 10^-3^ substitutions per site per year [69], with a total of around 5 to 14 nucleotide differences between independent SARS-CoV-2 isolates in the first half of 2020 [69, 70]. Here approximately 1 × 10^-3^ substitutions per site (i.e. approximately 30 SNPs per genome) were observed for the most distant SARS-CoV-2 2021 isolates. This indicated that it should be possible to analyze the time dependence of codon usage frequencies for SARS-CoV-2 ORFs, although only a small variation was expected, as it was previously observed by Hussain *et al. [40]* and Huang *et al*. [43]. These authors conducted similar time-series analysis, principally of averaged ENC and CAI values, showing that in the first months of COVID-19 pandemic both indices were slowly decreasing. In the analysis presented here, a dataset of sequences representing different geographic locations and SARS-CoV-2 variants, and moving along to July 2021, was analyzed. A trend in codon usage bias variation over time was observed in the CA of ACUF, which shows that the distance between SARS-CoV-2 and the highly expressed genes in different human tissues is slightly increasing on the principal axis (C1) and decreasing on the secondary axis (C2). C1 is mostly defined by U/A and C/G terminated codons and accounts for 73% of the total inertia. These results are in accordance with previous reports that showed that +ssRNA viruses possess A rich and C poor genomes, with a depletion of CpG and UpA dinucleotides, and third codon positions enriched in U [40, 76]. Such an increase in U ending codons could be caused both by selection or mutational biases, which are considered as the main forces of RNA virus evolution, due to their large population sizes and high mutation rates. Also, it has been shown that mutation is universally biased toward A and T in several species [77, 78] In the particular case of +ssRNA viruses, including SARS-CoV-2, a mutational bias towards U, which in CoVs was produced mainly by C→U mutations, was observed [77, 78]. This bias is remarkable in SARS-CoV-2, which was shown to present a high proportion of C→U changes relative to other types of SNP [69], with approximately a 4-fold excess of C→U substitutions [70]. However, mutations have to be fixed in the population, a process that takes time, and may be incomplete in SARS-CoV-2 circulating variants. Using the notion of incomplete purifying selection, it was proposed that in the longer term, selection towards A and against U takes place [76]. Whether the mutations appearing in SARS-CoV-2 circulating variants have been fixed is difficult to determine. Neutrality plots results suggested a minor effect of mutation bias and major effect of natural selection [40]. Here, using the California dataset, a linear dependence of GC12 (i.e., GC percentages of codon positions 1 and 2) with GC3 (i.e., GC percentages of codon position 3) (R^2^: 0.2455, p-value: < 2.2e-16) with a slope of 0.12370 was observed, suggesting that only 12% of the codon usage bias could be attributed to mutational bias. Moreover, a neutrality plot was made for sequences clustered by month (Table S2, Neutrality plots vs Date), indicating in average less than 20% of mutation bias contribution to the codon usage bias. An overall dependence with time was not found, however, spikes of lower slope (major contribution of selection) were observed on Sep-2020; Jan-2021 and May-2021. In those months the prevalent circulating variants were B.1 and B.1.243; B.1.2, B.1.427 and B.1.429; and B.1.1.7 (Alpha), respectively (Table S1, California-USA-Variants), which could indicate that in those variants a more complete selection may have taken place.

The obtained results are also compatible with previous codon usage analyses in SARS-CoV-2, where an antagonistic codon usage pattern with the human host was observed [37], and it has been suggested that it could play a role during the initial phase of the infection, reducing translation speed, but increasing its precision, and yielding accurate and correctly folded viral proteins [79]. Also, a slow viral translation and replication, may help the virus to avoid detection by the host immune system [40].

SARS-CoV-2 has a high average ENC value of around 45, indicating a low to moderate codon usage bias, which is lower than the average ENC values of human tissues (approximately 55, Fig. S7). It has been suggested that a weak codon usage bias might be an adaptive trait enabling viruses to replicate, without competing for the limited t-RNA resources, in a broader range of hosts presenting different codon usage patterns [37, 39, 40]. Recent reports have shown that SARS-CoV-2 ENC values have decreased over time in the first half of 2020, here this tendency was reinforced and extended to July 2021. A smaller ENC indicates that SARS-CoV-2 codon preference has increased, however, it doesn’t mean that it is adapting to the human codon usage pattern. In fact, the CA results and the analysis of the variation of ACUF over time indicate a greater polarization.

To test whether this small but significant variation in the codon usage bias could enhance the expression of SARS-CoV-2 proteins in humans, CAI was calculated using as reference the highly expressed proteins in different human tissues. CAI is accepted as an effective index of the degree of viral adaptation to a host’s cellular environment [80]. That SARS-CoV-2 presents lower CAI values in comparison to MERS-CoV and SARS-CoV, has been interpreted as a lower fitness and adaptation to human cellular systems, which is also in agreement with its milder clinical picture [37]. Moreover, our results indicate that SARS-CoV-2 proteins should present a higher expression in granulocytes, kidneys, heart, and the pituitary and thyroid glands; and lower expression in the skin, stomach, intestine, dendritic cells, monocytes and eyes. These results are partially in agreement with previous reports showing SARS-CoV-2 tropism for lungs, trachea, kidneys, heart, pancreas, brain and small intestine, but not for the large intestine, renal proximal tubules, and liver [81, 82]; and also with the prediction of vulnerable cells types based on the ACE2, TMPRSS2 and Furin expression profiles (i.e. lung AT2 cells, macrophages, cardiomyocytes, adrenal gland stromal cells, stromal cells in testis, ovary and thyroid cells) [83, 84]. Further, different types of immune cells can be infected, including granulocytes [85] which have been reported to be key modulators in SARS-CoV-2 immune response [84]; and also new evidence suggests that SARS-CoV-2 might have a great impact on the hypothalamus-pituitary-thyroid endocrine axis [87]. It should be noted that the alteration of the target cell (or tissue) normal functions will depend, not only on its susceptibility to SARS-CoV-2 infection, but also on the expression level of all SARS-CoV-2 proteins.

It has been observed, both here and in a previous report by Huang *et al*. [43], that CAI values for SARS-CoV-2 in humans have decreased over time, and it was hypothesized that it was likely that the efficiency of gene expression of SARS-CoV-2 in the human host could be decreasing. That SARS-CoV-2 CAI values are decreasing raises the question of whether MERS-CoV and SARS-CoV higher CAI values are a consequence of a larger adaptation time in the human host, or if their CAI values were higher since the beginning, and they are actually evolving towards a lower CAI.

An analysis of the variants Alpha, Beta, Gamma and Delta, revealed a difference of ACUF with the early 2020 isolates, and a differential codon usage bias between Gamma, Delta, and Alpha and Beta isolates which were somewhat overlapped. Particularly Delta and Gamma isolates, presented a higher ENC, which may explain the increase in the average ENC observed in the last months of analysis. With respect to CAI, all the analyzed variants presented lower CAI values when compared to February 2020 isolates, with Delta and Gamma presenting the lower values.

Finally, an analysis of each SARS-CoV-2 gene was made. It was previously reported that ORF1ab, S and N are the proteins accumulating most of the mutations [49], however, of these proteins only N presented considerable variation on its codon usage bias according to the CA results. In the case of M and E, which have been reported to evolve more slowly [39], only E presented some variation in its codon usage bias. Other proteins that showed some variation in the CA were ORF6, ORF7b, and ORF10. However this variation was not always reflected on the ENC and CAI values.

As a common trend, the variation of ENC and CAI over time was minimal, which was expected due to the small divergence time, and the fact that a ratio of non-synonymous to synonymous substitutions of 1.88 had been previously reported [69]. Nevertheless, significant differences were found, especially when Alpha or Delta variants became predominant. In particular, N, M, ORF7a and ORF3a presented high CAI and ENC values, with values similar to those registered for human proteins, and which indicated that these proteins may be required in higher amounts. A very good accordance of their CAI values with the ribosome profiling experiments reported by Finkel *et al*. [13] was found, being the most actively translated proteins N, M, ORF7a, ORF3a, ORF8, ORF6, ORF7b, S and E in descendant order. The nucleocapsid phosphoprotein (N), was among the proteins which presented more variation in the codon usage bias between SARS-CoV-2 isolates, and also the higher average CAI value. These facts could suggest a particular role of N in the evolution and adaptation of beta-CoVs to their mammal hosts.

In contrast, S, ORF8, ORF7b, and ORF6 presented relatively high CAI but lower ENC values. Proteins with a lower ENC use a more restricted set of codons, and if those codons are common with those used by highly expressed host proteins (i.e. viral proteins with higher CAI), a more effective competition for the aminoacilated t-RNA will take place. In accordance, Alonso and Diambra [57] observed a reduced translation rate of highly expressed host proteins which shared the codon usage bias of the virus, and the same approach was used by Maldonado *et al*. [58] to identify human genes that could be potentially deregulated due to the codon usage similarities between the host and the viral genes. Thus, a reduction of CAI over time, as was observed for S and ORF7a, may be compatible with a milder pathogenicity. ORF7b has the lowest ENC value of about 30, a relatively high CAI of 0.654, and protein levels similar to S [13]. The particular profile presented by ORF7b in the Delta variant, might explain some of the clinical differences presented by this variant. It was demonstrated that N mutations, as the N:R203M contained in the Delta variant, can produce an enhanced RNA packaging and replication, an improved fitness, and could also explain the increased spread of variants [88]. That N presented the most optimized codon usage profile in Delta could contribute to its increased fitness. E presented the lowest CAI, and its value remained nearly constant over time, with the observation that the Beta variant presented differentially lower CAI and higher ENC values. Also, the Beta variant presented high CAI for N and S. A recent report has informed that people infected with the Beta SARS-CoV-2 variant were more likely to need critical care and to die than are people infected with other variants, and it also seems to be more resistant to immunity generated by vaccines or previous infections [89]. The particular CAI/ENC profile uniquely shown by the Beta variant, probably involved in different expression levels of E protein, could be related with this variant severity. The Omicron variant appears to be more transmissible, and less pathogenic. These features could be related to the higher CAI for S, and the lower CAI for ORF7b respectively. ORF7b also presents the lowest ENC in the Omicron variant, suggesting a more antagonistic, yet restricted, codon usage, which might be related to this strain ability to avoid the immune response [90].

Finally, it has been proposed that the low adaptation of SARS-CoV-2 to the human codon usage could be a consequence of its recent transit from a well-adapted host, or an evolutionary strategy to avoid host defense [40]. However, most of the results seem to indicate that SARS-CoV-2 codon usage is getting further apart, instead of adapting for a higher and faster protein expression. Deoptimization of codons and codon pairs has been used as an attenuation strategy for viral vaccine development [37]. That SARS-CoV-2 codon usage pattern is getting away from that of the human host, and the decreasing CAI and ENC values observed since the onset of the pandemic, could indicate that the virus is evolving to be less pathogenic [40], and might end, with time, being similar to other CoVs causing common cold. It must be considered however, that this conclusion is only based on the evolutionary trends observed in the codon usage profile, and although they could exert a significant effect in SARS-CoV-2 pathogenicity, the occurrence of novel non-synonymous substitutions as the observed in the recent Omicron variant [74], will present a more direct effect.

## Funding

This research received no specific grant from any funding agency in the public, commercial, or not-for-profit sectors. M. J. L. and D.B. are researchers supported by the National Science and Technology Research Council (Consejo Nacional de Investigaciones Científicas y Técnicas - CONICET, Argentina), and the National University of La Plata (Universidad Nacional de La Plata, Argentina); E. G. M. is a PhD student supported by a grant from the CONICET.

## Conflicts of interest

The authors declare no conflict of interest.

## Declaration of competing interest

The authors declare that they have no known competing financial interests or personal relationships that could have appeared to influence the work reported in this paper.

## Supplementary material

Supplementary figures and tables are accessible through Github (https://github.com/maurijlozano/SARS-CoV-2-CodonUsage)

## Notes

https://github.com/maurijlozano/SARS-CoV-2-CodonUsage

### Competing Interest Statement

The authors have declared no competing interest.

