## Supplementary material. for "Analysis of SARS-CoV-2 synonymous codon usage evolution throughout the COVID-19 pandemic"

#### Figures

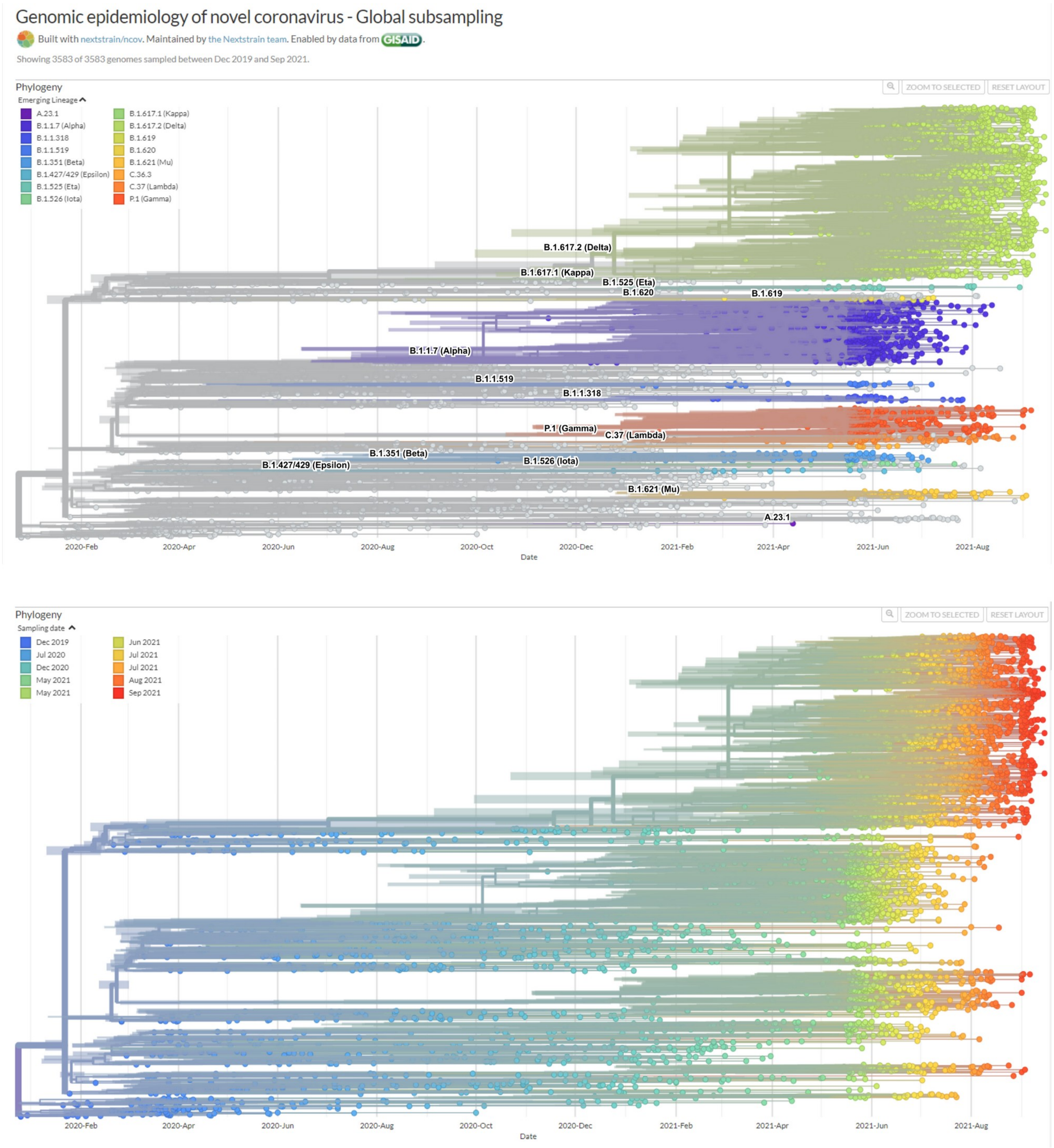


**Figure S1. Genomic epidemiology of novel CoVs – Global subsampling extracted from Nextstrain.**

Phylogenetic trees colored by variant and date. Extracted from Nextstrain web page.

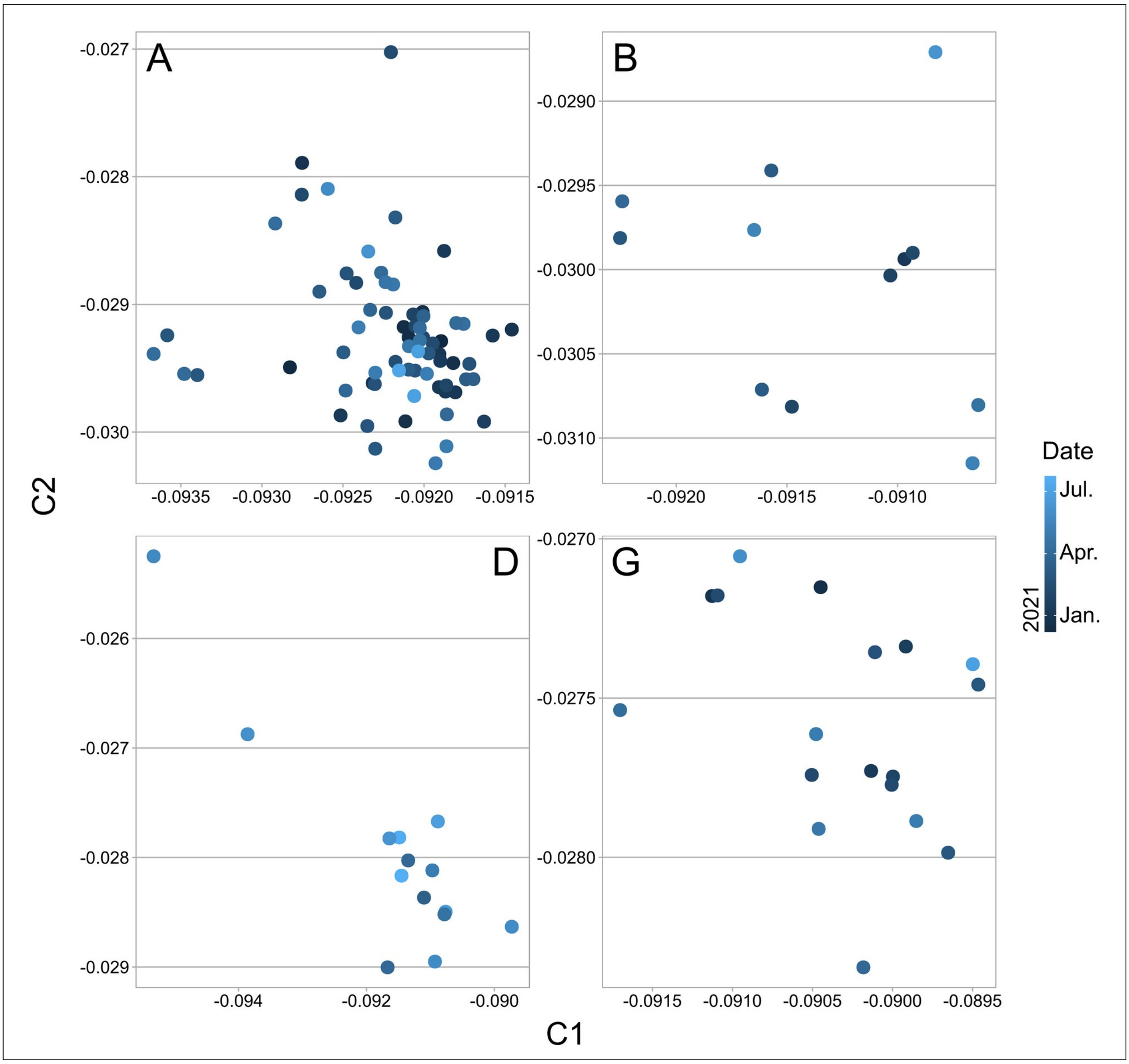


**Figure S2. Correspondence Analysis of concatenated SARS-CoV-2 genes averaged by date and grouped by variant.** A, Alpha; B, Beta; G, Gamma; D, Delta. Dark to light blue colors indicate dates from January 2021 to July 2021**.**

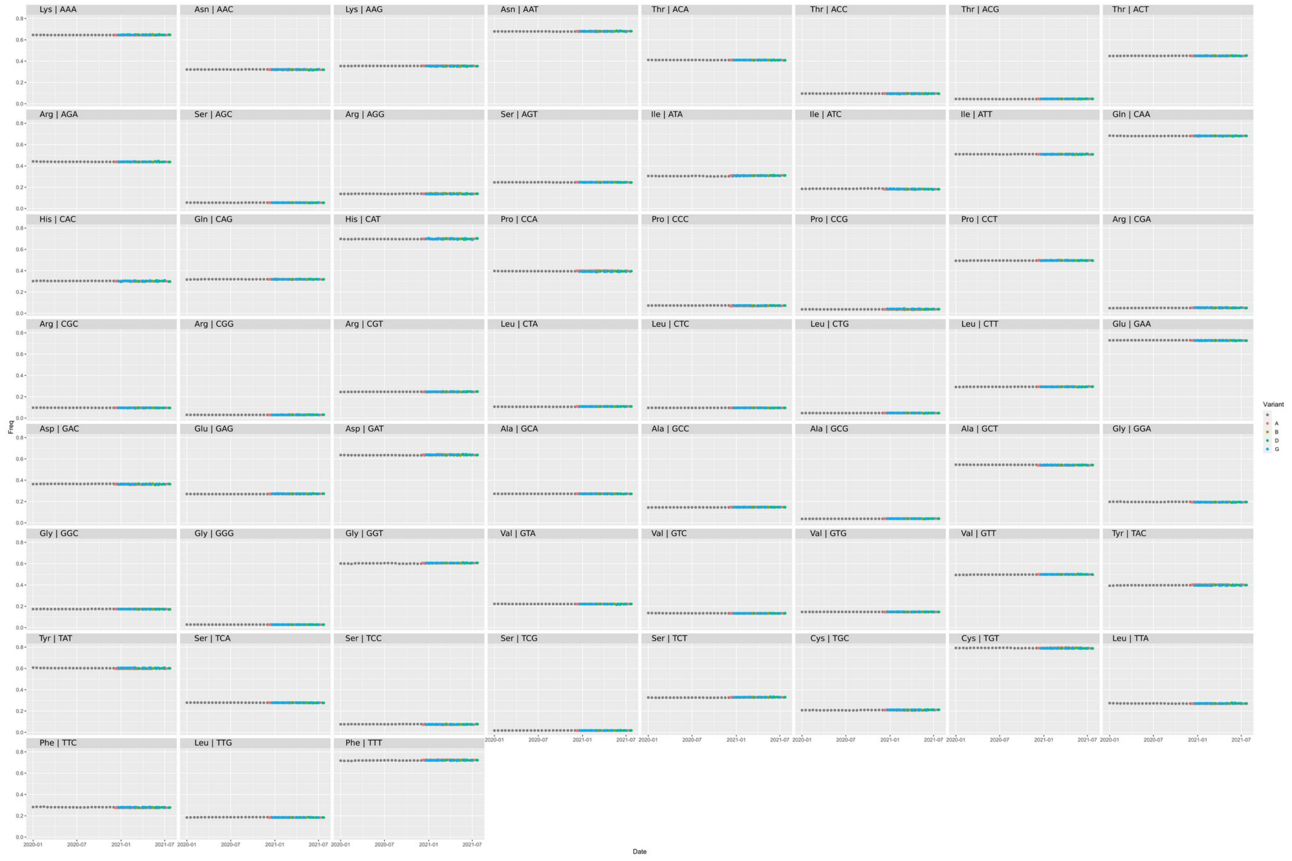


**Figure S3. Evolution of the Average Codon Usage Frequency for each codon over time.** ACUFs were calculated for concatenated SARS-CoV-2 genes grouped by fortnight, and plotted using the same scale ranging from 0 to 1. Black points represent the ACUF for SARS-CoV-2 isolates from the time series dataset. Color points correspond to selected SARS-CoV-2 variants of interest: A (Alpha) orange, B (Beta) dark red, D (Delta) light green and G (Gamma) blue.

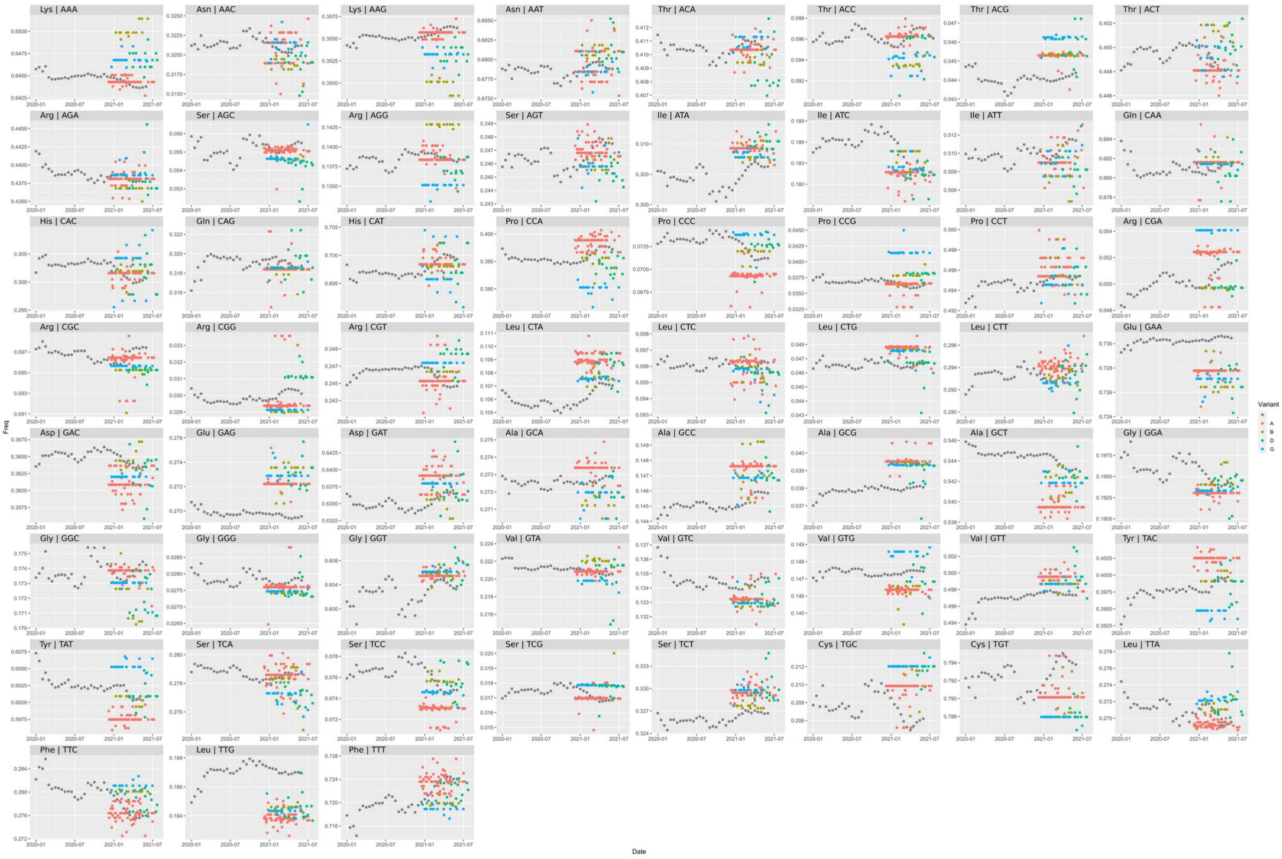

**Figure S4. Evolution of the Average Codon Usage Frequency for each codon over time with independent scales.** ACUFs were calculated for concatenated SARS-CoV-2 genes grouped by fortnight, and plotted using independent scales for each codon. Black points represent the ACUF for SARS-CoV-2 isolates from the time series dataset. Color points correspond to selected SARS-CoV-2 variants of interest: A (Alpha) orange, B (Beta) dark red, D (Delta) light green and G (Gamma) blue.

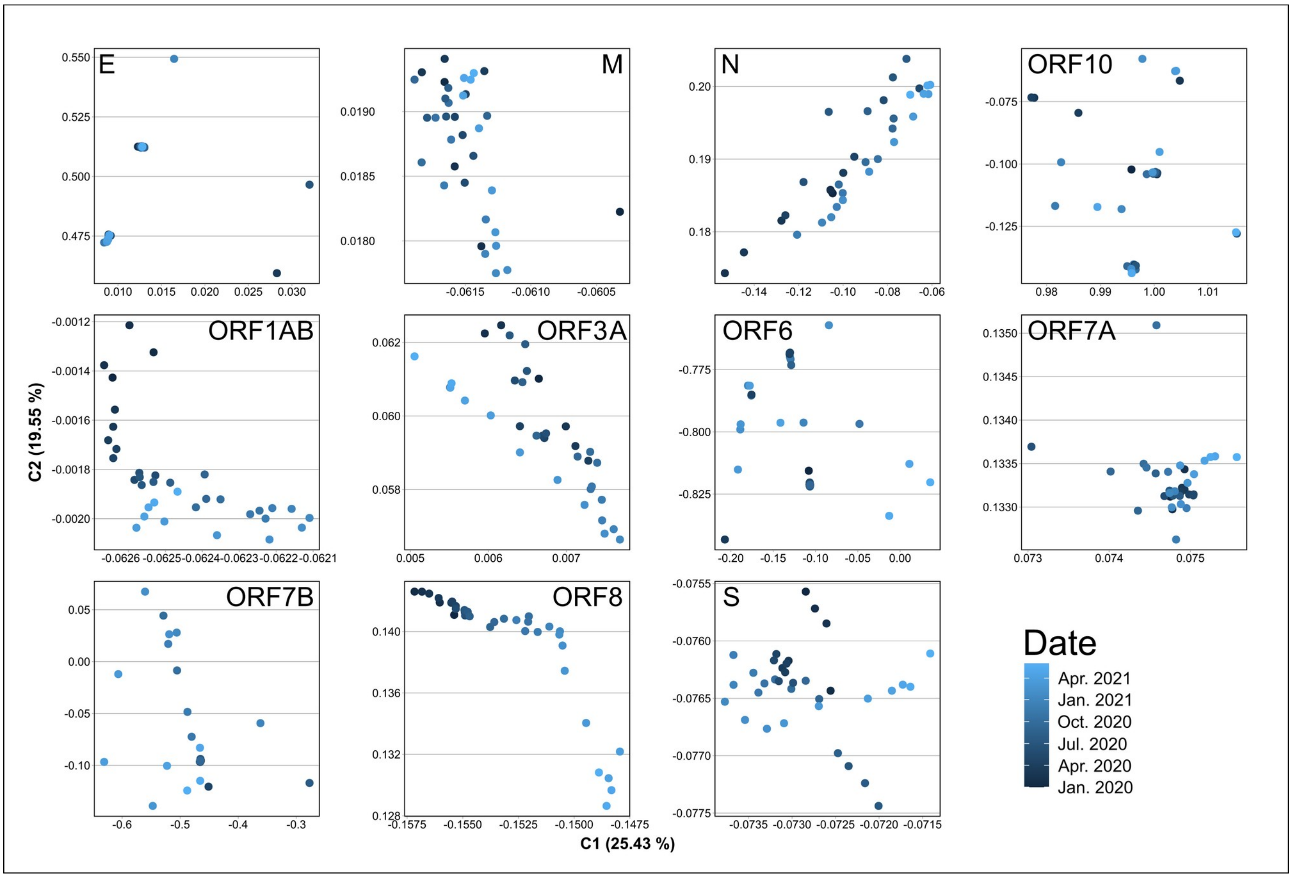


**Figure S5. Correspondence Analysis of ACUF for each SARS-CoV-2 ORF from the time-series dataset averaged by month.** The coding sequences corresponding to ORF1ab, S, M, N, E, ORF3a, ORF6, ORF7a , ORF7b, ORF8, and ORF10 were extracted from the SARS-CoV-2 time-series dataset, their ACUF were calculated, averaged by month, and CA was performed. A separated graphic for each ORF is shown. Dark to light blue colors indicate dates from Jan-2020 to Jun-2021.

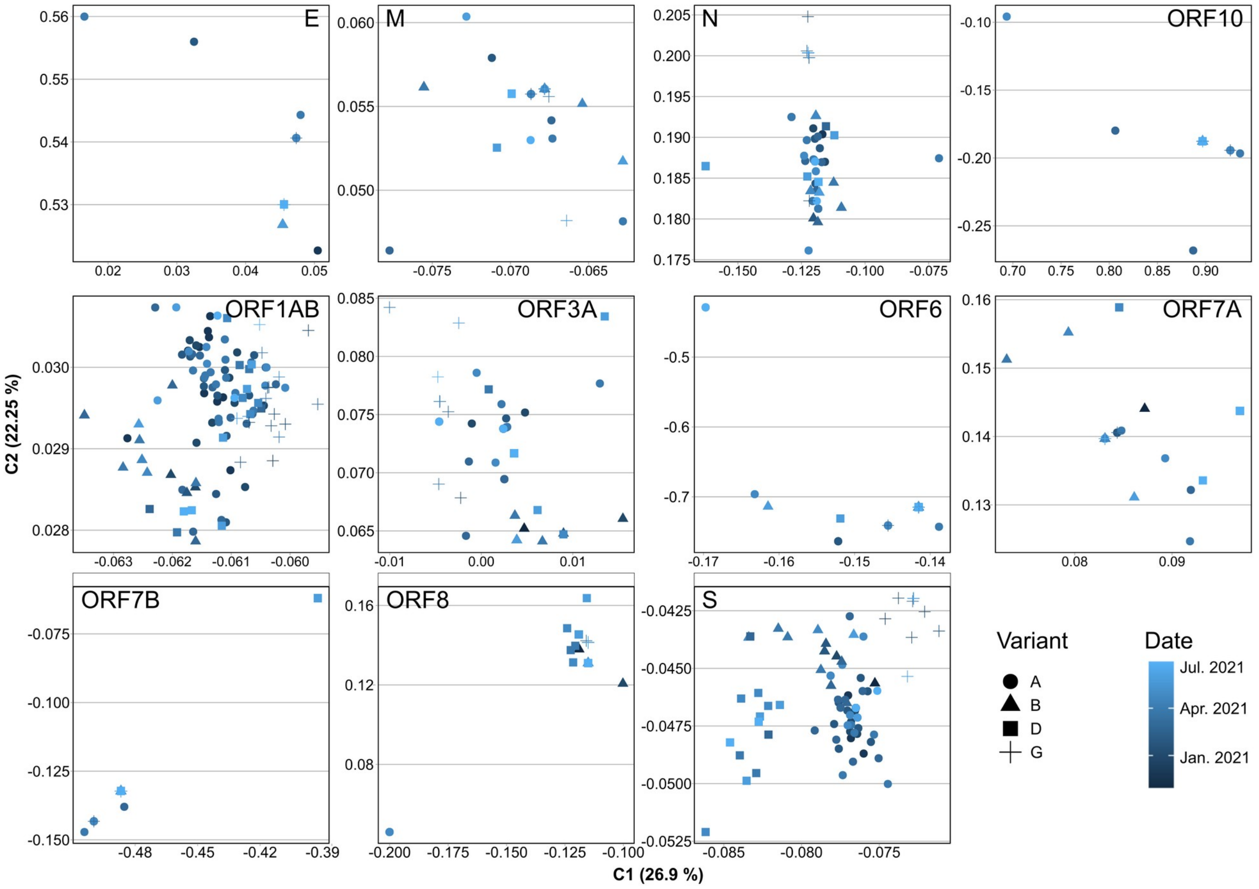


**Figure S6. Correspondence Analysis of Codon Usage Frequencies for each SARS-CoV-2 ORF from Alpha, Beta, Gamma and Delta variants.** The coding sequences corresponding to ORF1ab, S, M, N, E, ORF3a, ORF6, ORF7a , ORF7b, ORF8, and ORF10 were extracted from the genomes of SARS-CoV-2 Alpha (A), Beta (B), Gamma (G), and Delta (D) variants, their codon usage frequencies were calculated and CA was performed. A separated graphic for each ORF is shown. Shapes indicate the different SARS-CoV-2 variants. Colors indicate the date, from December 2020 (Dark blue) to July 2021 (light blue).

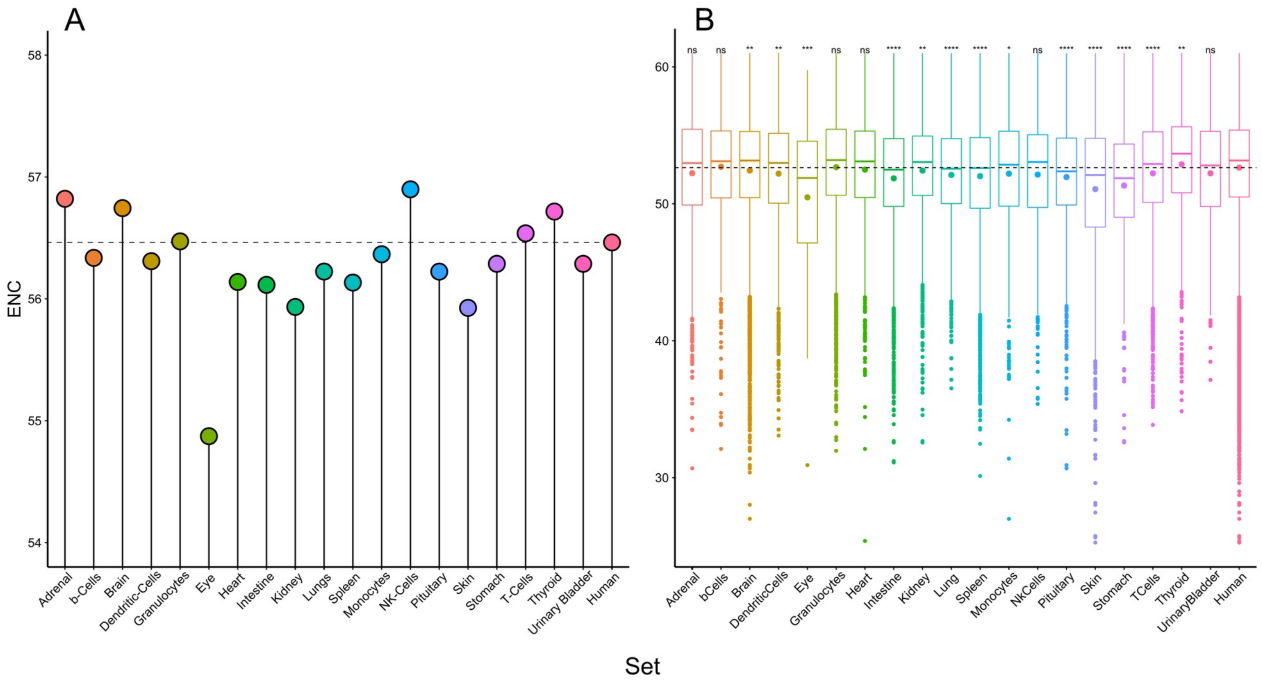


**Figure S7. Effective Number of Codons (ENC) for elevated proteins on each human tissue.** ENC values were calculated either, for the concatenated (A) or individual genes (B) corresponding to elevated proteins on each human tissue. In figure B (Boxplots) the middle lines represent medians, while the dots inside the boxes represent average ENC values.

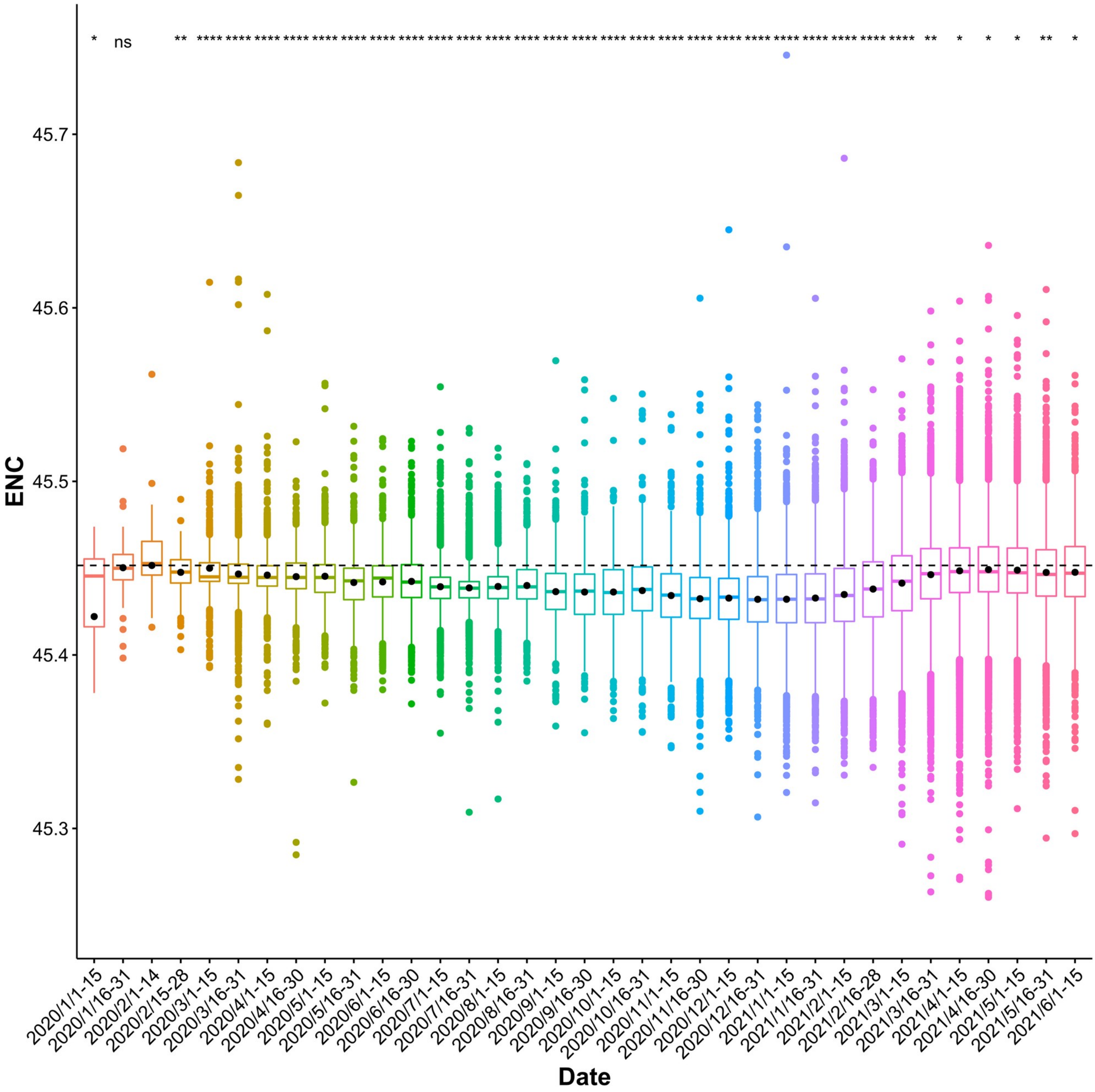


**Figure S8. Evolution of ENC values calculated for concatenated SARS-CoV-2 genes over time.** In the Boxplots, the middle lines represent the median, while the dots inside the boxes represent average ENC values. The dotted line at 45.45 indicates the average ENC value for February 2020. Asterisks represent significant differences with February 2020. P values were calculated using Wilcoxon rank sum test (* < 0.05, ** < 0.01, *** < 0.001, **** < 0.0001).

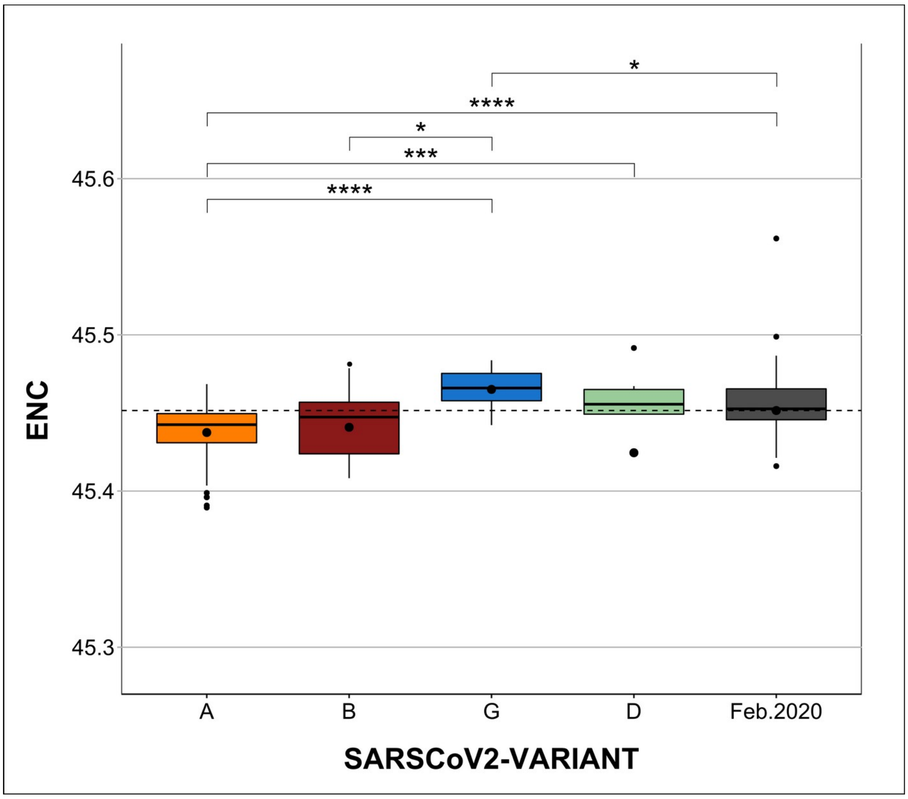


**Figure S9. Differences in ENC between SARS-CoV-2 variants.** ENC values calculated for concatenated SARS-CoV-2 genes of genomes selected by variant. A (Alpha), B (Beta), G (Gamma), and D (Delta). The dotted line at 45.45 indicates the average ENC value for February 2020. In the Boxplots, the middle lines represent the median, while the dots inside the boxes represent the average ENC values. Some outliers were left out of the figure for clarity. Asterisks represent significant differences. P values were calculated using Wilcoxon rank sum test (* < 0.05, ** < 0.01, *** < 0.001, **** < 0.0001) .

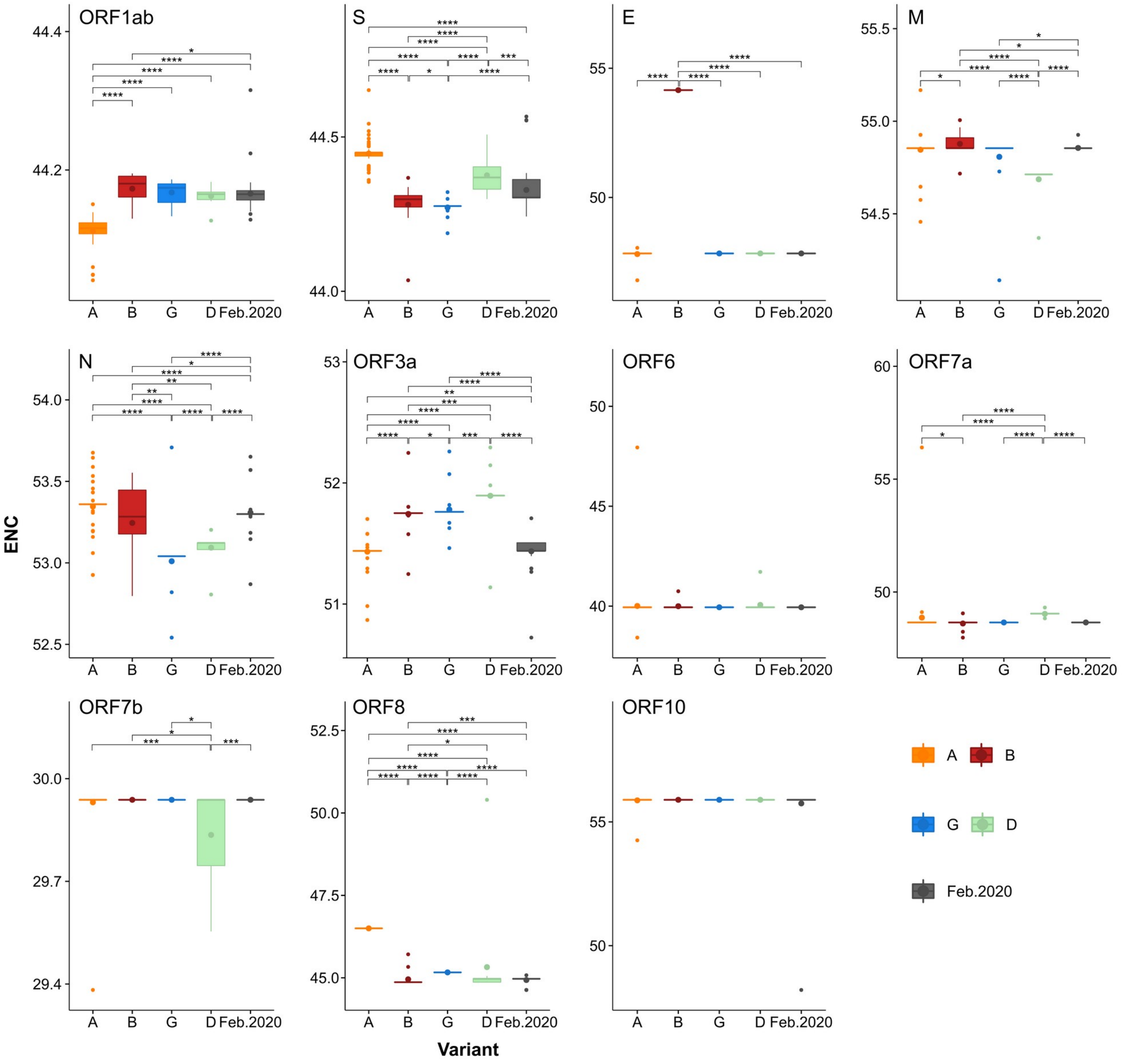


**Figure S10. Differences in ENC between SARS-CoV-2 variants for each ORF.** ENC values were calculated for individual SARS-CoV-2 ORFs for a selection of genomes representing different variants. A (Alpha) orange, B (Beta) dark red, D (Delta) light green and G (Gamma) blue, black represents Feb-20 isolates. In the Boxplots, the middle line represents the median, while the dots inside the boxes represent average ENC values. Asterisks represent significant differences. P values were calculated using Wilcoxon rank sum test (* < 0.05, ** < 0.01, *** < 0.001, **** < 0.0001).

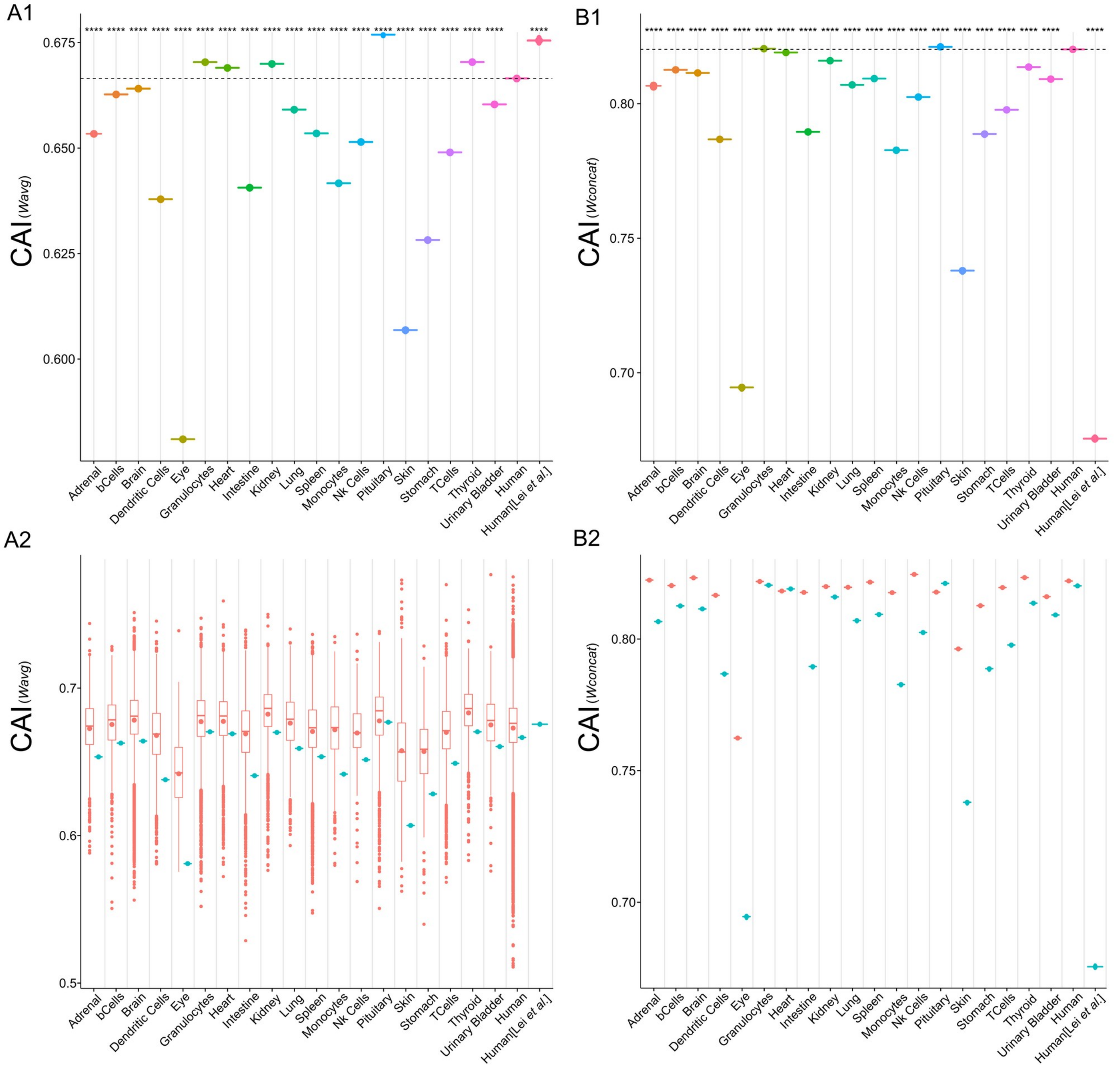


**Figure S11. CAI values for concatenated SARS-CoV-2 genes calculated using highly expressed proteins from different human tissues as reference.** A1 and B1) CAI values were calculated for concatenated genes of SARS-CoV-2 February 2020 isolates, using the highly expressed proteins in different human tissues as reference sets. For A1 and A2, CAI values were calculated using *Wavg*, while for B1 and B2, CAI values were calculated using *Wconcat*. A2 and B2 Show the tissue specific differences in CAI between SARS-CoV-2 (same values as A1 and B1, in Cyan) and human genes (dark pink). In A2, CAI values were calculated for all the highly expressed genes in each tissue using the corresponding *Wavg*. In B2, CAI was calculated for the concatenated highly expressed genes in each tissue using the corresponding *Wconcat*. Human[Lei *et al*.] label corresponds to CAI values calculated for the concatenated SARS-CoV-2 genes using *WLei* (same values for A1,A2,B1 and B2). In the Boxplots, the middle lines represent the median, while the dots inside the boxes represent average CAI values. In A1 and B1, the dotted line indicates the SARS-CoV-2 CAI values calculated using all the coding sequences from the human genome as reference set. Asterisks represent significant differences. P values were calculated using Wilcoxon rank sum test (* < 0.05, ** < 0.01, *** < 0.001, **** < 0.0001).

**[PDF File]**

**Figure S12. Evolution of CAI values calculated for concatenated SARS-CoV-2 genes over time.** CAI values were calculated for concatenated SARS-CoV-2 genes using reference sets corresponding to highly expressed proteins in different human tissues. A figures) CAI values calculated using *Wavg*. B figures) CAI values calculated using *Wconcat*. In the Boxplots, the middle lines represent the median, while the dots inside the boxes represent average CAI values. Dotted line: average CAI value for February 2020. Asterisks represent significant differences with February 2020. P values were calculated using Wilcoxon rank sum test (* < 0.05, ** < 0.01, *** < 0.001, **** < 0.0001).

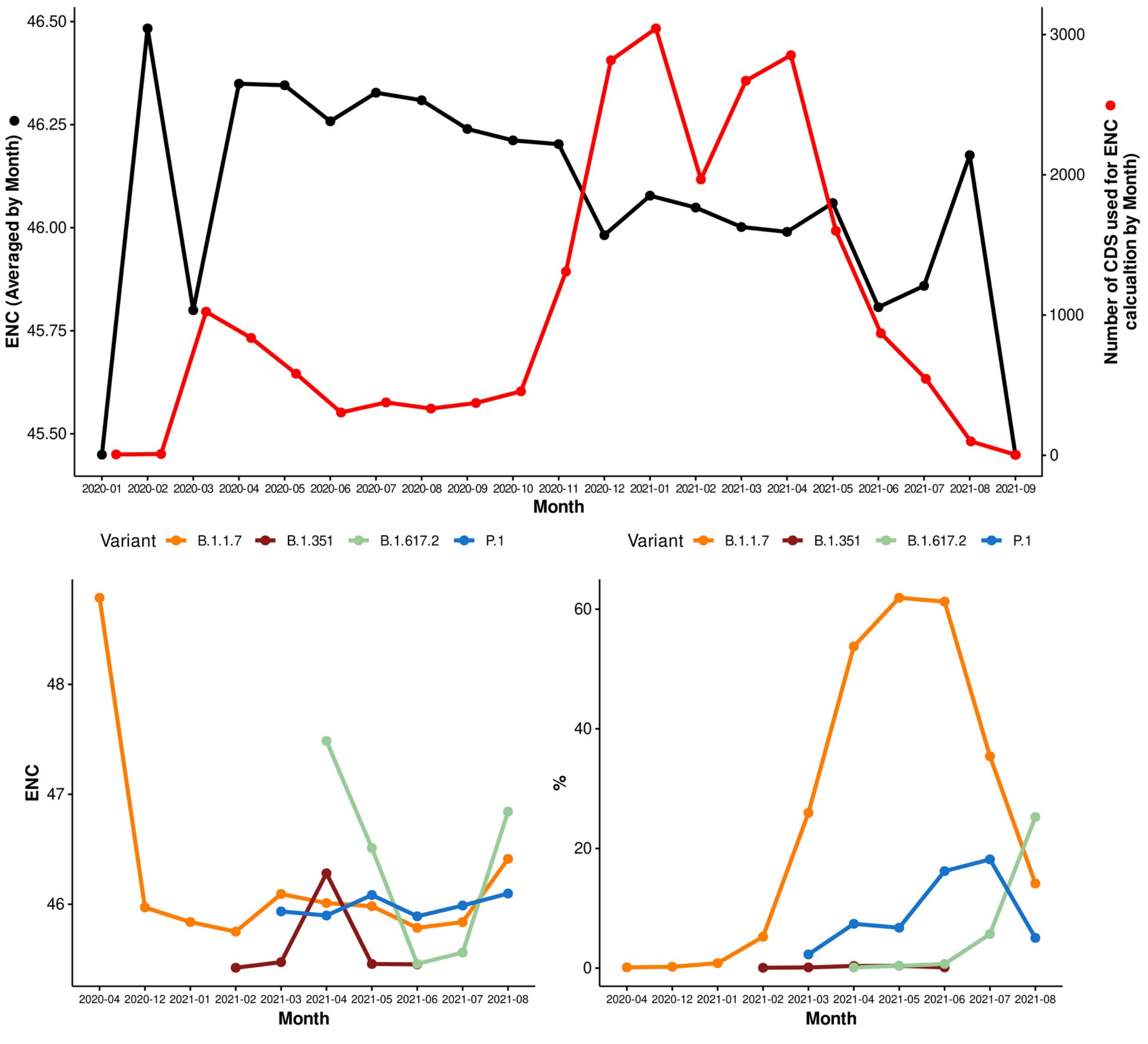


**Figure S13. Evolution of ENC over time for the California dataset.** ENC was calculated for concatenated SARS-CoV-2 genes and averaged by month. A) ENC calculated for the complete dataset. Black line, Evolution of average ENC values over time. Red line, total number of coding sequences (CDS) analyzed for each month. B) Left, ENC values for selected variants of interest. Right, percentage of coding sequences belonging to each variant.

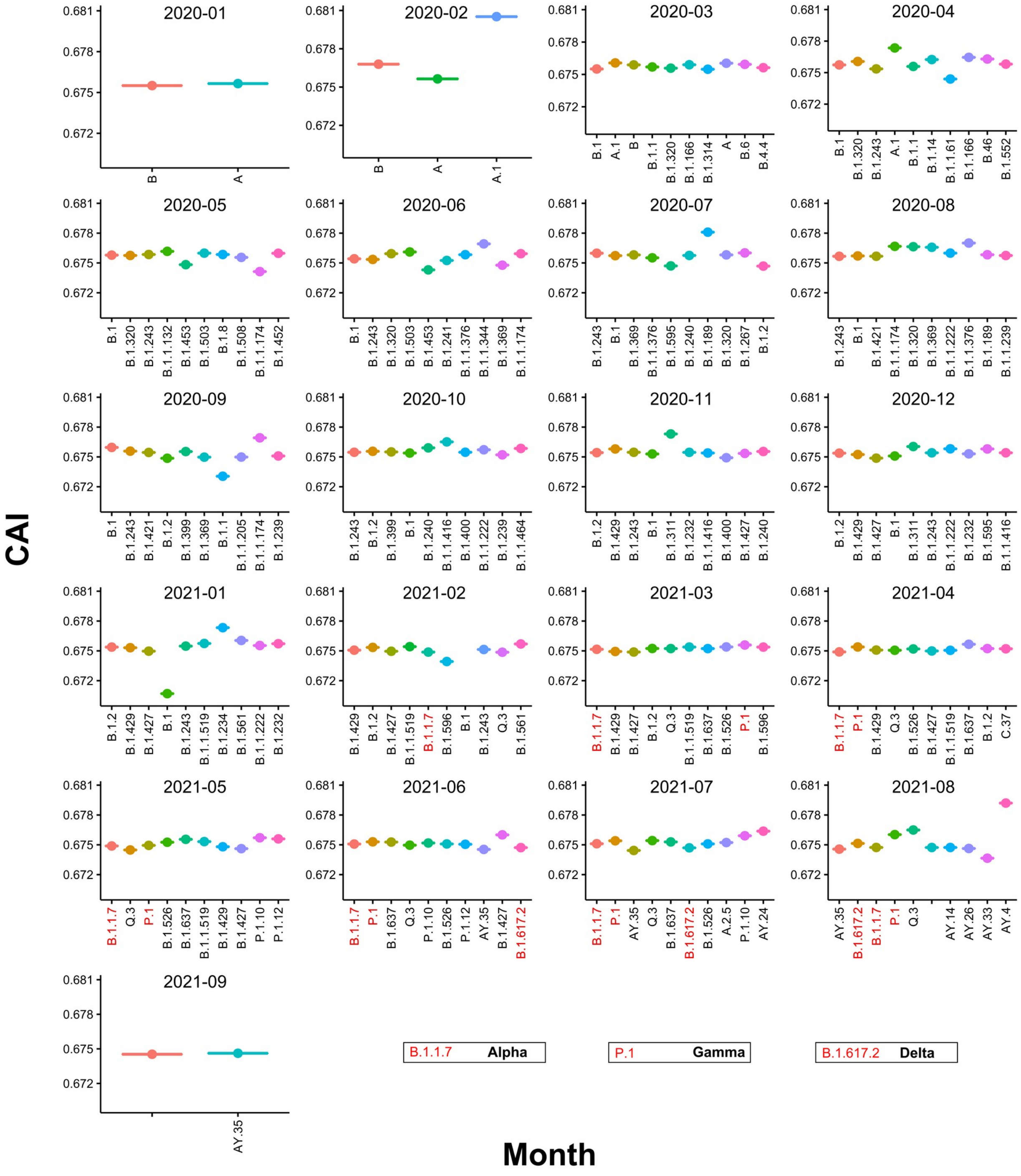


**Figure S14. Average CAI values for the ten most represented variants in the California dataset from Jan-2020 to Sep-2021.** CAI values were calculated for concatenated SARS-CoV-2 genes using *WLei*. SARS-CoV-2 isolates are named following the Pangolin lineage classification. Alpha (B.1.1.7), Gamma (P.1), Delta (B.1.617.2). In the Boxplots, the middle lines represent the median, while the dots inside the boxes represent average CAI values.

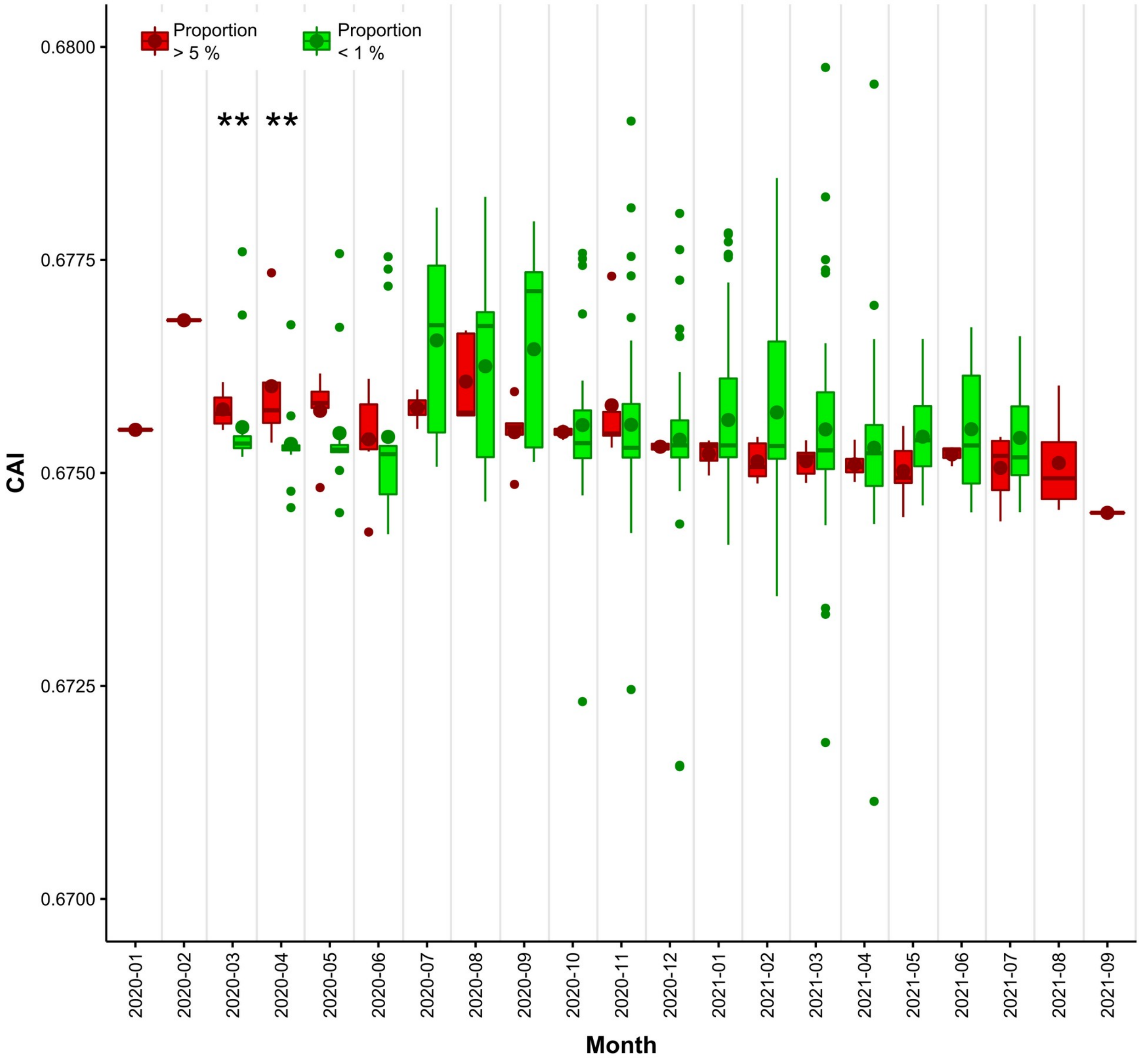


**Figure S15. Evolution of CAI values for the most and less abundant SARS-CoV-2 variants in the California dataset.** Green boxes represent variants with less than 1 percent of relative abundance, while red boxes represent variants with more than 5 percent of relative abundance. In the Boxplots, the middle lines represent the median, while the dots inside the boxes represent average CAI values. Asterisks represent significant differences. P values were calculated using Wilcoxon rank sum test (** < 0.01).

**[PDF File]**

**Figure S16. Evolution of CAI for SARS-CoV-2 ORFs over time for the California dataset.** CAI was calculated for each SARS-CoV-2 gene using *WLei* and averaged by month. A) CAI values calculated for each ORF using the complete dataset. Black line, Evolution of average CAI values over time. B) Left, CAI values for each ORF averaged by month and the selected variants of interest. Right, percentage of coding sequences belonging to each variant.

### Tables

**Table S1. NCBI accession numbers for the sequences used in this work.** Sheet 1- Sequences: Accession numbers for the Beta-CoVs reference sequences, and for SARS-CoV-2 genomes of the variants Alpha, Delta, Gamma and Delta used on this work. Sheet 2 – Time series for CA: Accession number for all the SARS-CoV-2 genomes used on the time series experiments. Sheet 3 – California USA: Accession numbers for the genomes of SARS-CoV-2 isolates from California – USA. Sheet 4 – California USA variants: Relative abundance of SARS-CoV-2 complete genomes from California USA, clustered by Pangolin lineage.

**Table S2. SARS-CoV-2 Preferred Codons and linear dependence of GC3 vs GC12 calculated using the California dataset.** Sheet 1 - SARSCoV2 Preferred Codons. Sheet 2 - Neutrality plots vs Date: GC3 and GC12 were calculated for SARS-CoV-2 genome sequences from the California dataset. The linear dependence of GC12 and GC3 was evaluated for SARS-CoV-2 isolates clustered by month. Sheet 3 – Omicron ENC and CAI: Average ENC and CAI values for Omicron variant, and for its ORFs.
