## Supplementary figure S12. for "Analysis of SARS-CoV-2 synonymous codon usage evolution throughout the COVID-19 pandemic"

**Figure S12. Evolution of CAI values calculated for concatenated SARS-CoV-2 genes over time.**

**A.** *W* calculated as the average of *wi* for every highly expressed gene in a specific human tissue (*Wavg*).

### Adrenal Gland

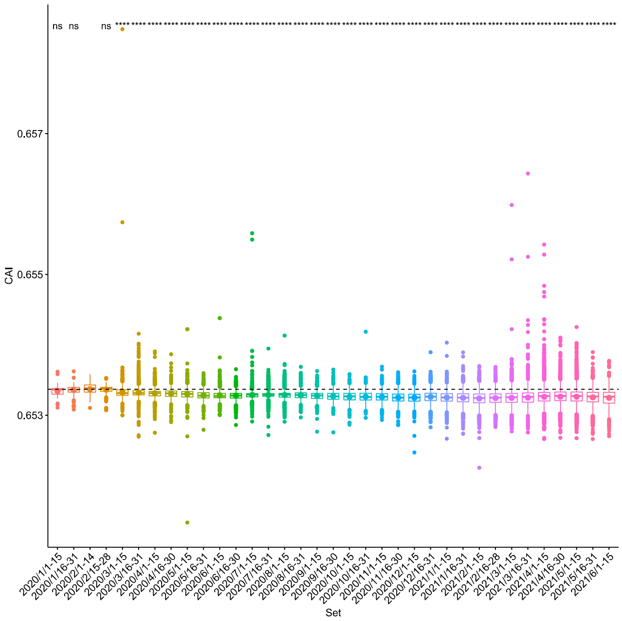


### Thyroid Gland

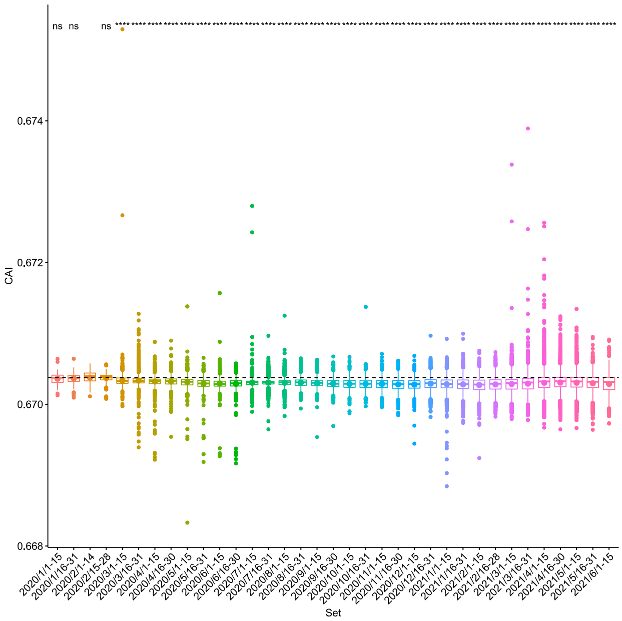


**B.** *W* calculated for concatenated highly expressed genes in a specific human tissue (*Wconcat*).

### Adrenal Gland

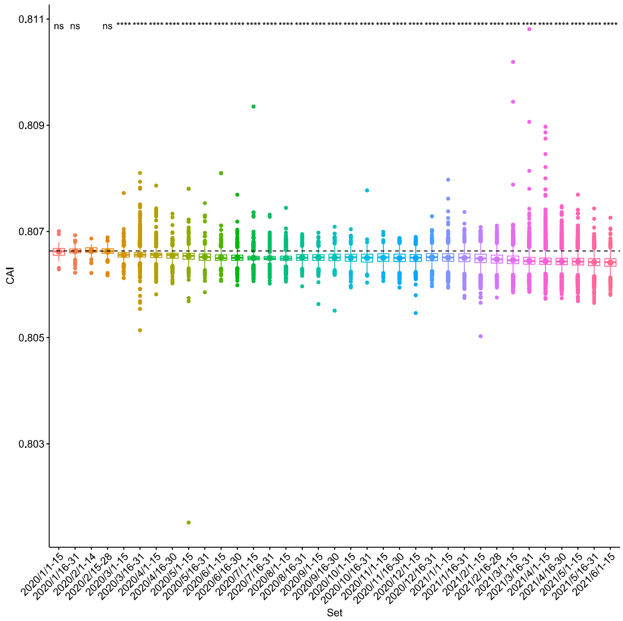


### Thyroid Gland

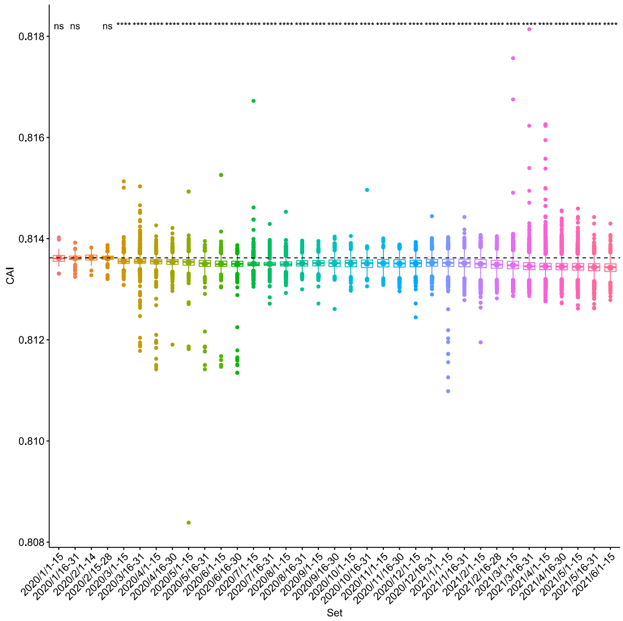

**Figure S12. Evolution of CAI values calculated for concatenated SARS-CoV-2 genes over time.**

**A.** *W* calculated as the average of *wi* for every highly expressed gene in a specific human tissue (*Wavg*).

**B.** *W* calculated for concatenated highly expressed genes in a specific human tissue (*Wconcat*).

### Pituitary Gland

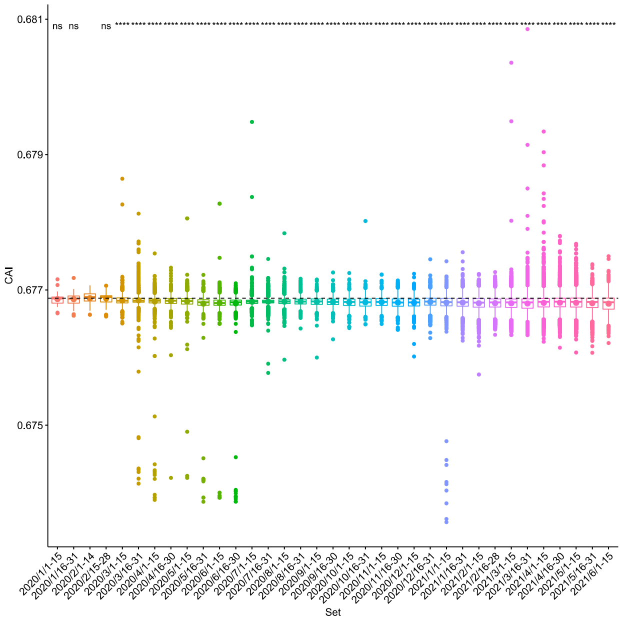


### Pituitary Gland

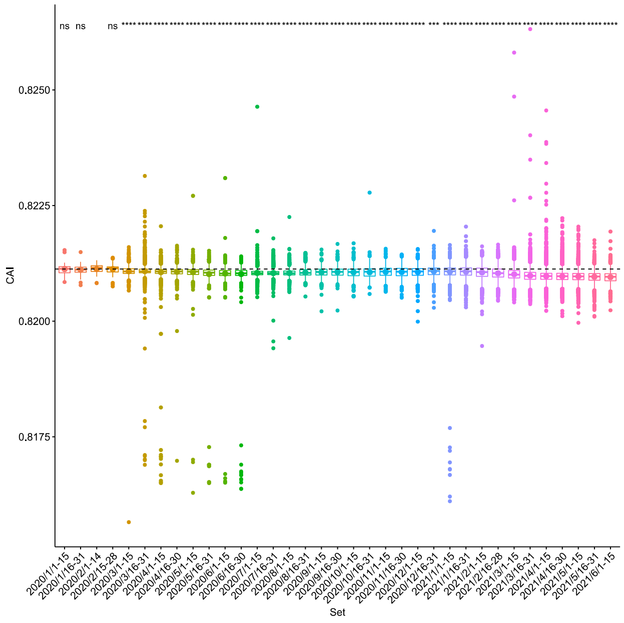


### Kidneys

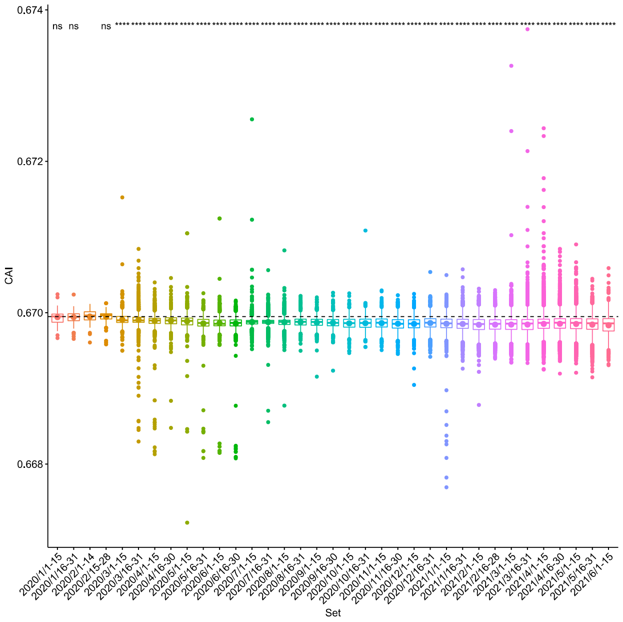


### Kidneys

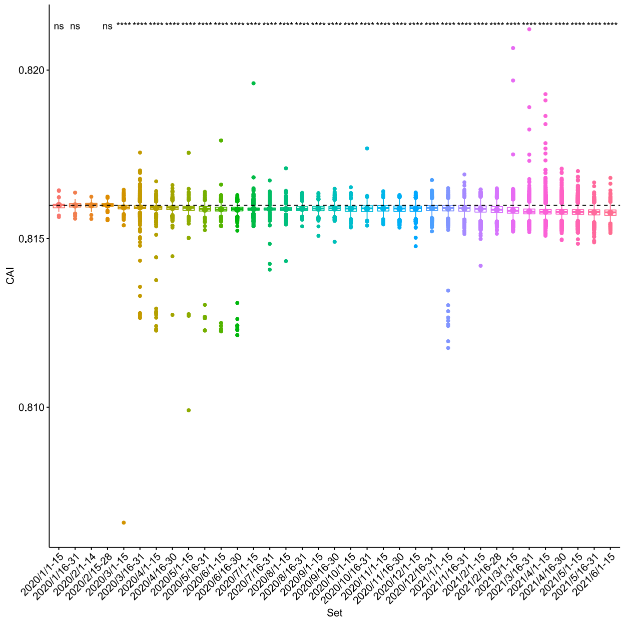

**Figure S12. Evolution of CAI values calculated for concatenated SARS-CoV-2 genes over time.**

**A.** *W* calculated as the average of *wi* for every highly expressed gene in a specific human tissue (*Wavg*).

**B.** *W* calculated for concatenated highly expressed genes in a specific human tissue (*Wconcat*).

### Intestine

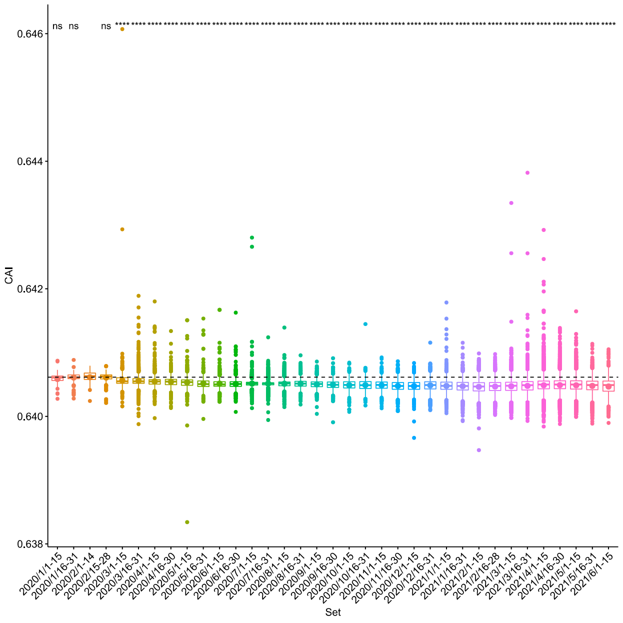


### Intestine

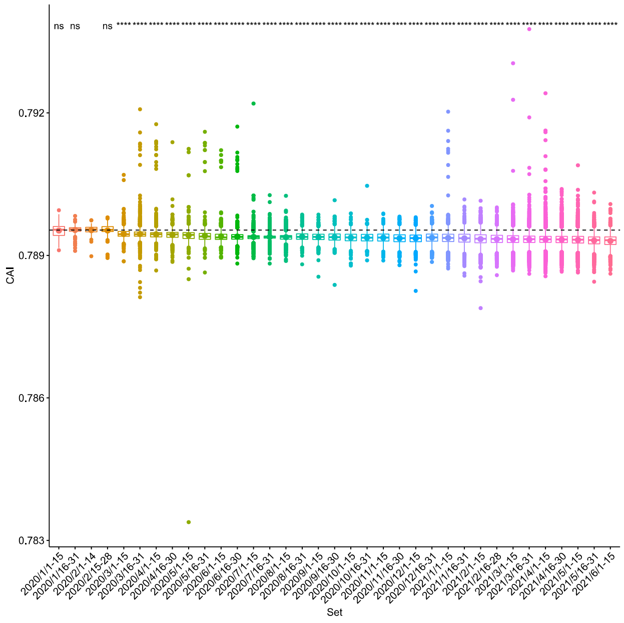


### Heart

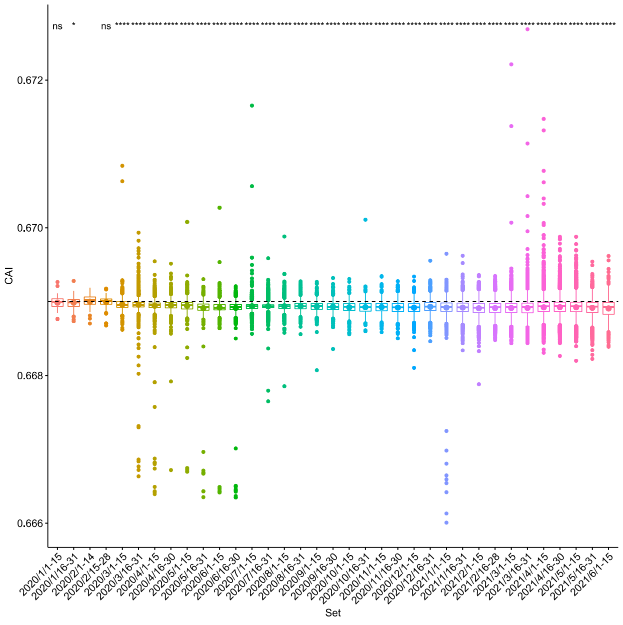


### Heart

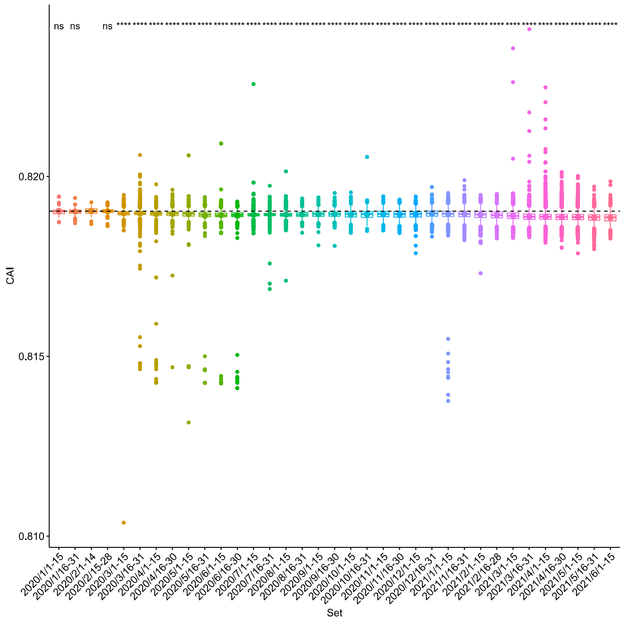

**Figure S12. Evolution of CAI values calculated for concatenated SARS-CoV-2 genes over time.**

**A.** *W* calculated as the average of *wi* for every highly expressed gene in a specific human tissue (*Wavg*).

### Lungs

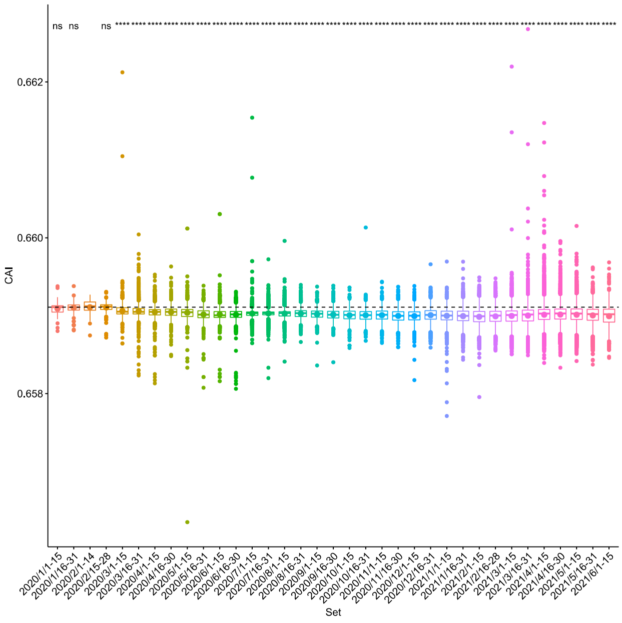


### Brain

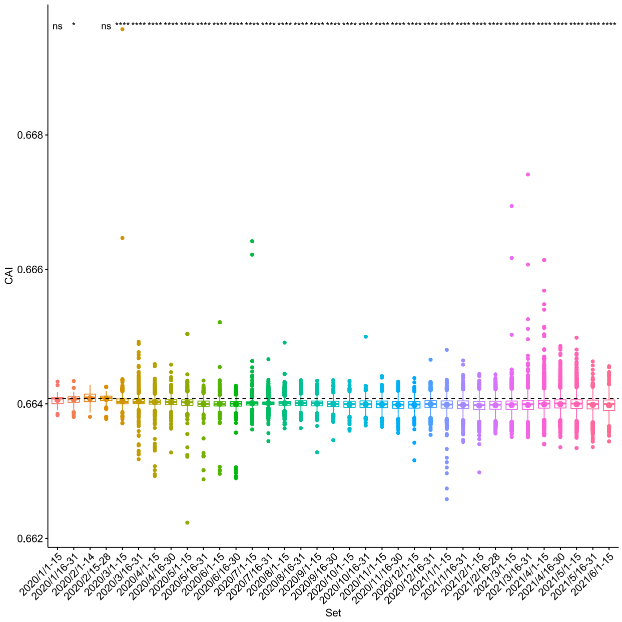


**B.** *W* calculated for concatenated highly expressed genes in a specific human tissue (*Wconcat*).

### Lungs

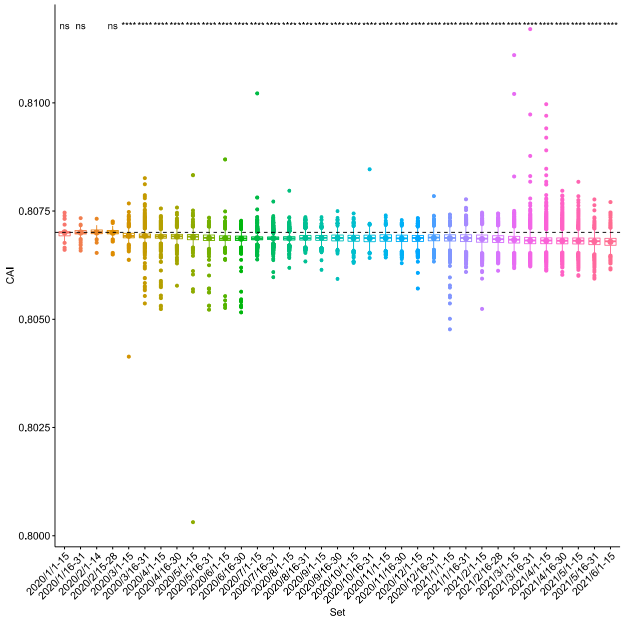


### Brain

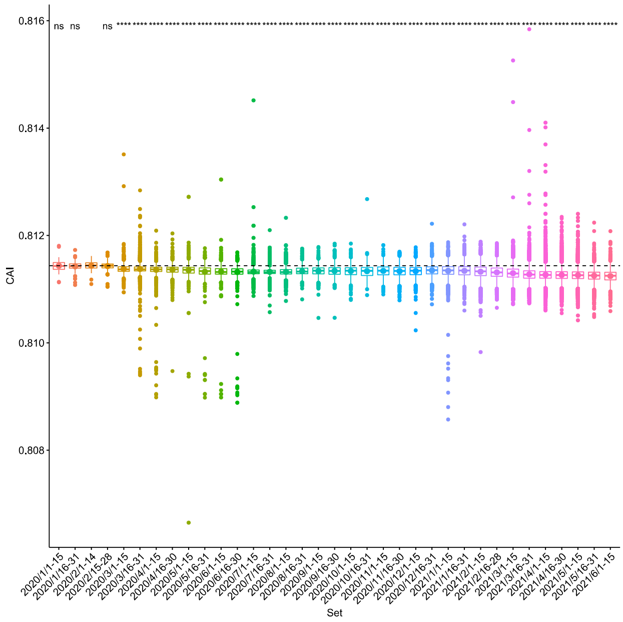

**Figure S12. Evolution of CAI values calculated for concatenated SARS-CoV-2 genes over time.**

**A.** *W* calculated as the average of *wi* for every highly expressed gene in a specific human tissue (*Wavg*).

**B.** *W* calculated for concatenated highly expressed genes in a specific human tissue (*Wconcat*).

### Stomach

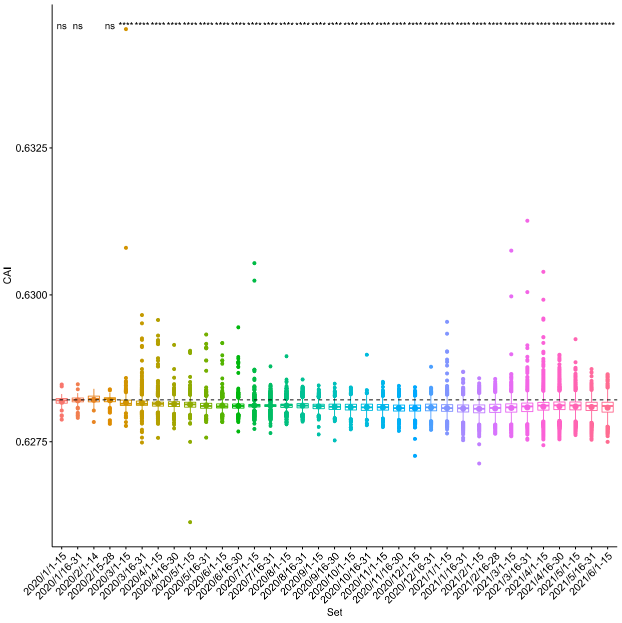


### Stomach

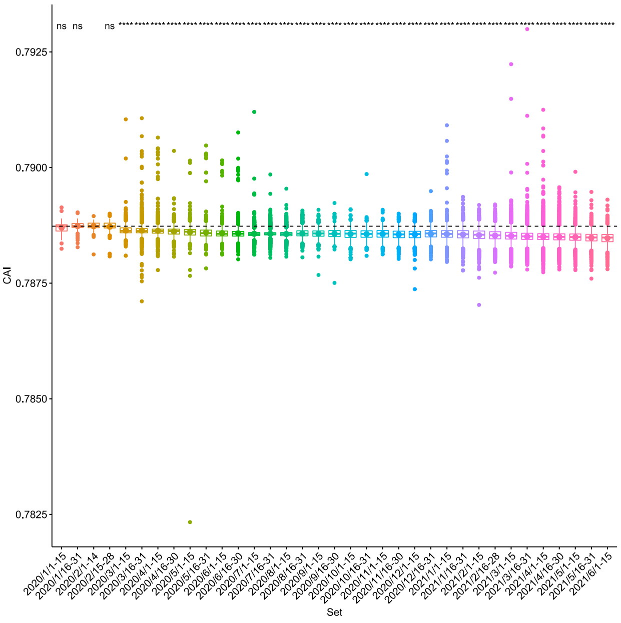


### Spleen

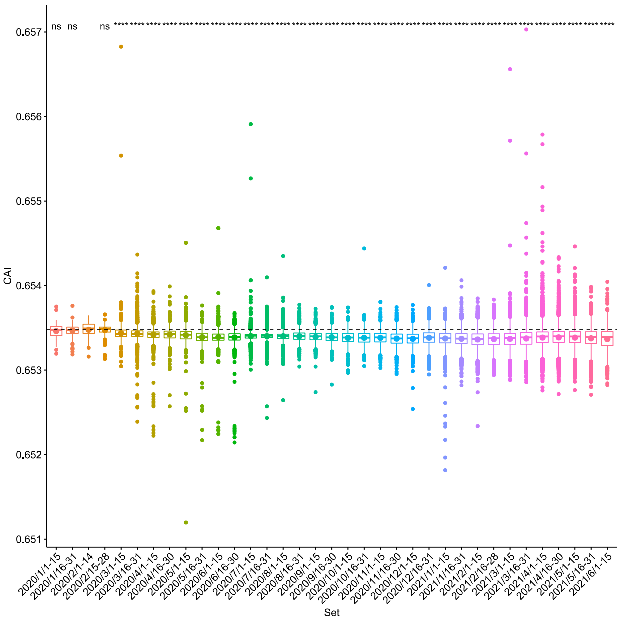


### Spleen

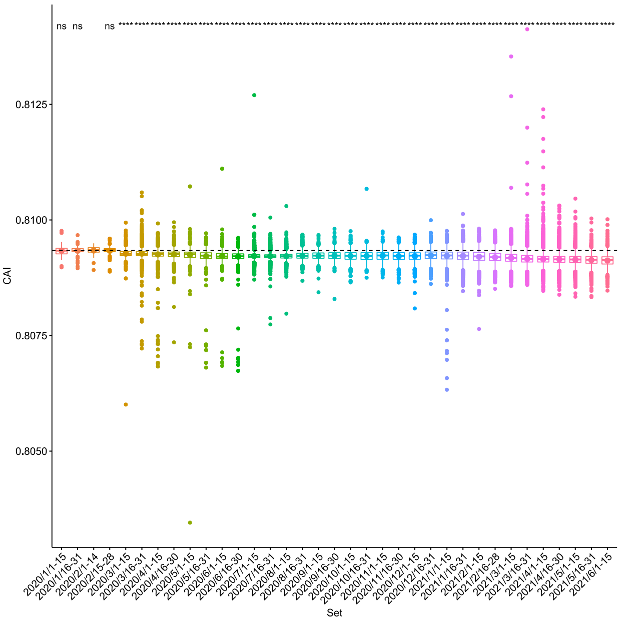

**Figure S12. Evolution of CAI values calculated for concatenated SARS-CoV-2 genes over time.**

**A.** *W* calculated as the average of *wi* for every highly expressed gene in a specific human tissue (*Wavg*).

**B.** *W* calculated for concatenated highly expressed genes in a specific human tissue (*Wconcat*).

**Skin**

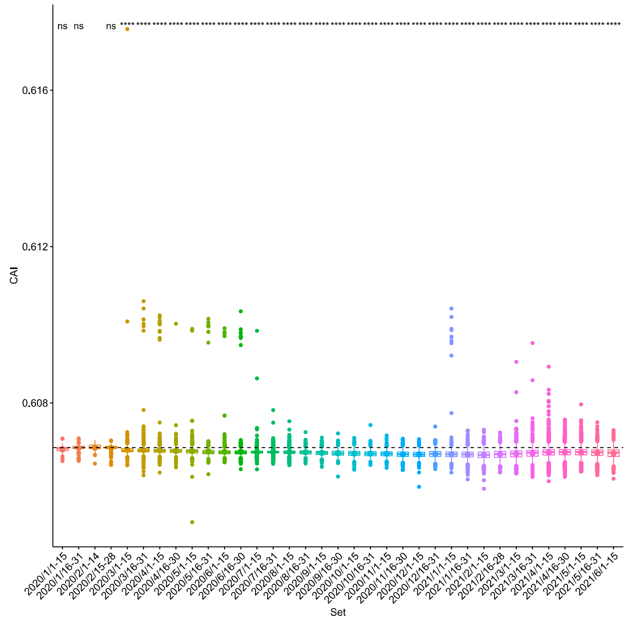


**Skin**

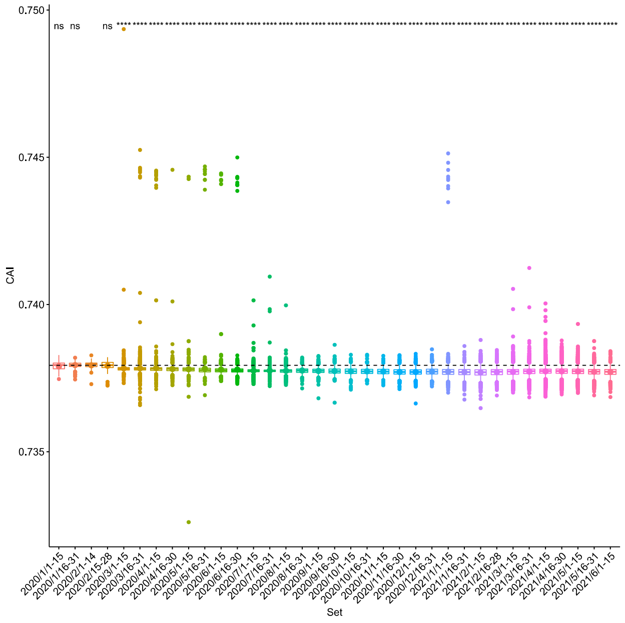


**Eye**

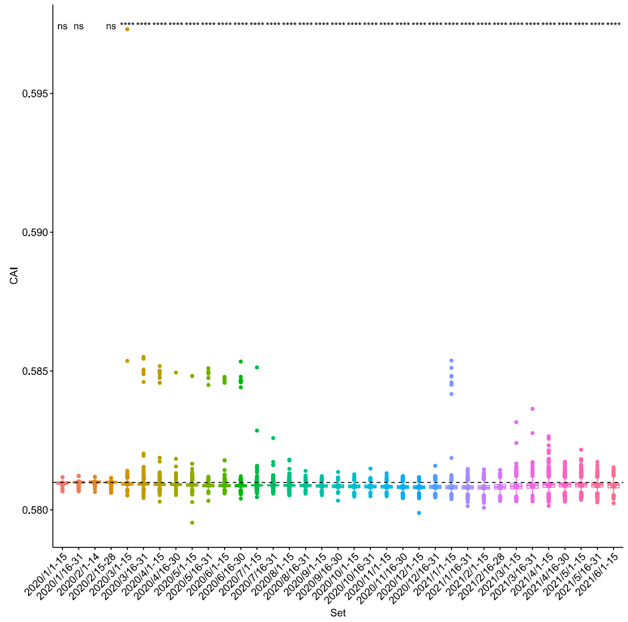


**Eye**

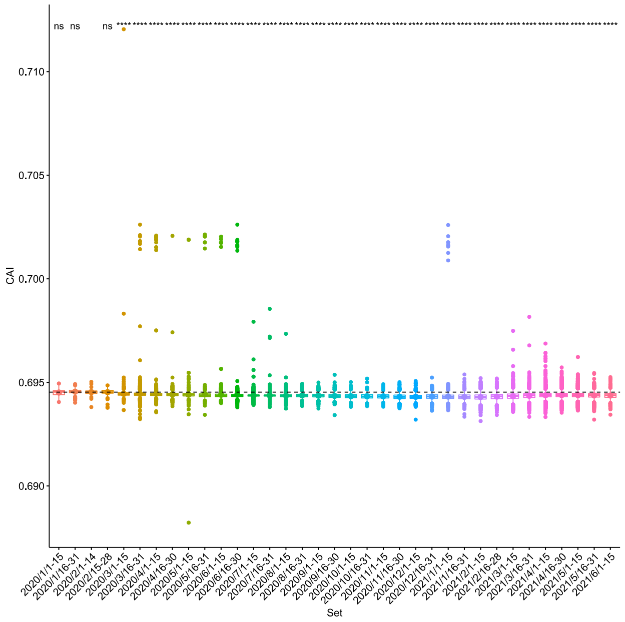

**Figure S12. Evolution of CAI values calculated for concatenated SARS-CoV-2 genes over time.**

**A.** *W* calculated as the average of *wi* for every highly expressed gene in a specific human tissue (*Wavg*).

### Urinary Bladder

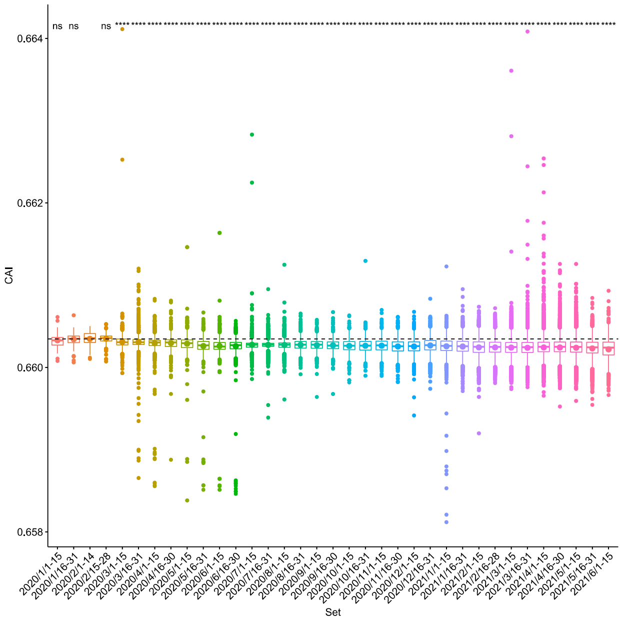


### Granulocytes

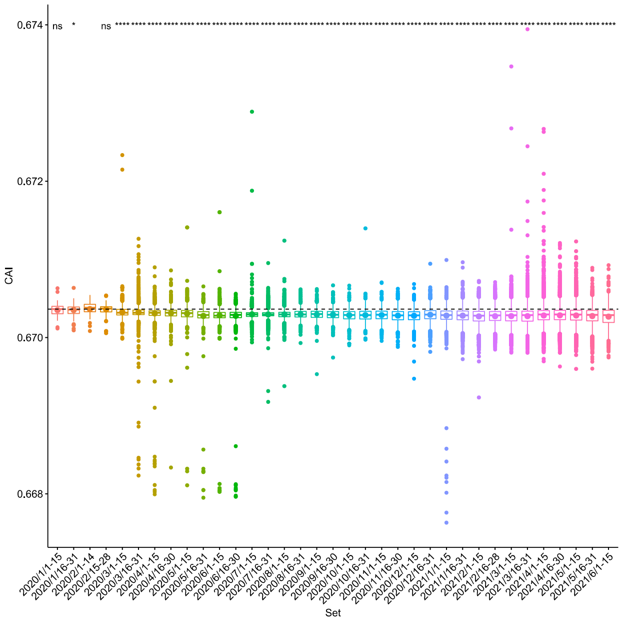


**B.** *W* calculated for concatenated highly expressed genes in a specific human tissue (*Wconcat*).

### Urinary Bladder

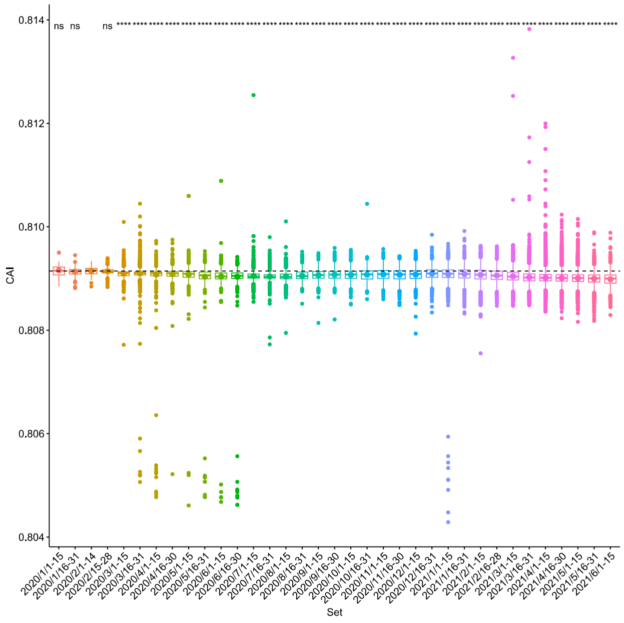


### Granulocytes

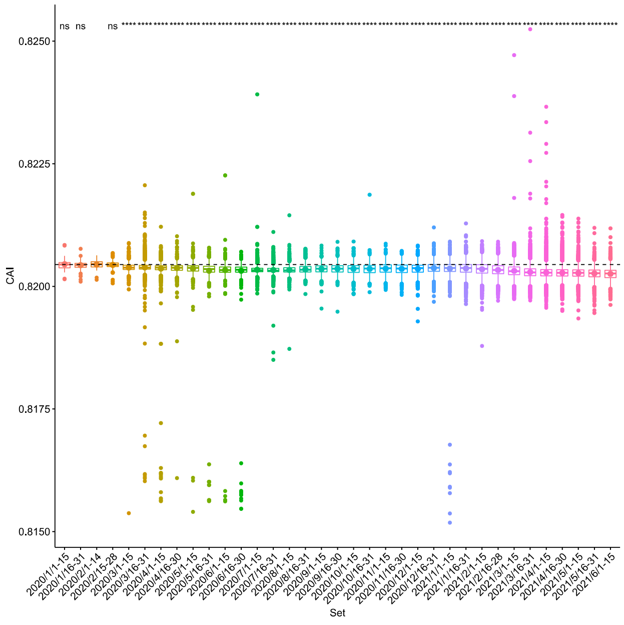

**Figure S12. Evolution of CAI values calculated for concatenated SARS-CoV-2 genes over time.**

**A.** *W* calculated as the average of *wi* for every highly expressed gene in a specific human tissue (*Wavg*).

### Monocytes

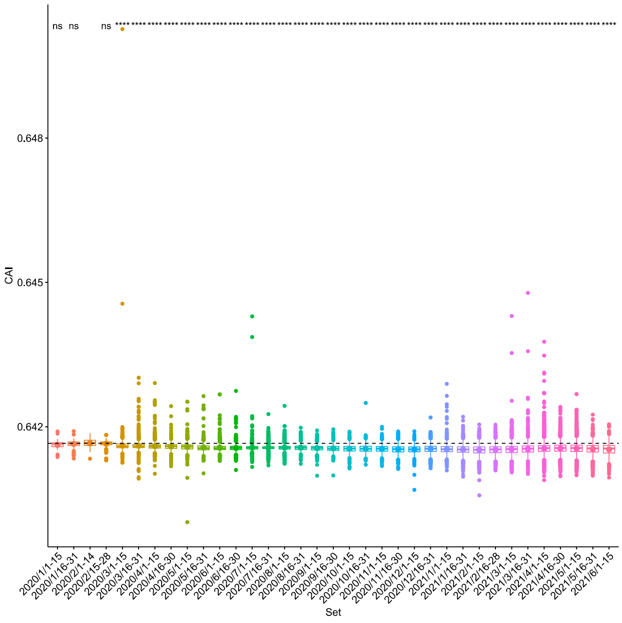


### NK-Cells

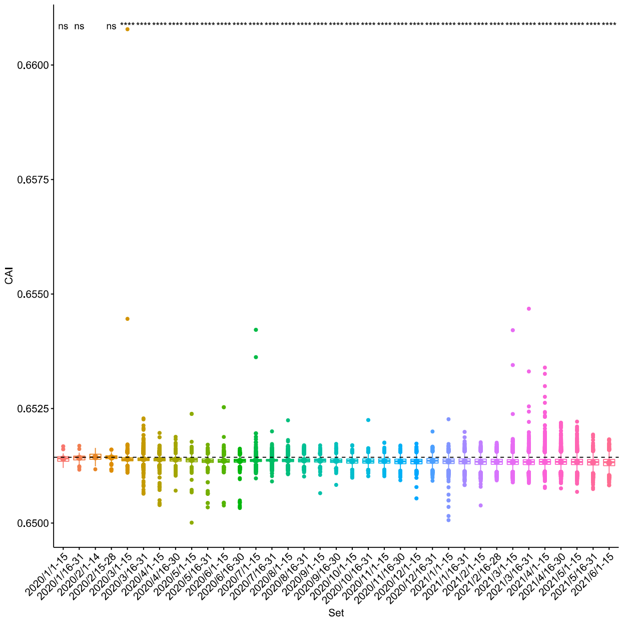


**B.** *W* calculated for concatenated highly expressed genes in a specific human tissue (*Wconcat*).

### Monocytes

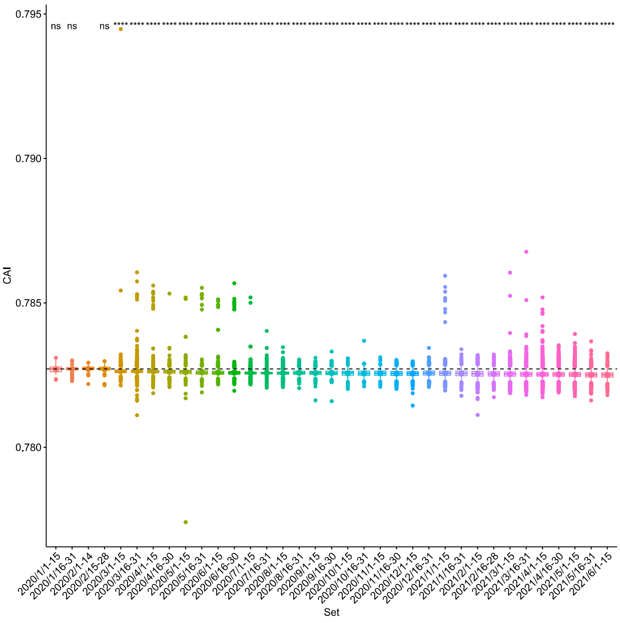


### NK-Cells

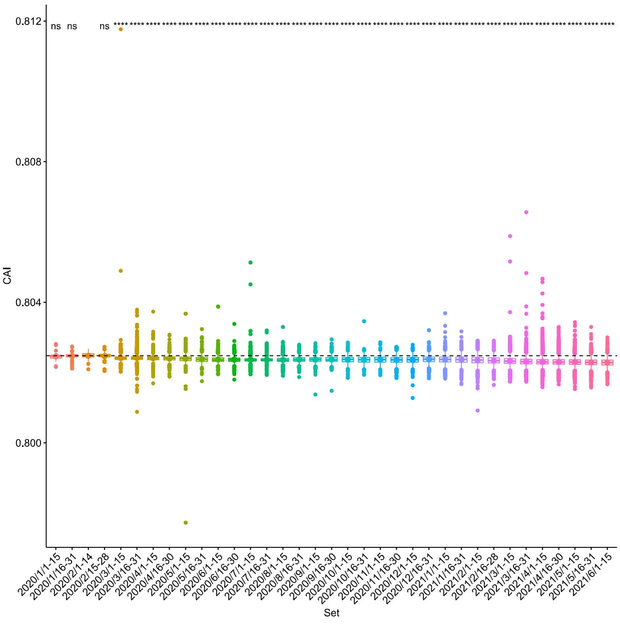

**Figure S12. Evolution of CAI values calculated for concatenated SARS-CoV-2 genes over time.**

**A.** *W* calculated as the average of *wi* for every highly expressed gene in a specific human tissue (*Wavg*).

**B.** *W* calculated for concatenated highly expressed genes in a specific human tissue (*Wconcat*).

### B-Cells

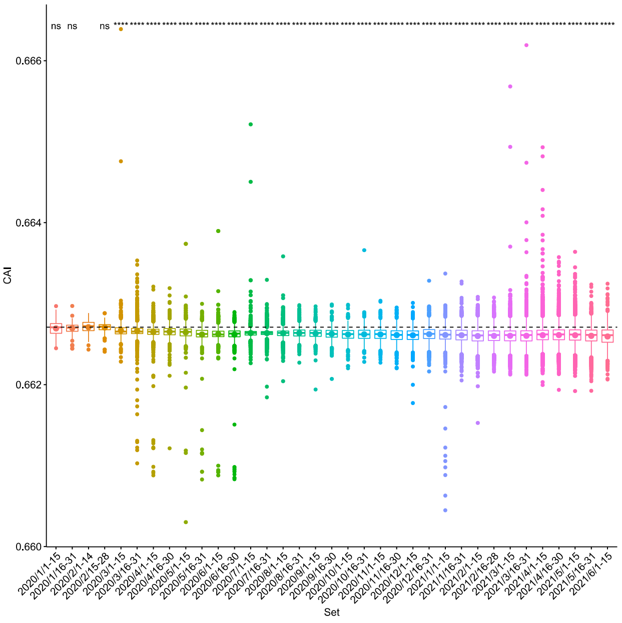


### B-Cells

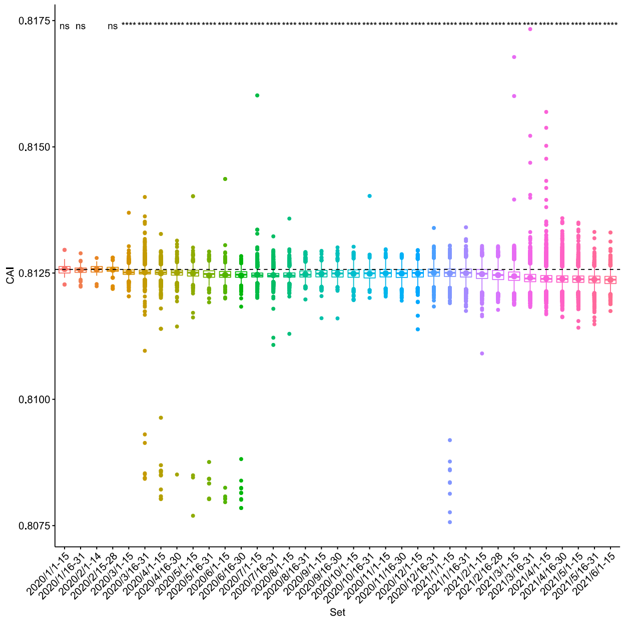


### T-Cells

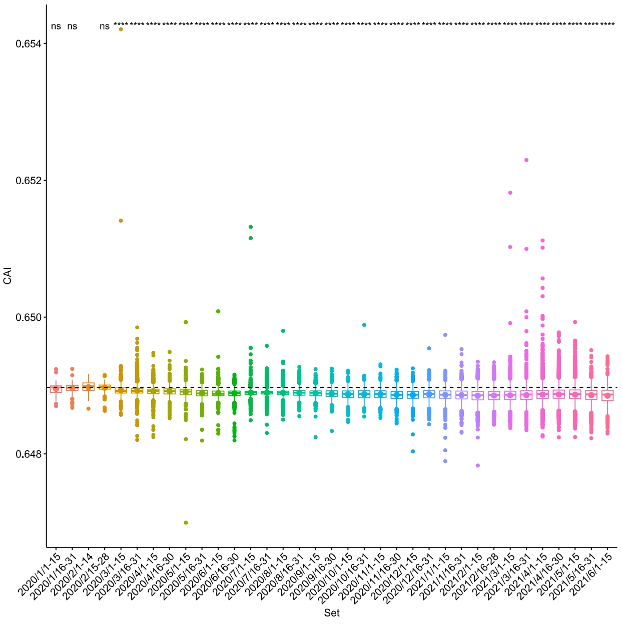


### T-Cells

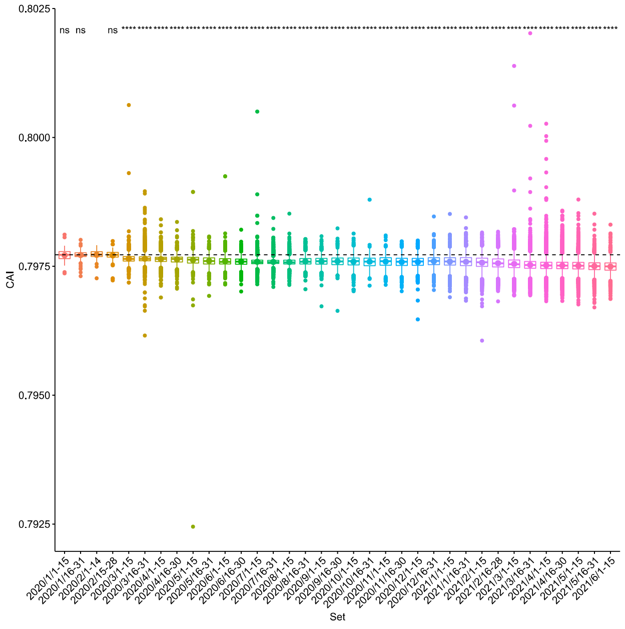

**Figure S12. Evolution of CAI values calculated for concatenated SARS-CoV-2 genes over time.**

**A.** *W* calculated as the average of *wi* for every highly expressed gene in a specific human tissue (*Wavg*).

**B.** *W* calculated for concatenated highly expressed genes in a specific human tissue (*Wconcat*).

### Dendritic Cells

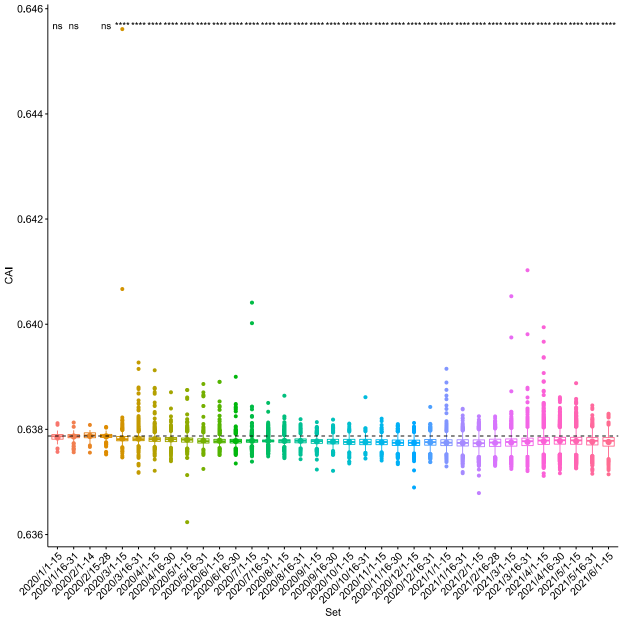


### Dendritic Cells

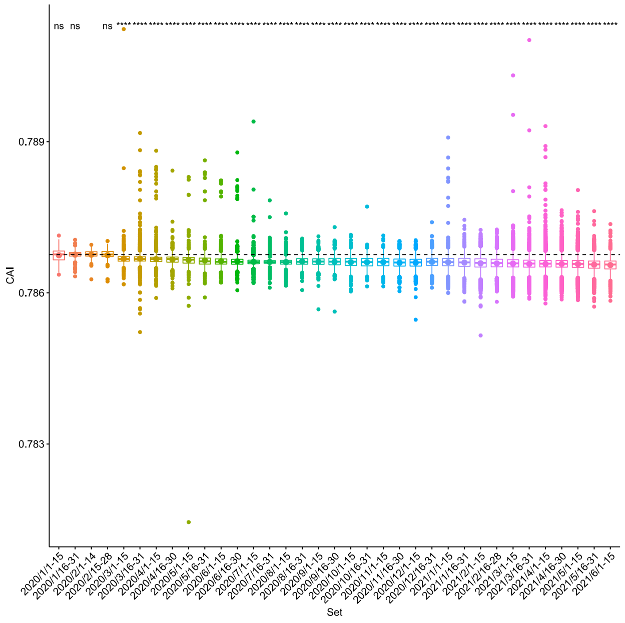


### Lei *et al*.

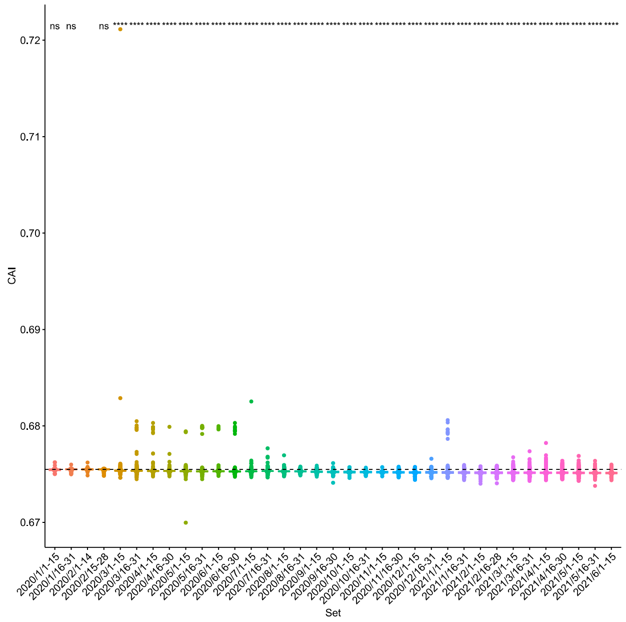
