## Supplementary figure S16. for "Analysis of SARS-CoV-2 synonymous codon usage evolution throughout the COVID-19 pandemic"

**Figure S16.** Evolution of CAI for SARSCoV2 ORFs over time for the California dataset.

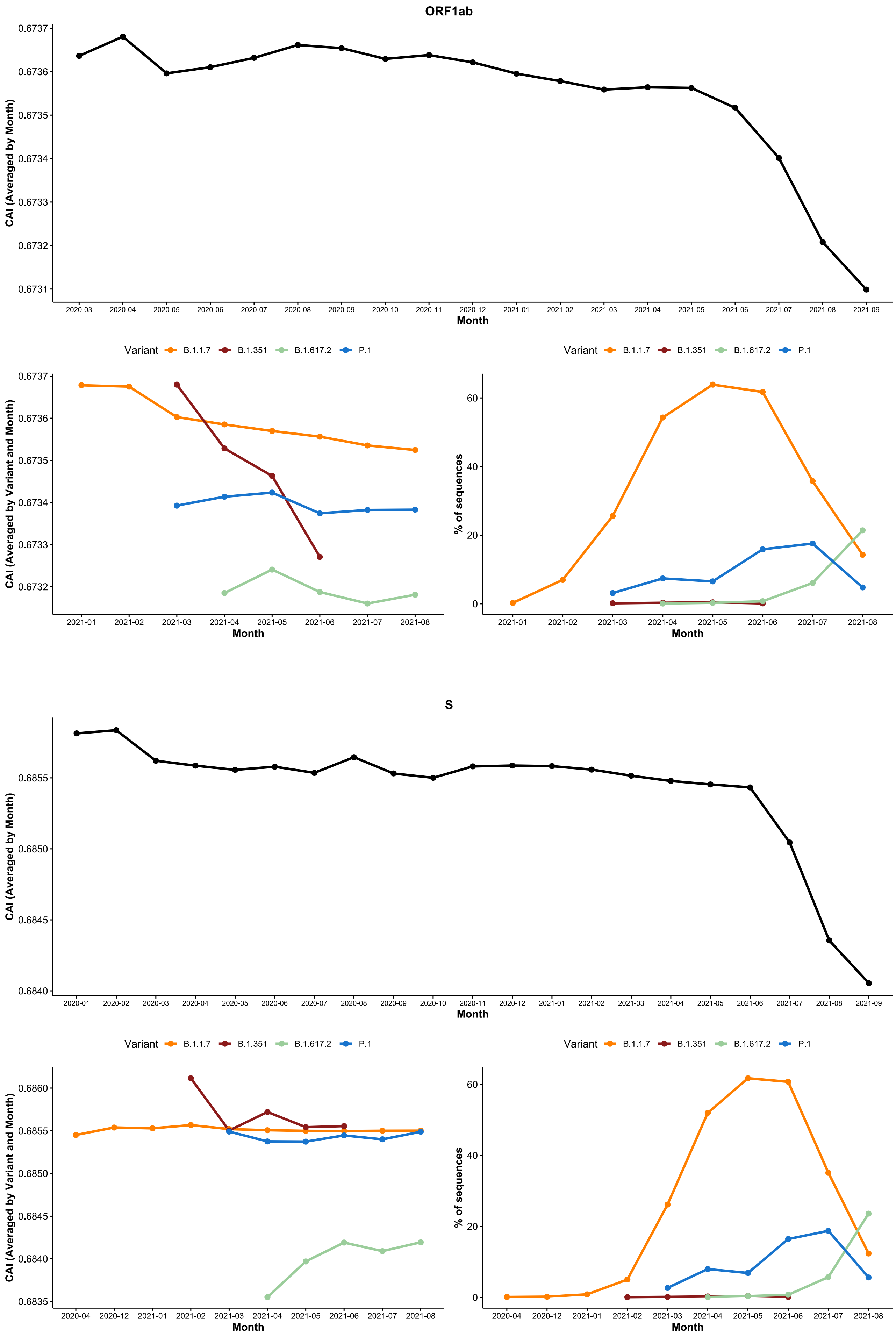

**Figure S16.** Evolution of CAI for SARSCoV2 ORFs over time for the California dataset.

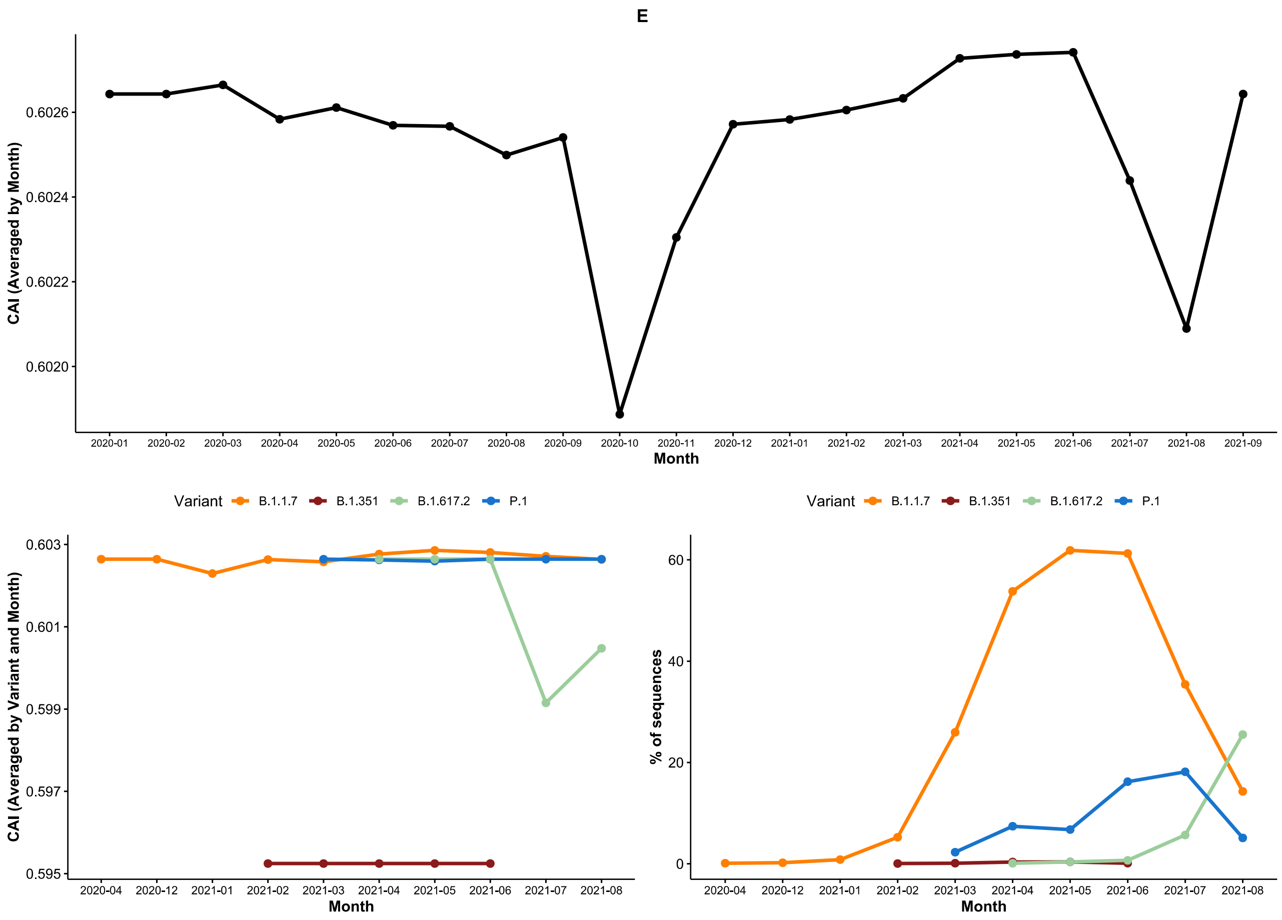


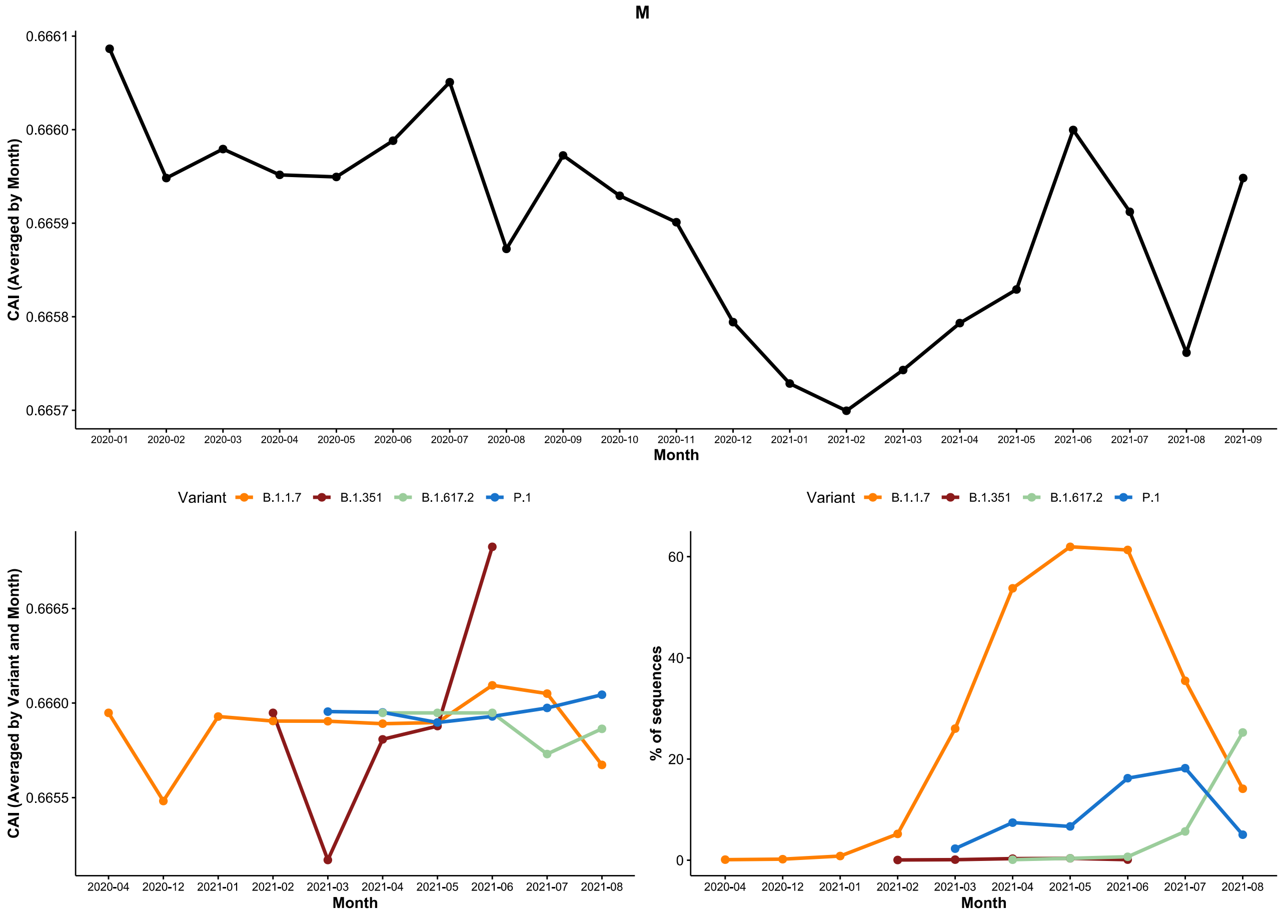

**Figure S16.** Evolution of CAI for SARSCoV2 ORFs over time for the California dataset.

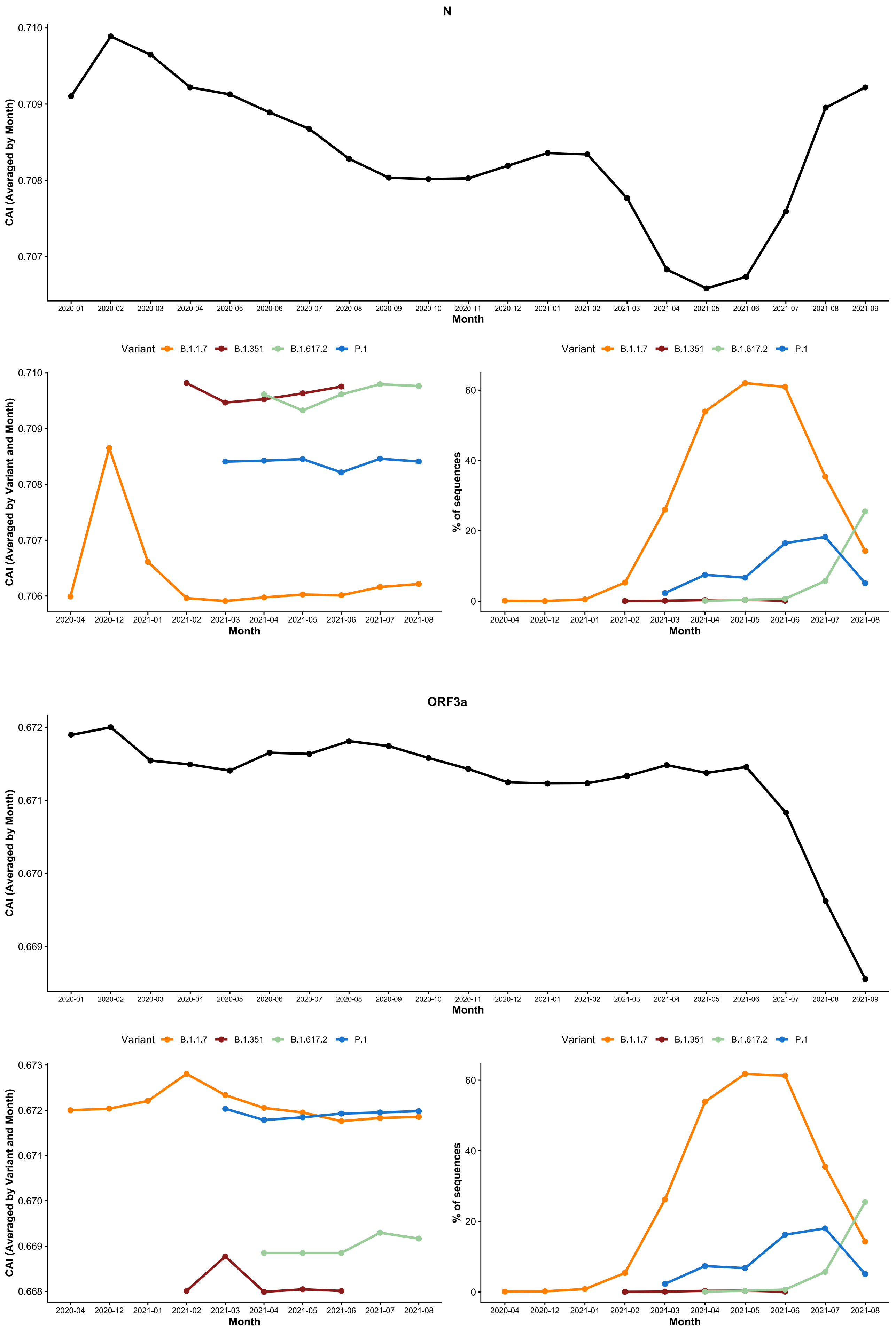

**Figure S16.** Evolution of CAI for SARSCoV2 ORFs over time for the California dataset.

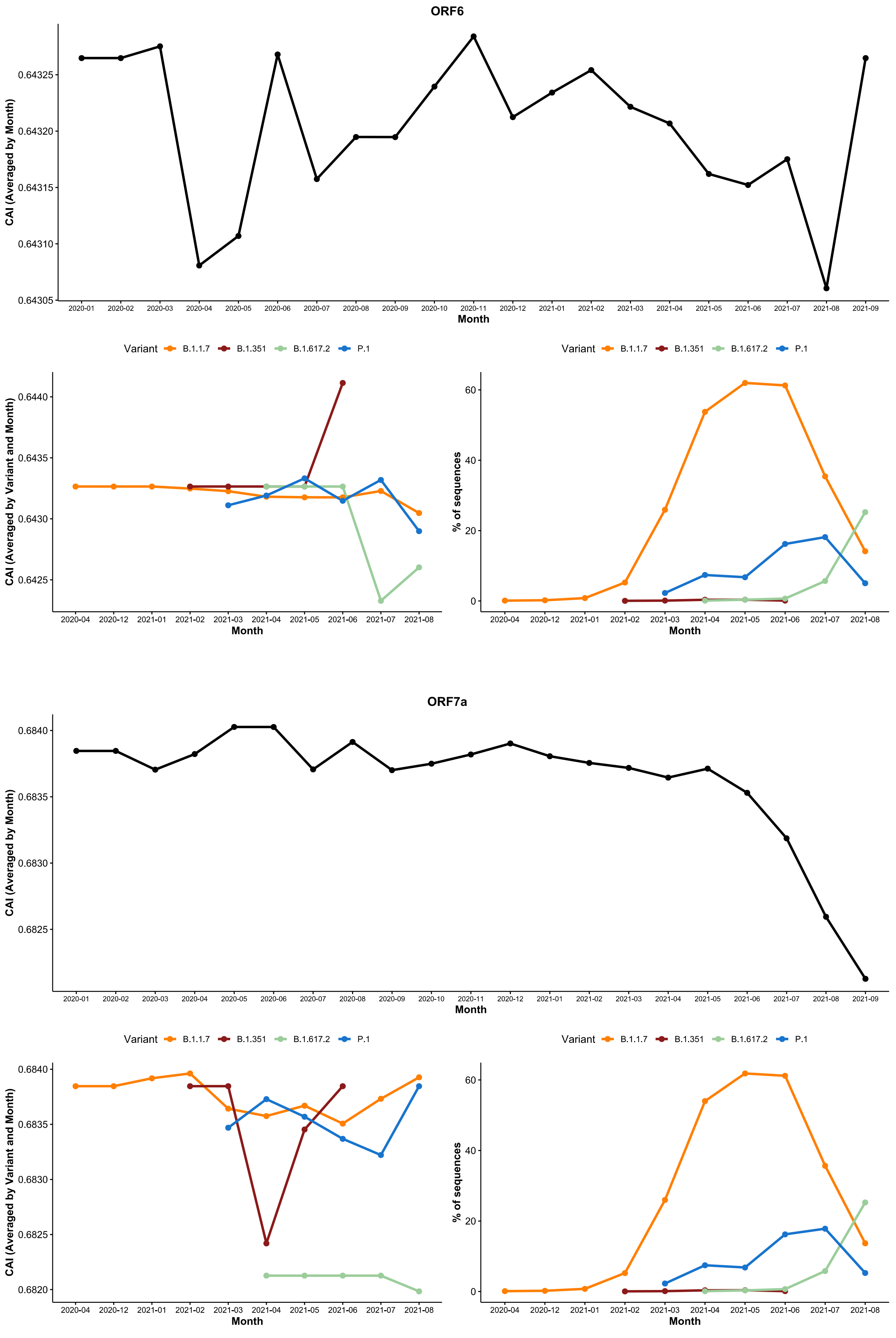

**Figure S16.** Evolution of CAI for SARSCoV2 ORFs over time for the California dataset.

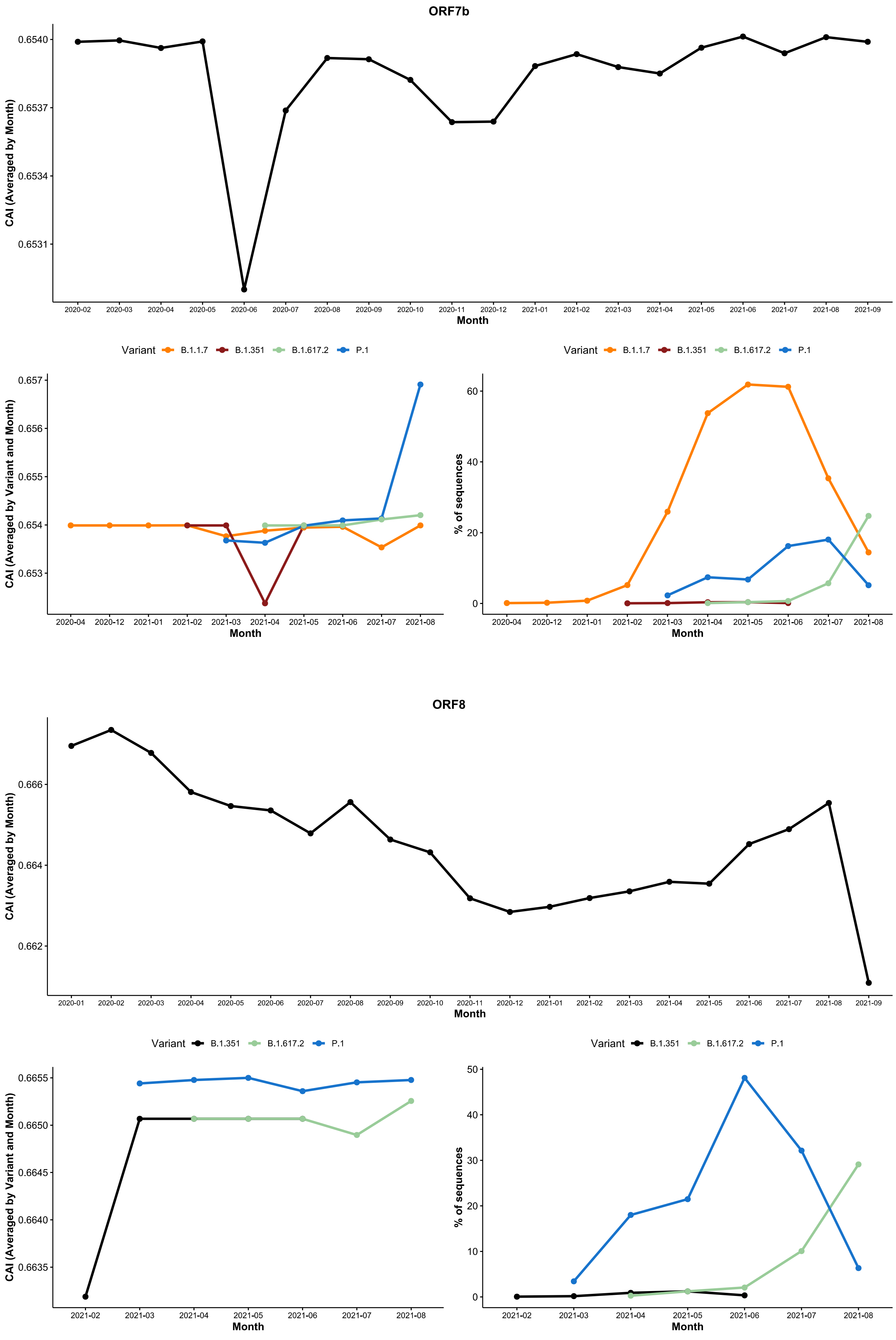

**Figure S16.** Evolution of CAI for SARSCoV2 ORFs over time for the California dataset.

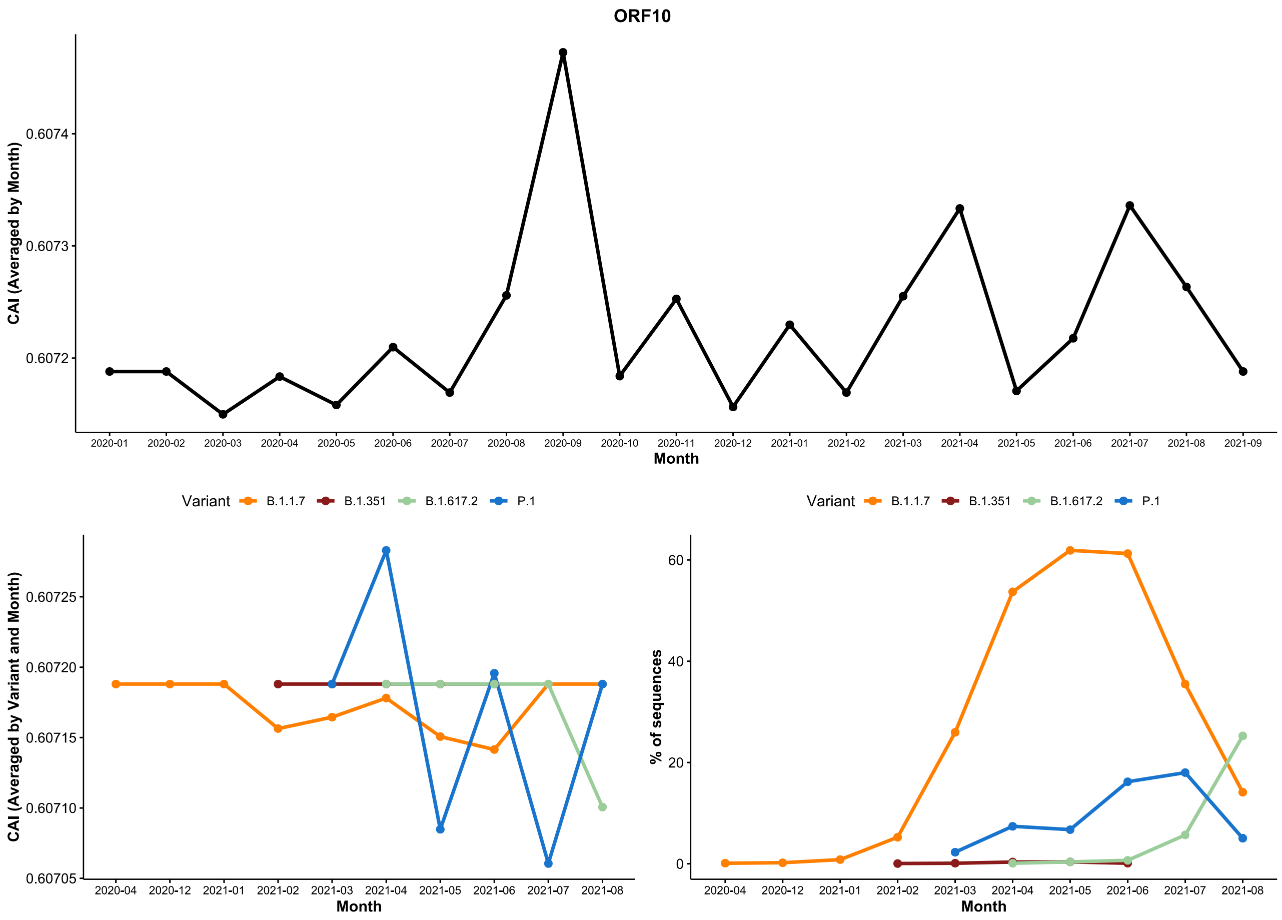
